# The human transcription factor occupancy landscape viewed using high-resolution *in situ* base-conversion strand-specific single-molecule chromatin accessibility mapping

**DOI:** 10.1101/2025.06.27.662080

**Authors:** Georgi K. Marinov, Benjamin R. Doughty, Julia M. Schaepe, Tong Wang, Madison M. Smith, Minyong Chen, Anshul Kundaje, Zhiyi Sun, William J. Greenleaf

## Abstract

Chromatin accessibility profiling is a key tool for mapping the location of *cis*-regulatory elements (cREs) in the genome and tracking chromatin state dynamics during development, in response to various external and internal stimuli, and in disease contexts. Single-molecule footprinting (SMF) methods that rely on the labeling of individual accessible DNA bases have emerged in recent years as a powerful chromatin accessibility mapping approach, as they provide not just an average readout of accessibility over a given genomic position but also the distribution of accessibility states within a population at the level of individual original DNA molecules. However, methylation-based SMF approaches have often been limited either in their resolution or in labeling readout accuracy. We developed a deaminase-based high-resolution strand-specific single molecule footprinting approach that uses highly active sequence context-independent endogenous methylation-insensitive double-strand DNA (dsDNA) deaminases (CseDa01 and LbDa02 C→U/T-ssSMF), which convert accessible cytosines into uracils (which are converted to thymine after PCR amplification). We demonstrate the application of the method to mapping single-molecule accessibility states in human cell lines in both a short and a long-read format, and quantifying the occupancy states of individual transcription factors (TFs) as well as TF co-accessibility and strand-specific accessibility patterns.

## Introduction

In the vast majority of eukaryotes, the majority of genomic DNA is wrapped around nucleosomal particles, making it physically inaccessible to transcription factors and transcription initiation machinery. In turn, active *cis*-regulatory regions, such as promoters, enhancers, and insulators, are characterized by an open-chromatin, nucleosome-depleted configuration, and this property often defines the position and status of active cREs along the genome ^1,2^. The physical protection of nucleosome-bound DNA was appreciated very early in the development of the molecular biology toolkit, demonstrated by the use of nucleases such as DNase I to identify open chromatin at promoter and enhancer elements around individual genes ^3–5^. Later technological advances enabled the coupling of DNAse I-based chromatin accessibility profiling to microarray ^6,7^ and high-throughput sequencing ^8,9^ readouts, eventually allowing truly genome-wide open chromatin profiling. Finally, the Tn5 transposase, in the form of ATAC-seq ^10^ (**A**ssay for **T**ransposase-**A**ccessible **C**hromatin using **seq**uencing), became the tool of choice for cleavage-based chromatin accessibility mapping due to its high sensitivity and ease of application. Both DNase-seq and ATAC-seq allow not just for the mapping of open chromatin regions, but also for the footprinting of occupancy by individual TFs, at least in some cases ^9–11^.

While cleavage-based methods efficiently identify candidate cREs, little information about the long-range physical organization of and co-accessibility patterns within individual DNA molecules can be derived from these short-read data, and because they are enrichment-based, they also do not allow for the measurement of absolute occupancy/protection levels. Single-molecule footprinting is an orthogonal method for measuring chromatin accessibility along single DNA molecules that addresses these shortcomings. Initial SMF-like methods used DNA methyltransferases (MTases) such as the Dam DNA MTase ^12,13^ (which generates m^6^A in GATC contexts), the CpG context-specific M.SssI 5-methylcytosine (5mC) MTase ^13,14^ (applicable in yeast, where there is no endogenous methylation), and the GpC context-specific M.CvPI MTase ^15,16^. Subsequently, M.CvPI GpC-context 5mC labeling was coupled to bisulfite conversion and high-throughput sequencing readout as NOMe-seq ^17,18^. While powerful, the resolution of these methods is limited by the sparseness of the sequence contexts targeted for modification, as CpG or GpC dinucleotides are found on average only once more than every 20 bp in the genome, which is considerably longer than the length of the recognition sites of most eukaryotic transcription factors. To improve this resolution, a combination of both 5mC MTases was used for higher-resolution SMF in organisms lacking endogenous methylation, such as *Drosophila melanogaster* ^19^. More recently, CpC context-specific 5mC MTases such as M.CviQIX ^20^ have also been described; however, equalizing the activities of three different enzymes is challenging. Another drawback of traditional NOMe-seq is the use of bisulfite conversion, which is highly destructive to DNA and thus limits read lengths; this limitation is to an extent addressed by non-destructive enzymatic conversion methods (e.g. EM-seq ^21,22^) enabling high-quality SMF readouts ^23,24^.

Higher resolution and longer read lengths have more recently been achieved by combining m^6^A methylation and long-read single-molecule sequencing, first using the m^6^A MTase EcoGII ^25^ coupled with long-read nanopore sequencing for long-range single-molecule profiling (SMAC-seq ^26^), later followed by Hia5 ^27^ coupled with PacBio sequencing (Fiber-seq ^28^). However, these approaches too suffer from shortcomings, including the inability to amplify the m^6^A-labeled DNA, the relative inefficiency of the m^6^A MTases (which only label 50-70% of accessible adenines), and the high error rates of modification detection using long-read single-molecule sequencing platforms.

The ideal labeling reagent would exhibit near-100% activity for accessible substrates, allow amplification, exhibit no sequence preference, and be unaffected by the presence of endogenous methylation. The use of DNA deaminases has substantial potential benefits over MTases ^26^, as these enzymes can convert accessible bases *in situ* enabling both subsequent amplification and direct readouts in a variety of formats as well as their portability to single-cell applications, and initial implementation of this deaminase-based SMF approach ^41^ demonstrated a number of these benefits. Extending this methodological toolkit, we describe the use of the recently-identified deaminases CseDa01 and LbDa02 dsDNA cytosine deaminases as a tool for high-resolution chromatin accessibility profiling, in several different readout formats, as CseDa01/LbDa02-C→T-SMF. We apply a strand-specific version of the method (C→T-ssSMF) to map and quantify TF footprinting and co-accessibility patterns in human lymphoblastoid cells.

## Results

### High-resolution chromatin accessibility profiling using the non-specific CseDa01 dsDNA deaminase

The use of cytosine deaminases for chromatin accessibility profiling ^26^ has been hampered by the lack of available enzymes that act highly efficiently on dsDNA ^21,22,29^. A large set of deaminases were recently characterized for their activity on dsDNA ^30^, revealing two – CseDa01 and LdDa02 – with ideal properties for SMF, i.e. very high activity on dsDNA, no sequence specificity, and complete insensitivity to 5mC methylation. The latter is important because while CpG positions can be filtered out computationally (as they are endogenously methylated in many eukaryotes), that results in decrease of the effective resolution. Furthermore, CpG positions are often concentrated in regulatory elements in mammals, there are transcription factors, such as NRF1 ^31^, that are specifically methylation sensitive, and some eukaryotes also exhibit extensive 5mC methylation in non-CpG contexts ^32,33^.

Previous work (reproduced in Supplementary Figure 1) demonstrated that the CseDa01 and LbDa02 enzymes exhibit the highest activity out of all deaminases on dsDNA (as well as on ssDNA), have no sequence specificity, and exhibit equal activity on both unmodified substrates and common cytosine modifications. For comparison, enzymes such as MsddA show relatively high activity on unmodified C, but very low activity on 5mC, while SsdA and DddA show much lower absolute activity, and exhibit even lower activity on 5mC compared with unmodified C.

We thus set out to develop a high-resolution version of the SMF assay based on the *in situ* conversion of cytosines to uracils using either of these two enzymes (Figure 1a). Such an assay would provide a resolution of ~5 bp for mammalian genomes, which is slightly lower than the theoretical resolution of m^6^A-based SMF (Figure 1b), but with the advantages of allowing for reliable base calling, more efficient modification of accessible positions, and the ability to carry the accessibility tags through PCR amplification.

**Figure 1:**
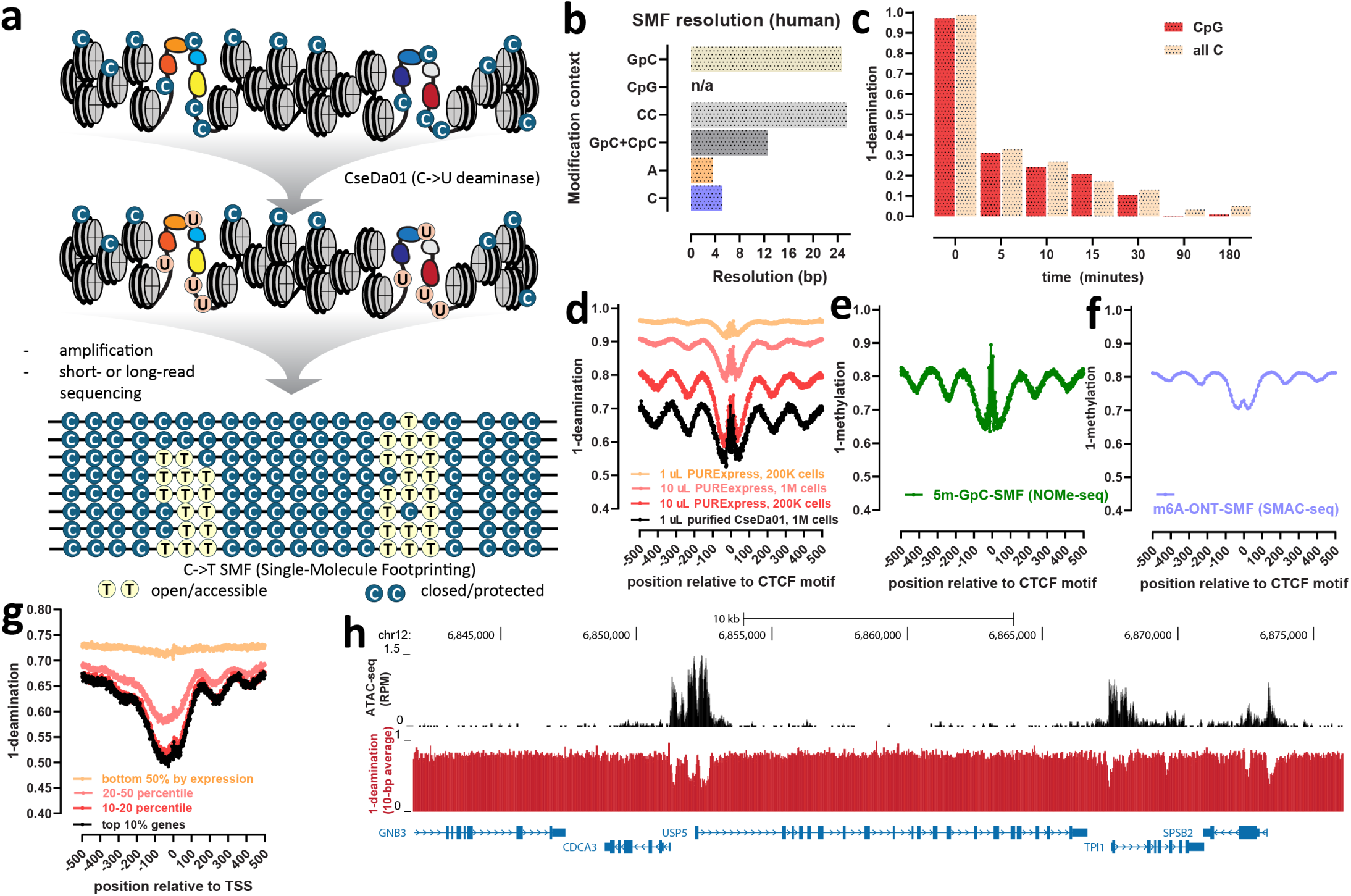
High-resolution chromatin accessibility profiling using the non-specific methylation-insensitive CseDa01 dsDNA deaminase. (a) Outline of the C→T-SMF assay. (b) Average resolution of accessibility profiling using different enzymatic modifications. Shown is the average distance between instances of each sequence context (or combination of) in the human genome. (c) The CseDa01 enzyme efficiently deaminates modified and unmodified cytosines. Human gDNA was isolated and 1 *µ*g of it was incubated with 1 *µ*L of CseDa01 PUREexpress prep for the indicated number of minutes. The average genome-wide deamination levels around all CpG and all C positions in the human genome are shown. (d-g) CseDa01-C→T-SMF accurately captures chromatin accessibility patterns in mammalian cells. (d) Short-read C→T-SMF metaprofiles around CTCF motifs within CTCF ChIP-seq peaks for experiments carried out with PURExpress CseDa01 preps or purified CseDa01 and the indicated amount of enzyme and number of cells (K562 cells, crosslinked). (e) Metaprofiles around CTCF motifs within CTCF ChIP-seq peaks for GpC methylation-based SMF (5mC-GpC-SMF; K562 cells, crosslinked). (f) Metaprofiles around CTCF motifs within CTCF ChIP-seq peaks for m^6^A nanopore-based SMF (m^6^A-ONT-SMF; GM12878 cells, crosslinked). (g) Short-read C→T-SMF metaprofiles around TSSs divided by expression levels (K562 cells, crosslinked). (h) Correspondence between ATAC-seq and short-read C→T-SMF profiling of chromatin accessibility. Shown is an example UCSC Genome Browser snapshot of ATAC-seq accessibility levels, measured in reads per million (RPM), and C→T-SMF accessibility measurements (shown as a smoothed 10-bp average of the inverse of deamination levels).

We used two different preparations of the two enzymes in our work – either in vitro translation (using the PURExpress kit) products (as will be shown further, these enzymes are sufficiently highly active even for unpurified PURExpress preps to be viable tools for probing accessibility) and purified recombinant CseDa01 (see the Methods section for details).

We first measured the activity of CseDa01 on naked human genomic DNA (gDNA), by deaminating 1 *µ*g of gDNA with 1 *µ*L of PURExpress CseDa01 prep over a time course. After generating Tn5 transposition libraries from the deaminated DNA (see the Methods section for details), we observed ~70% conversion within the first five minutes, with complete deamination achieved by 90 minutes. We observed no meaningful difference in conversion levels between CpG positions and non-CpG positions (Figure 1c), confirming CseDa01’s methylation insensitivity.

Next, we aimed to identify the amount of enzyme required to accurately map chromatin accessibility *in vivo*. Because recent studies by us and others ^24^ have shown that the dissociation kinetics of TFs can be very fast relative to the MTase treatment in traditional NOMe-seq/SMF, making it impossible to see footprints for such TFs, we carried out most of our experiments on formaldehyde-crosslinked cells (which we refer to as XL-SMF). We used Tn5 transposition to generate sequencing libraries for these tests (see

Methods for details). As we observed slightly more *in vivo* activity with CseDa01 than with LbDa02 (Supplementary Figure 2), and due to the reported much higher activity of CseDa01 for more exotic modifications such as glucosylated 5hmC (5gmC), even if those are rare in the human genome ^30^, we focused on CseDa01 as the enzyme of choice for subsequent experiments. We determined optimal enzyme levels by comparison to GpC-based NOMe-seq datasets using genome-wide metaprofiles around CTCF motifs within ChIP-seq peaks (as these are characterized by very high chromatin accessibility and a stereotypical nucleosome positioning pattern flanking them) and around transcription start sites (TSSs). By comparing C→T-SMF CTCF metaprofiles to NOMe-seq metaprofiles we determined that 1 *µ*L of purified CseDa01 (at ~0.58 ng/*µ*L) provides similar measures of accessibility when deaminating ~1×10^6^ crosslinked human cells, while 10 *µ*L of CseDa01 PURExpress prep produce an equivalent accessibility profile from ~200,000 human cells (Figure 1d). Metaprofiles around CTCF sites for these conditions are comparable to GpC and m^6^A methylation-based accessibility mapping metaprofiles (Figure 1e and Figure 1f), though we note that we consistently observe 5-10% lower baseline protection levels over nucleosome-occupied DNA than is measured using GpC methylation. These metaprofiles also accurately capture the key aspects of CTCF occupancy – a highly accessible region around the CTCF binding site, a protection footprint over it, and strongly positioned nucleosomes nearby. Other basic aspects of chromatin accessibility are also captured – highly expressed genes are more accessible on average over their promoters than genes that are expressed at lower levels (Figure 1g), and average deamination profiles closely match ATAC-seq profiles (Figure 1h and Supplementary Figure 4). We also successfully generated high-quality deamination-based SMF data from native unfixed samples (Supplementary Figure 3). Overall, these results demonstrated that the CseDa01 deaminase can be used to produce high-quality, high-resolution, single molecule chromatin accessibility data.

We also note that the toolkit of molecular probes to map chromatin accessibility has seen a recent proliferation of assays. For example, m^6^A methylation as a tool for accessibility mapping has been published under the names SMAC-seq ^26,36^, Fiber-seq ^28^, SAMOSA ^37^, SAM-seq ^38^, MATAC-seq ^39^ and STAM-seq ^40^; nanopore-based NOMe-seq/SMF under the names NanoNOMe ^34^ and MeSMLR-seq ^35^. While this manuscript was in preparation, several other dsDNA deaminases were used as probes for chromatin accessibility, first described as FOODIE ^41^, and subsequently as ACCESS-ATAC ^42^, TDAC-seq ^43^, and DAF-seq ^44^). To simplify and unify the nomenclature and help guide readers, we subsequently use the following descriptive convention for SMF assays and their variations:

[enzyme*]-[modification/conversion]-[readout*]-[XL*]-SMF

Where * refers to optional information, i.e. short-read sequencing needs not be specified directly (but long-read platform do) and “XL” refers to crosslinked SMF, and the enzyme name need only be mentioned when introducing the assay in full.

Under this convention the general class of deamination-based SMF mapping methods we present is described as C→T-SMF, if the DNA is amplified after deamination, and C→U-SMF if DNA is read out natively using single-molecule sequencing. C→T-XL-SMF implies crosslinking. Thus CseDa01-C→T-XL-SMF is the full name of our method, as shown in Figure 1. Traditional NOMe-seq would be described as M.CvPI-5mC-GpC-SMF (or 5mC-GpC-SMF in short), SMAC-seq as EcoGII-m^6^A-ONT-SMF (or m^6^A-ONT-SMF in short) and Fiber-seq as Hia5-m^6^A-PB-SMF.

### Single-molecule accessibility profiling using short- and long-read C**→**T-SMF

We then turned our attention to examining single-molecule C→T-SMF profiles using different library building protocols and sequencing readouts. We obtained ~300× sequencing coverage for the GM12878 lymphoblastoid cell line by preparing strand-specific libraries that were sequenced on a Ultima Genomics sequencer (see the Methods section for details), which we refer to as CseDa01-C→T-XL-ssSMF (i.e. strand-specific SMF). Figures 2a and 2b show C→T-ssSMF single molecules over representative occupancy sites for the CTCF and NRF1 transcription factors. We observe footprints over the occupancy site both at the level of single molecules and in aggregate profiles for both proteins.

**Figure 2:**
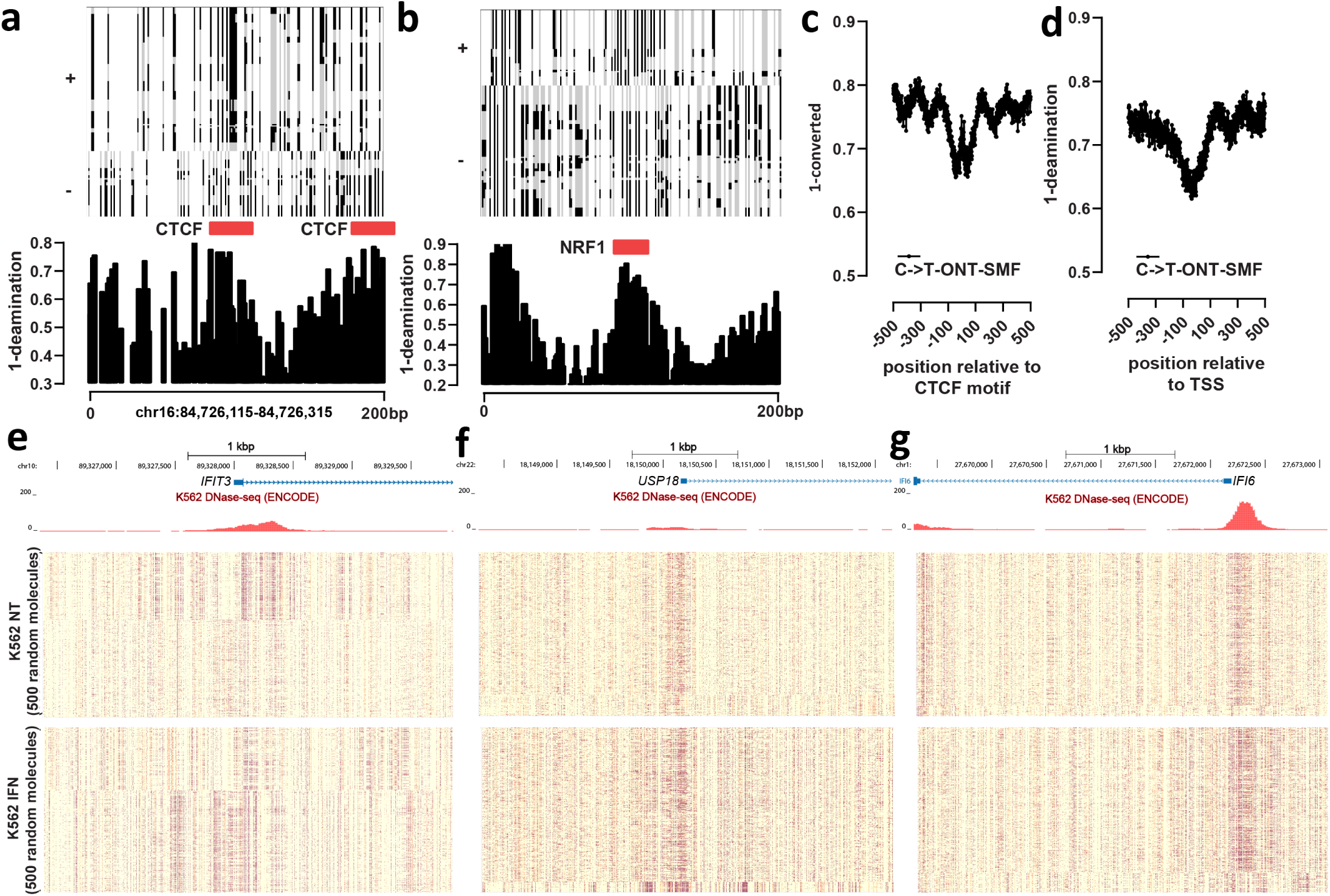
Single-molecule accessibility profiling using short- and long-read CseDa01-C→T-SMF. (a) Representative CseDa01-C→T-SMF footprint patterns around a CTCF binding site. (b) Representative CseDa01-C→T-SMF footprint patterns around an NRF1 binding site. (c) CseDa01-C→T-ONT-SMF metaprofiles around CTCF motifs within CTCF ChIP-seq peaks (K562 cells, crosslinked) (d) CseDa01-C→T-ONT-SMF metaprofiles around TSSs (K562 cells, crosslinked) (e-f) Targeted CseDa01-C→T-ONT-SMF amplicons around the *IFIT3*, *USP18*, and *IFI6* genes. Shown are 500 random single molecules (at 5-bp resolution) for untreated (“NT”) and interferon-treated (“IFN”) K562 cells.

We also generated low-coverage CseDa01-C→T-ONT-SMF for the K562 cell line by carrying out whole-genome amplification of deaminated DNA and sequencing the resulting DNA using nanopore sequencing on a MinIon flow-cell. This readout reproduces metaprofiles over CTCF sites and TSSs that are similar to short-read based C→T-SMF (2c-d) demonstrating the viability of using long reads together with CseDa01 SMF inputs.

Finally, we generated high-coverage targeted C→T-ONT-SMF for the promoters of three genes (*IFIT6*, *IFI3* and *USP18*) in K562 cells with and without treatment with interferon (IFN). We used primers to amplify these amplicons that were carefully designed to avoid potentially deaminated bases (see the Methods section for details). These promoters are partially (*IFIT6* and *USP18*) or fully (*IFI3*) accessible in resting cells, with accessibility levels increasing upon IFN treatment (Figure 2e-g). Overall, these data demonstrate the successful generation of CseDa01-based SMF datasets in a variety of both short- and long-read based formats.

### Transcription factor footprinting patterns in the human genome

We used the high-depth CseDa01-C→T-XL-ssSMF dataset to address several questions – first, we aimed to measure aggregate levels of TF footprinting in the human genome for a collection of TFs; second, we aimed to measure the relationship between this footprinting strength and TF occupancy levels as measured by ChIP-seq; third, we asked if TF foot-printing exhibits differential patterns on the two strands of DNA. To these ends, we compiled sets of TF recognition motifs likely to be highly occupied for 64 TFs for which ChIP-seq data exists from the ENCODE Project Consortium ^1,2^, by identifying binding motifs that are both within ≤100 bp of the ChIP-seq peak, and also the nearest to the peak. These selection criteria aim to exclude potential “bystander” motif matches that might be in the vicinity of the actually occupied site but not bound by the TF. We split these sets of motif matches into quintiles based on ChIP-seq peak strength, and calculated two sets of metaprofiles for each TF – deamination levels over all sites for the forward/plus and reverse/minus strands, and overall deamination levels for each quintile. Detailed results for all TFs are shown in Supplementary Figures 5 to 68. Examining these metaprofiles revealed six general footprinting patterns.

The first pattern is exemplified by the CTCF insulator protein, which is well known for its ability to drive strong nucleosome positioning in its vicinity ^45,46^. This positioning is also observed in our data, as expected; however, we also observe strong strand asymmetry in deamination profiles over the positioned nucleosomes around CTCF (Figure 3a). Such strand assymetry of protection was also been observed in yeast using EcoGII-m^6^A-ONT-SMF ^26^ based on the highly accurate chemical maps of nucleosomes that are available for *Saccharomyces cerevisiae* ^47^. It arises as a result of the physical structure of the nucleosome which preferentially exposes one strand of DNA on one side of the nucleosomal dyad, and the other strand in a mirrored pattern on the other side ^26^. We do not have chemical maps that precisely position CTCF-flanking nucleosomes in GM12878 cells, but despite that potential lack of precision the magnitude of the strand asymmetry observed is larger than that found in yeast (~5% difference compared to ~1% difference). We discuss the likely reasons for these observations in a later section. We do not observe substantial strand asymmetry within the CTCF occupancy site itself (Supplementary Figure 18)

**Figure 3:**
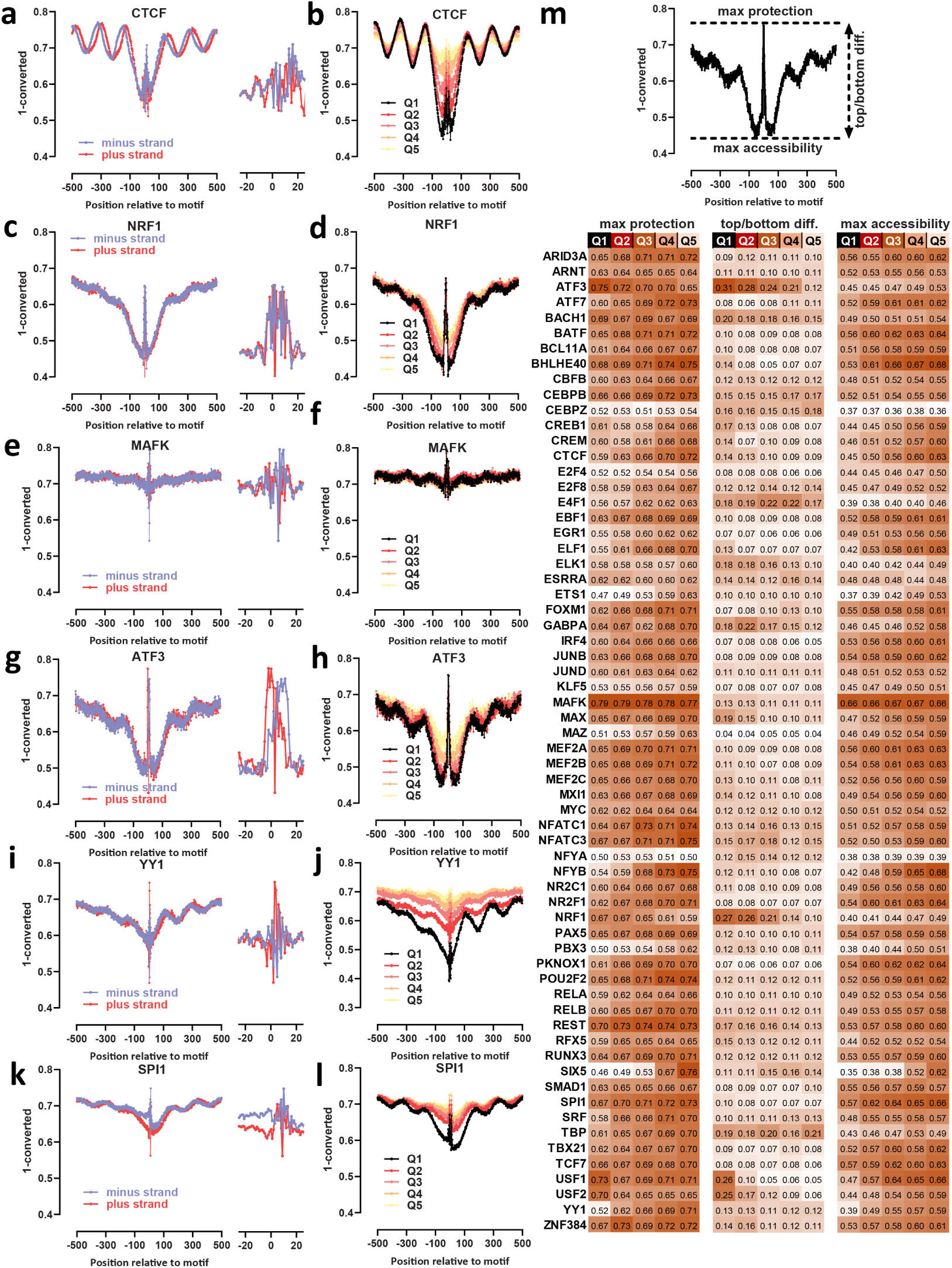
Transcription factor footprinting patterns in the human genome. (a-l) Metaprofiles around the motifs for each TF within ENCODE ChIP-seq peaks and closest to the ChIP-seq peak summit. For each TF, the forward- and reverse-strand C→T-ssSMF metaprofiles are shown on the left, while on the right the profiles are split into quintiles according to the strength of ChIP signal. A zoomed-in forward- and reverse-strand metaprofile is also shown. (m) Quantification of footprinting strength over motifs, split by ChIP peak strength. Three different scores are shown – “maximum protection” shown in the left columns measures the distance from zero to the protection peak over the motif, while “top versus bottom difference” measures the difference between the protection peak over the motif and the lowest protection level around the motif in the metaprofile. The“maximum accessibility” metric shows the depth of the accessibility well and thus average local accessibility associated with each TF.

As an example of our second observed pattern, the NRF1 TF exhibits extremely strong footprinting over the TF recognition motif itself (Figure 3c-d), but weak positioning of nearby nucleosomes and little strand asymmetry over the occupancy site (Supplementary Figure 48).

The MAFK factor is known to have a repressive role ^48,49^ and to occupy sites in closed chromatin, and is an example of our third observed pattern of very weak open chromatin signature with a footprint largely overlaid on top of the baseline protection levels (Figure 3e-f). The NRSF/REST transcription factor is also a well-known repressor, specifically of neuronal genes ^50^, and can strongly position nucleosomes ^51,52^. Its footprinting profiles are a mixture of that of CTCF and MAFK, i.e. inaccessible chromatin and open chromatin regions with strongly positioned nucleosomes nearby (Supplementary Figures 55).

The fourth general type of footprint pattern is exemplified by ATF3, a member of the large family of bZIP TFs. ATF3 exhibits both very strong footprints and extreme strand asymmetry (Figure 3g-h), with as much as 25% difference in absolute accessibility levels between the two strands. This asymmetry is consistent with the structure of bZIP TFs, which typically function as homodimers and/or heterodimers ^53,54^, or oligomers ^55^. This asymmetric protection pattern is consistent with two TFs in a dimer binding DNA on opposite strands in such a manner that the bound strand is well protected while the opposite strand is much more exposed (see also Figure 4b for more details).

**Figure 4:**
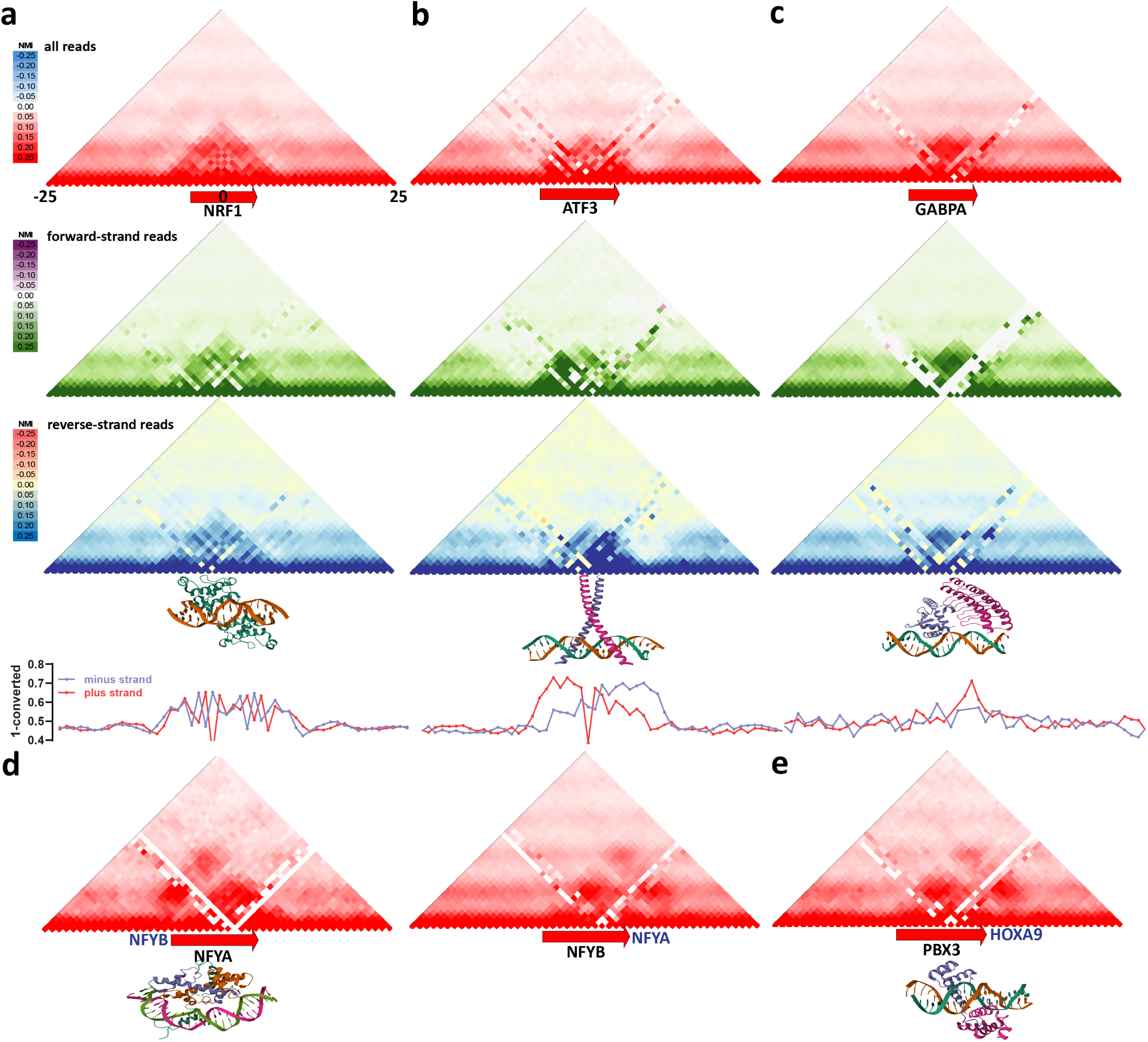
Co-accessibility patterns around transcription factor binding sites. (a) Average NMI values around NRF1 occupancy sites for all reads and forward- and reverse-strand (with respect to the TF motif) reads only. Also shown is the crystal structure of an NRF1 DNA binding domain (DBD) dimer (PDB accession 8K3D), as well as strand-specific footprint profiles around NRF1 occupancy sites. (b) Average NMI values around ATF3 occupancy sites for all reads and forward- and reverse-strand (with respect to the TF motif) reads only. Also shown is the crystal structure of a bZIP dimer (PDB accession 1JNM), as well as strand-specific footprint profiles around ATF3 occupancy sites. (c) Average NMI values around GABPA occupancy sites for all reads and forward- and reverse-strand (with respect to the TF motif) reads only. Also shown is the crystal structure of a GABP dimer (PDB accession 1AWC), as well as strand-specific footprint profiles around GABPA occupancy sites. (d) Average NMI values around NFY occupancy sites. Also shown is the crystal structure of an NFY complex (PDB accession 4AWL) (e) Average NMI values around PBX3 occupancy sites. Also shown is the crystal structure of a PBX-HOXA dimer (PDB accession 1PUF)

Strand-specific accessibility is observed for many other TFs, though not always as clearly as for ATF3. Most bZIP factors, e.g. ATF7 (Supplementary Figure 8), BACH1 (Supplementary Figure 9), CREB1 (Supplementary Figure 16), and CREM (Supplementary Figure 17), exhibit some amount of asymmetry. Some bZIP factors show weaker strand asymmetry, e.g. JUNB (Supplementary Figure 31), JUND (Supplementary Figure 32) and CEBPZ (Supplementary Figure 15).

Other TFs that are known to function as dimers, for example bHLH factors such as USF1/USF2 ^56,57^, also exhibit very strong strand asymmetry of a similar mirror pattern (Supplementary Figures 65 and 66); asymmetry is also observed, though less distinctly, for MYC (Supplementary Figure 41), MAX (Supplementary Figure 35), E4F1 (Supplementary Figure 21), MXI1 (Supplementary Figure 40), and BHLHE40 (Supplementary Figure 12).

We also identify some TFs that exhibit strand-specific footprinting patterns, but only on one strand without a mirrored footprint on the other strand. These include GABPA (Supplementary Figure 29), BATF (Supplementary Figure 10), CBFB (Supplementary Figure 13), CEBPB (Supplementary Figure 14), NFATC3 (Supplementary Figure 43) and the members of the NFY complex ^58,59^ (Supplementary Figures 44 and 45).

A fifth accessibility/footprinting pattern is unique to the YY1 TF (Figure 3i-j), which exhibits very strong footprints combined with strong nucleosome positioning downstream of the YY1 motif, with the top quintile of YY1 occupancy sites exhibiting this pattern most clearly. YY1 is a structural component of enhancer-promoter loops ^60^, and thus its occupancy sites are strongly enriched for promoters of active genes. The positioned nucleosomes observed are likely largely associated with proximal active TSSs. However, YY1 sites nevertheless appear to be a strong positioning factor themselves, because nucleosomes are positioned downstream of a YY1 binding site approximately as strongly as they are around annotated TSS for the top 10% most active genes (compare Figure 3j to Figure 1g). Finally, YY1 exhibits orientation dependence, positioning mucleosomes specifically downstream (to the right) relative to the orientation of the YY1 motif.

We also identify a final unique pattern, exhibited most strongly by the SPI1 TF (Figure 3k-l and Supplementary Figure 60), but also by MAZ (Supplementary Figure 36), BCL11A (Supplementary Figure 11), and to a lesser extent ELF1 (Supplementary Figure 24). In these cases the region upstream (to the left) of the motif is more accessible due to preferential accessibility on the minus strand. This region of increased accessibility is broad and not easily attributable to specific DNA sequence elements.

Finally, we quantified the absolute occupancy of the TF based on the levels of protection against deamination. We note that footprint protection levels arise from a combination of TF occupancy and protection from modification/conversion due to occupancy by other proteins, including nucleosomes and other chromatin-associated factors. The complex origins of protection signals are clearly seen in the case of CTCF, where sites in the top ChIP quintile are much more accessible around the motif than sites in the bottom ChIP quintile (Figure 3b), while the absolute height of the protection footprints at the motif is lowest for the top quintile and highest for the bottom, likely due to protection by nucleosomes occupying sites that are not bound by CTCF. In the extreme, factors like MAFK actually bind to closed chromatin, making their occupancy levels impossible to measure, as the nucleosome itself is already protecting DNA over the binding site.

Given these considerations, we measured occupancy levels using two different metrics that provide lower and upper bounds on occupancy levels – first, the absolute height of footprints, and second, the difference between the absolute height of the footprint and the bottom of the “accessibility well”. The latter metric aims to provide a minimum estimate of occupancy levels, while the former sets the upper bound. Figure 3m summarizes these metrics for all TFs. For many TFs, such as CTCF mentioned above, absolute protection levels are lowest for the strongest ChIP sites, as those are also most accessible as a baseline. TFs in this class also exhibit weak or no correlation with ChIP strength by the top-versus-bottom difference metric. For other very strongly footprinting factors, such as USF1, USF2, NRF1, REST, and ATF3, the top ChIP sites also exhibit the highest levels of protection by both metrics. Thus, for example, for ATF3 and NRF1, we estimate that a minimum of ~30% of top-quintile sites are protected from deamination. We also measured the overall levels of local accessibility for each TF (calculated as the difference between the nucleosomal protection baseline and the bottom of the “accessibility well”), and this metric correlates well with occupancy for all TFs except for MAFK.

### Co-accessibility patterns around transcription factor binding sites

Next, we quantified co-accessibility patterns around TF occupancy sites in the human genome using the GM12878 C→T-ssSMF dataset. To this end, we calculated the Normalized Mutual Information (NMI) metric that we previously developed ^26^, which aims to measure the extent of co-accessibility observed above that expected by chance given average protection/accessibility levels. We computed this metric genome-wide for all pairs of informative positions for all TFs examined (Supplementary Figures 69–72). In all these co-accessibility maps, we observed a globally increased level of co-accessibility at a distance of approximately 10-bp; this likely reflects the length of the DNA double helix turn.

Co-accessibility maps, together with with crystal structures, helped us further rationalize our observations of strand-specific TF occupancy. For example, Figure 4a shows average NMI profiles around NRF1 occupancy sites. NRF1 functions as a homodimer ^61^, but unlike many bZIP factors exhibit only limited footprint strand asymmetry. We find that positions extending to a few basepairs outside the NRF1 recognition motif are strongly preferentially co-accessible. Consistent with this extended co-accessibilty pattern, the NRF1 crystal structure demonstrates that monomers each contact these positions, while also likely protecting substantially both strands over much of the central binding region. Strand-specific co-accessibility maps that show only a slight offset between the co-accessibility footprints on the forward- and reverse-strands are consistent with the uniform protection of both DNA strands observed in the crystal structure.

In contrast, bZIP TF dimers are structured as thin *α*-helices that bind to the DNA major groove, likely leaving the other strand prominently exposed (Figure 4b). For ATF3 we observe a co-accessibility pattern that peaks within the monomer binding sites, with a lower-intensity co-accessibility peak between the two. This monomer-centric pattern is supported by strand-specific co-accessibility maps, which exhibit a strong offset between the co-accessibility footprints on the forward- and reverse-strands with respect to the ATF3 motif.

A distinct pattern is exemplified by the GABPA TF motif, which exhibits increased accessibility over only one strand, with no mirror pattern around its binding site. These patterns might be explained by GABP ^62^ dimers preferentially only contacting one strand, consistent with the GABP-DNA co-crystal structure. This asymmetric, strand-specific binding mode is consistent with the lack of substructure within the co-accessibility signature (Figure 4c), as well as in the stronger and wider co-accessibility footprint on the forward (with respect to the motif) strand.

Co-accessibility maps can also reveal co-occupancy patterns between TF co-factor pairs. For example, Figure 4d shows average NMI profiles for the NFY-A and NFY-B TFs. We observe a co-accessibility signature between the NFY-A motif and the region to the left of it, and a similar co-accessibility pattern at the NFY-B motif, but to the right/downstream of the motif. These proteins function as part of the NFY complex ^58,59^, and bind DNA together. Another example is the pairing of the PBX and HOXA homeodomain factors ^63–65^. Figure 4e shows the average NMI profile for the PBX3 TF in GM12878, exhibiting a prominent co-accessibility peak to the right of the PBX3 recognition motif corresponding to the HOXA co-factor. TFs with such clear co-accessibility signatures also include CEBPZ (Supplementary Figure 69k), USF1 and USF2 (Supplementary Figure 72g-h), and others.

## Discussion

In this study, we first hypothesized that CseDa01 and LbDa02 deaminases might be the ideal molecular probes for mapping chromatin accessibility at high-resolution using *in situ* base conversion due to them exhibitin the strongest reported activity on dsDNA substrates out of all known cytosine deaminases and essentially complete lack of sensitivity to endogenous cytosine methylation. We developed and optimized a variety of protocols for the use of these enzymes to profile chromatin accessibility at the single-molecule level in different short-read and long-read formats, as CseDa01/LbDa02-C→T-SMF. We then focused on CseDa01-C→T-XL-ssSMF in the GM12878 lymphoblastoid cell line to map the fine-grained strand-specific features of transcription factor occupancy.

Several emerging features of the dsDNA deaminase that we use (which are likely to be shared by all such enzymes) are notable. We find very strong strand-specific protection/accessibility patterns both for nucleosomes and individual transcription factor occupancy sites, much stronger than those previously observed with m^6^A methylation. However, we consistently find that CseDa01 treatment of chromatin results in the conversion of a somewhat higher fraction of nucleotides at baseline, i.e. over nucleosomes, than is observed by GpC methylation of chromatin using the M.CviPI enzyme. These differences probably have to do with the different mechanisms of actions of MTases and deaminases. Methylation by M.CviPI methyltransferase involves making contact with both strands, likely requiring more complete physical access to DNA for efficient labeling. While the m^6^A MTases are not sequence-specific and will label any A positions on just one strand, they belong to the same general class of enzymes and may have similar requirements for access to dsDNA. In contrast, deaminases such as CseDa01 act on both dsDNA and on ssDNA, and are therefore likely more active on DNA substrates with more limited exposure, or exposure to only one strand. Therefore efficient probing of strand-specific accessibility comes at the cost of higher conversion rates over otherwise strongly protected substrates, such as nucleosomes.

We used CseDa01-C→T-XL-ssSMF to provide lower-bound and upper-bound estimates for the occupancy of 64 humans TFs in the GM12878 cell line. However, the final quantitative measurement of TF occupancy across the genome will require a more comprehensive analysis, possibly using different approaches and tools. Because of the rapid dissociation kinetics of many human TFs, we used formaldehyde crosslinking for the experiments presented in this study. However, fixation does not strictly “freeze” a snapshot of the accessibility landscape at a given moment, but integrates it over an extended period of time (in this case, formaldehyde crosslinking was carried out for 15 minutes). We expect a combination of much more highly active/concentrated probes on native chromatin and/or more rapid crosslinking approaches ^66,67^ to address these remaining open questions.

## Materials and Methods

Except for when explicitly stated otherwise, all analyses were carried out using custom-written Python or R scripts.

### Cell lines and cell culture

GM12878 and K562 cells were grown following established protocols ^1,2^, i.e. RPMI 1640 supplemented with 15% and 10% FBS, respectively, and 1% penn-strep.

Interferon induction was peformed by adding 10 ng/mL of IFN*β* (Peprotech, 300-02BC) for 6 hours as previously described ^23^.

### dsDNA deaminase production (PURExpress)

CseDa01 and LbDa02 enzymes were produced using PURExpress as follows. First, codon-optimized coding sequences as described by Vaisvila et al. 2024 were ordered as linear gBlocks from IDT with flanking sequences added containing the T7 promoter at 5’ end (GCGAATTAATACGACTCACTATAGGGCTTAAG-TATAAGGAGGAAAAAAT) and T7 terminator at 3’ end (CTAGCATAACCCCTCTCTAAACGGAGGGGTT-TATTTGG). The gBlock sequences were the following: CsaDa01:

GCGAATTAATACGACTCACTATAGGGCTTAAGTATAAGG AGGAAAAAATATGGATGATGCACTGGCGGAAAAGGCGCG TCAAGCCCACAACGTGCTGAAACGCAACATCCCGGGTAA ACCGGAAATCGGCTTTAACCAGTCCACGGTGTCTATTGG CGAAGGTACTCTGCCGTCTGGCGAGAAACAACTGTTTGC TAGCGGTAACGGTGCTAAACTGACCCCGGCCCAGCGTGC AAAACTGATCGAGCTGGGTGTTCCACCAGAAAACATTTA TTCTGGCGTAGCGTACATGAAGAAACCAGTTAACAAATC CAAAGTTAAATCCGTGCGTAAAGCCTACAACCTGGAAAA CCACGCTGAGCGTGTAATTATTCGCAACGCACCGAAAGG TACGAAATTCAAGAAATGGGGTATCTCCTGGGCGAAATA CCAGAAGAACGAACCGTGTTCTAACTGTAAACCTCATGT ACAGTGTGCATCCTAACTAGCATAACCCCTCTCTAAACG GAGGGGTTTATTTGG

LbDa02:

GCGAATTAATACGACTCACTATAGGGCTTAAGTATAAGG AGGAAAAAATATGAACGACGGTAACTCCGAATCTGGCTC CGATGCGCTGGACAAGAAAATTAAACAGGCAGAACAGGC CCAAAAGAAAGCGTCCGAACTGAGCGACAAATACGGCTA TGGCACTGCTCCTAAGAAAACTGTGGCTTCTGACGGTAC CATCCATACTCTGAGCGGTTGGAAACCACTGGATGAGAA CAGCGAATTCACCCGTGTCTCTACTGATGACGTAATTAA TAAAAGCAACGATATTGGCCACAAACTGCGTAACGCGGG TGCGAACGATCAGGGTGTGCCAGGTCGTTACAACGCCAG CCACGCTGAAAAGCAACTGTCTATTATCTCTGATAAACC TATTGGTATCAGCCAGCCGATGTGTACCGATTGTCAAGG TTATTTCAAATCCCTGGCTATCGGCGAGGGTAAAGAATA CGTAACTGCTGATCCGAACTACATCCGCATCTTTTCTCC AGATGGTACTGTGAACGAGATTGAACGTCAACGTTAACT AGCATAACCCCTCTCTAAACGGAGGGGTTTATTTGG

The gBlock sequences were PCR-amplified using flanking sequence primers, then purified using the MinElute PCR Purification Kit (Qiagen), quantified using a Qubit 1X dsDNA HS Assay Kit (ThermoFisher) and confirmed to be the right size using an Agilent TapeStation.

Then 100-200 ng of input DNA per reaction were used to run multiple PUREexpress (NEB #E6800S) reactions for ≥2 hours at 37 °C. The reactions were pooled and deamination activity tested for each such batch on naked DNA and/or chromatin.

### Purification of recombinant CseDa01 enzyme

To express the CseDa01 enzyme in *E. coli*, a pQE-30-based (Qiagen) expression vector was constructed, incorporating a T7 promoter-driven, N-terminal 6×His-tagged CseDa01 gene along with a T5 promoter-driven CseDa immunity protein gene. This plasmid was transformed into the *E. coli* expression strain T7 Express lysY/Iq (C3013, NEB). Protein purification was carried out using a denaturation-refolding protocol adapted from Mok et al. ^68^.

### Deamination of naked DNA

#### Genomic DNA isolation

Naked gDNA was isolated from K562 cells using the NEB Monarch HMW DNA Extraction Kit kit (NEB #T3050L).

#### Enzymatic treatment

For tests of CseDa01 and LbDa02 activity, 1 *µ*g of purified HMW gDNA was incubated with 1 *µ*L of enzyme for 3 hours at 37 °C in a total volume of 100 *µ*L of deamination buffer ^30^.

The reaction was stopped by adding 5× volume of PB Buffer and then the Qiagen MinElute kit was used for DNA purification.

DNA concentration was measured using a Qubit 1X dsDNA kit (Thermo Fisher Scientific).

#### Deaminated gDNA transposition libraries generation

Short-read libraries were prepared using Tn5 transposition as follows. Purified SMF DNA was diluted to a total amount of 20 ng in a volume of 11.5 *µ*L, then 12.5 *µ*L of 2× TD Buffer and 1 *µ*L Tn5 (produced locally following the protocol described by Picelli et al. ^69^) was added, and the mixture was incubated at 55 °C for 5 minutes. The reaction was stopped by adding 250 *µ*L of PB Buffer and the transposition product was purified using the Qiagen MinElute kit, eluting in 20 *µ*L EB Buffer.

PCR amplification was carried out by mixing 20 *µ*L transposed DNA, 2.5 *µ*L i5 and 2.5 *µ*L i7 PCR primers (previously described ^70^), and 25 *µ*L 2× Q5U PCR Master Mix (i.e. a uracil-tolerant polymerase), with the following settings:

72 °C for 5 minutes
98 °C for 30 seconds
10 cycles of:

98 °C for 10 seconds
63 °C for 30 seconds
72 °C for 45 seconds
Hold at 4 °C

Final libraries were purified and QC-ed using a Qubit Qubit 1X dsDNA kit (Thermo Fisher Scientific) and an Agilent TapeStation, and sequenced on a NextSeq 550 instrument.

#### Data processing

Adapters were trimmed from reads using Trimmomatic ^71^ (version 0.36). Trimmed reads were aligned against the hg38 version of the human genome assembly using bwa-meth with default settings. Duplicate reads were removed using picard-tools (version 1.99) if sequencing in a paired-end format. Methylation calls were extracted using MethylDackel (https://github.com/dpryan79/MethylDackel).

### C**→**T-SMF experiments

#### Native SMF

For native SMF, 1×10^6^ cells were first centrifuged at 1,000 *g* for 2 minutes, then resuspended in 200 *µ*L RSB-Lysis buffer (10 mM Tris-HCl pH 7.4, 10 mM NaCl, 3 mM MgCl_2_, 0.1% IGEPAL CA-630, 0.1% Tween-20, 0.01% Digitonin) and incubated on ice for 3 minutes. Subsequently, 800 *µ*L RSB-wash Buffer (10 mM Tris-HCl pH 7.4, 10 mM NaCl, 3 mM MgCl_2_, 0.1% Tween-20) were added, and nuclei were pelleted by centrifugation at 500 *g* for 5 minutes at 4 °C.

Nuclei were then resuspended in 250 *µ*L 1×APOBEC Buffer (from NEB EM-seq kit: NEB #E7120). Deamination treatment was carried out by adding 10 *µ*L concentrated CseDa01, then incubating at 37 °C for 15 minutes.

The reaction was stopped by adding 800 *µ*L PB Buffer and then the Qiagen MinElute kit was used to purify deaminated DNA.

#### XL-SMF

For crosslinked SMF, cells were first centrifuged at 1,000 *g* for 2 minutes, then resuspended in 1 mL 1×PBS with 27 *µ*L 37% formaldehyde added (to a final concentration of 1%). Crosslinking was carried out for 15 minutes at room temperature on a rotator, then quenched with 100 *µ*L 2.5 M glycine solution. Crosslinked cells were centrifuged at 1,000 *g* for 2 minutes, resuspended in 1 mL 1×PBS, and centrifuged again.

Permeabilization was carried out either by resuspending cells in Lysis Buffer (10 mM Tris-HCl pH 8.0, 10 mM NaCl, 0.2% Igepal CA630) and incubating on ice for 10 minutes, then centrifuging for 2 minutes at 1,000 g, or by following the omni-ATAC lysis protocol ^70,72^.

Cells (200,000 cells were used as input) were then resuspended in 1 mL of 1×APOBEC Buffer (from the NEB EM-seq kit). Deamination treatment was carried out by adding 2 *µ*L RNAse A and 10 *µ*L PURExpress CseDa01 or LbDa01, then incubating at 37 °C for 3 hours with shaking at 800 rpm in a Thermomixer.

For purified CseDa01, 1×10^6^ cells and 1 *µ*L of concentrated CseDa01 were used.

After deamination, cells were again pelleted by centrifugation at 1,500 *g* for 2 minutes, then resuspended in 200 *µ*L IP Elution Buffer (1% SDS, 0.1 M NaHCO_3_); 2 *µ*L Proteinase K were added, and reverse crosslinking was carried out overnight at 65 °C.

DNA was then isolated using the MinElute kit (Qiagen) using 600 *µ*L PB buffer and following the manufacturer’s instructions, eluting in 52 *µ*L EB buffer. DNA concentration was measured using the Qubit 1X dsDNA kit (Thermo Fisher Scientific).

#### Transposition short-read library preparation

Short-read libraries were prepared using Tn5 transposition as described above. Purified SMF DNA was diluted to a total amount of 20 ng in a volume of 11.5 *µ*L, then 12.5 *µ*L of 2× TD Buffer and 1 *µ*L Tn5 were added, and the mixture was incubated at 55 °C for 5 minutes. The reaction was stopped by adding 125 *µ*L of PB Buffer and the transposition product was purified using the Qiagen MinElute kit, eluting in 20 *µ*L EB Buffer.

PCR amplification was carried out by mixing 20 *µ*L transposed DNA, 2.5 *µ*L i5 and 2.5 *µ*L i7 PCR primers, and 25 *µ*L 2× Q5U PCR Master Mix (i.e. a uracil-tolerant polymerase).

Final libraries were purified and QC-ed using a Qubit Qubit 1X dsDNA kit (Thermo Fisher Scientific) and an Agilent TapeStation, and sequenced on a NextSeq 550 instrument to assess global deamination profile properties.

#### Illumina ssDNA library preparation

For single-stranded libraries, SMF DNA was first sheared using a Covaris E220 instrument to an average size of 400-500 bp.

The xGen Methyl-Seq DNA Library Prep Kit (IDT)) was then used to generated Illumina-compatible libraries, which were sequenced on a sequenced on a NextSeq 550 instrument in a 1×300 bp format. SMF samples were split into four equal subsamples, and each of these samples was converted into a separate library, in order to maximize the complexity of the final dataset.

#### Ultima library conversion

Illumina libraries were converted into Ultima-compatible libraries using a constant primer (CTGTGTGCCTTGGCAGTCTCAGCTCAGACGTGTGCTCTTCC-GATC*T) and an index primer (CCATCTCATCCCTGCGTGTCTCCGACTGCACXXXXXXXXXXXGATC-TACACGACGCTCTTCCGATC*T, where the portion indicated as “XXXXXXXXXXX” corresponds to the index sequences). For each library, ~10 ng were used as input, diluted in a 20 *µ*L volume, then mixed with 2.5 *µ*L of each primer and 25 *µ*L of 2× NEB Next High Fidelity PCR Master Mix (NEB Cat # M0541L). Libraries were amplified using the following settings:

98 °C for 45 seconds
7 cycles of:

98 °C for 15 seconds
60 °C for 30 seconds
72 °C for 60 seconds
72 °C for 10 minutes
Hold at 4 °C

Ultima libraries were sequenced by Ultima Genomics.

#### Whole-genome DNA amplification

Whole-genome amplification (WGA) of SMF DNA was carried out using the Qiagen REPLI-g kit following manufacturer’s instructions.

#### Targeted amplification

Targeted amplification was carried out starting with 100 ng amplified SMF gDNA diluted in 8 *µ*L total volume with ultrapute H_2_O, mixed with 1 *µ*L of forward primer, 1 *µ*L of reverse primer, and 10 *µ*L of 2× Q5U PCR Master Mix, with the following settings:

98 °C for 45 seconds
28 cycles of:

98 °C for 20 seconds
*T_m_* for 30 seconds
72 °C for 5 minutes
72 °C for 10 minutes
Hold at 4 °C

The primer pairs used were GGTAGGATGGAGGAATGA and CTCCCTCTCACCAAAAC for the *IFI6* locus (*T_m_* = 62 °C), GAAGGAYGGAAGGAAGA and CACTTCCCCTRCCTTA for the *USP18* locus (*T_m_*= 60 °C), and GAATGGGTGAAGAGAGAGG and TCCTT-TACCTACCCAAATCT for the *IFIT3* locus (*T_m_* = 60 °C).

Amplification was verified using a D5000 TapeStation kit.

#### Nanopore sequencing

Amplicons were mixed and sequenced using the Oxford Nanopore Technologies Flongle sequencing platform (FLG114 with the SQK-LSK114 kit).

Whole-genome SMF samples were sequenced using the Oxford Nanopore Technologies MinION platform using the SQK-LSK114 kit.

### C**→**T-SMF data processing

#### Data processing of short-read libraries

Short read SMF sequencing data was processed as previously described ^73^ with the modification that accessible/protected C positions were flipped (while in the EM-seq and bisulfite readouts used in cytosine methylation-based SMF unmethylated Cs are converted, in C→T-SMF base conversion is direct).

Briefly, adapters were trimmed from reads using Trimmomatic ^71^ (version 0.36). Trimmed reads were aligned against the hg38 version of the human genome using bwa-meth with default settings. Duplicate reads were removed using picard-tools (version 1.99) if sequencing in a paired-end format. Methylation calls were extracted using MethylDackel (https://github.com/dpryan79/ MethylDackel). Additional analyses and visualization were carried out using custom-written Python scripts (https://github.com/georgimarinov/GeorgiScripts) as previously described ^73^.

#### Nanopore sequencing data processing

Nanopore reads were mapped against the hg38 version of the human genome using minimap2 ^74^.

Additional analyses and visualization were carried out using custom-written Python scripts (https://github.com/georgimarinov/GeorgiScripts) as previously described ^36^.

### NOMe-seq/GpC-SMF

NOMe-seq/GpC-SMF experiments were carried out as previously described ^73^ with some modifications.

Briefly, 1×10^6^ cells were pelleted by centrifugation, resuspended in 1 mL 1× PBS and pelleted again, then resuspended in 200 *µ*L RSB-Lysis buffer (10 mM Tris-HCl pH 7.4, 10 mM NaCl, 3 mM MgCl_2_, 0.1% IGEPAL CA-630, 0.1% Tween-20, 0.01% Digitonin) and incubated on ice for 3 minutes. Then 1200 *µ*L RSB-wash Buffer (10 mM Tris-HCl pH 7.4, 10 mM NaCl, 3 mM MgCl_2_, 0.1% Tween-20) were added, and nuclei were pelleted by centrifugation at 500 *g* for 10 minutes at 4 °C. Nuclei were then resuspended in 100 *µ*L M. CviPI Reaction Buffer and 200 U of M.CviPI GpC MTase (NEB) were added, together with SAM (S-adenosylmethionine) at a final concentration of 0.6 mM. The reaction was incubated at 37 °C for 7.5 minutes in a Thermomixer at 1,000 rpm, then another 100 U of enzyme and more SAM were added and the incubation was continued for another 7.5 minutes.

The reaction was stopped by adding 175 *µ*L of the Cell Lysis Buffer from the NEB Monarch gDNA extraction kit together with 3 *µ*L RNase A and 1 *µ*L Proteinase K, and gDNA purification was carried out following the instructions for the NEB Monarch kit.

Final sequencing libraries were prepared on Covaris-sheared DNA using the NEB EM-seq kit, and sequenced as 2×150mers on an Illumina NovaSeq instrument (through Novogene).

### NOMe-seq/GpC-SMF data processing

NOMe-seq/GpC-SMF sequencing data was processed as previously described ^73^. Briefly, adapters were trimmed from reads using Trimmomatic ^71^ (version 0.36). Trimmed reads were aligned against the hg38 version of the human genome using bwa-meth with default settings. Duplicate reads were removed using picard-tools (version 1.99) if sequencing in a paired-end format. Methylation calls were extracted using MethylDackel (https://github.com/dpryan79/MethylDackel). Additional analyses and visualization were carried out using custom-written Python scripts (https://github.com/georgimarinov/GeorgiScripts) as previously described ^73^.

### ATAC-seq experiments

ATAC-seq experiments were carried out as previously described ^70^.

Briefly, 5×10^4^ cells were pelleted by centrifugation, resuspended in 200 *µ*L 1× PBS and pelleted again, then resuspended in 50 *µ*L RSB-Lysis buffer (10 mM Tris-HCl pH 7.4, 10 mM NaCl, 3 mM MgCl_2_, 0.1% IGEPAL CA-630, 0.1% Tween-20, 0.01% Digitonin) and incubated on ice for 3 minutes. Then 450 *µ*L RSB-wash Buffer (10 mM Tris-HCl pH 7.4, 10 mM NaCl, 3 mM MgCl_2_, 0.1% Tween-20) were added, and nuclei were pelleted by centrifugation at 500 *g* for 5 minutes at 4 °C.

Tagmentation of chromatin followed by resuspending nuclei with 25 *µ*L 2× TD buffer (20 mM Tris-HCl pH 7.6, 10 mM MgCl_2_, 20% Dimethyl Formamide), 2.5 *µ*L transposase (custom produced ^69^) and 22.5 *µ*L nuclease-free H_2_O, and incubating at 37 °C for 30 min in a Thermomixer at 1000 RPM. Transposed DNA was isolated using the Qiagen MinElute kit and PCR amplified as descrbed above using the following procedure:

72 °C for 5 minutes
98 °C for 30 seconds
10 cycles of:

98 °C for 10 seconds
63 °C for 30 seconds
72 °C for 30 seconds
Hold at 4 °C

Libraries were then sequenced on an Illumina NextSeq instrument as 2×36mers.

### ATAC-seq data processing

ATAC-seq data processing was carried out as previously described ^75^. Briefly, demultipexed fastq files were mapped to the hg38 assembly of the human genome as 2×36mers using Bowtie ^76^ with the following settings: -v 2 -k 2 -m 1 --best --strata. Duplicate reads were removed using picard-tools (version 1.99). Peak calling was carried out using MACS2 ^77^ as previously described ^75^. Additional analyses and visualization were carried out using custom-written Python scripts (https://github.com/georgimarinov/GeorgiScripts).

### Transcription factor motif mapping

Motif annotations were obtained form the CIS-BP database ^78^ (version 1.0.2). Motifs were mapped against the hg38 version of the human genome assembly using the FIMO program ^79^ in the MEME-SUITE of motif analysis tools ^80^.

### Co-accessibility assessment

Co-accessibility was measured between each pair of informative positions using the normalized mutual information (NMI) metric previously described in Shipony & Marinov et al. ^26^.

### External sequencing datasets

Additional datasets and annotations were obtained from the ENCODE portal as indicated in the corresponding figures and figure legends.

## Data availability

Sequencing data associated with the manuscript is available through GEO accession GSE301497.

## Author contributions

G.K.M. conceived project, designed and carried out experiments and data analysis and wrote the manuscript with input from all authors. T.W. produced deaminase enzymes. M.M.S., M.C. and Z.S provided key reagents. J.M.S. and B.D. designed and carried out experiments. W.J.G. and A.K. supervised the project.

## Acknowledgments

This work was supported by NIH grants U01HG011762, R01 NS128028, DP1HG013599, R01HL171611, and R01HG013317 to W.J.G..

## Supplementary Materials

### Supplementary Figures

**Supplementary Figure 1:**
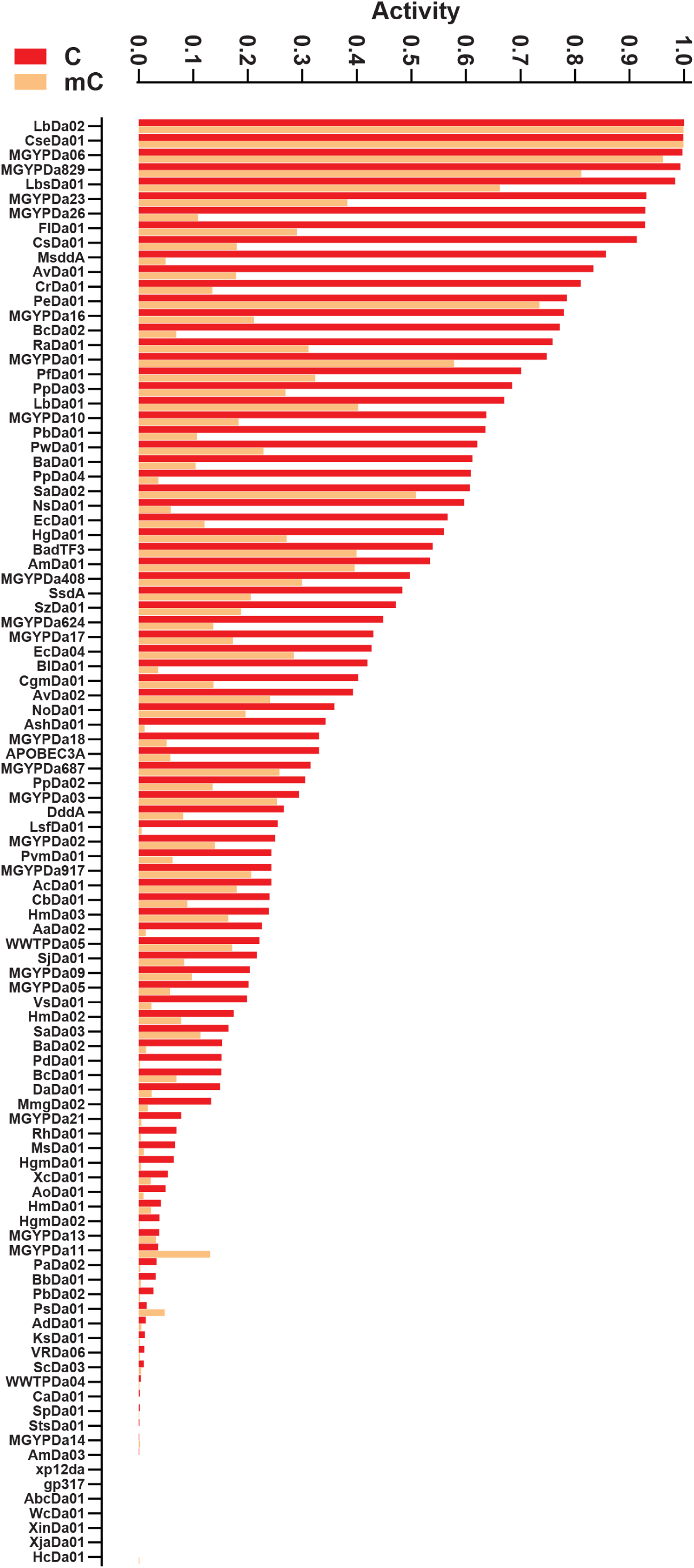
Relative activity of various deaminases against unmethylated and methylated dsDNA substrates. Data was obtained from Vaisvila et al. 2024 ^30^.

**Supplementary Figure 2:**
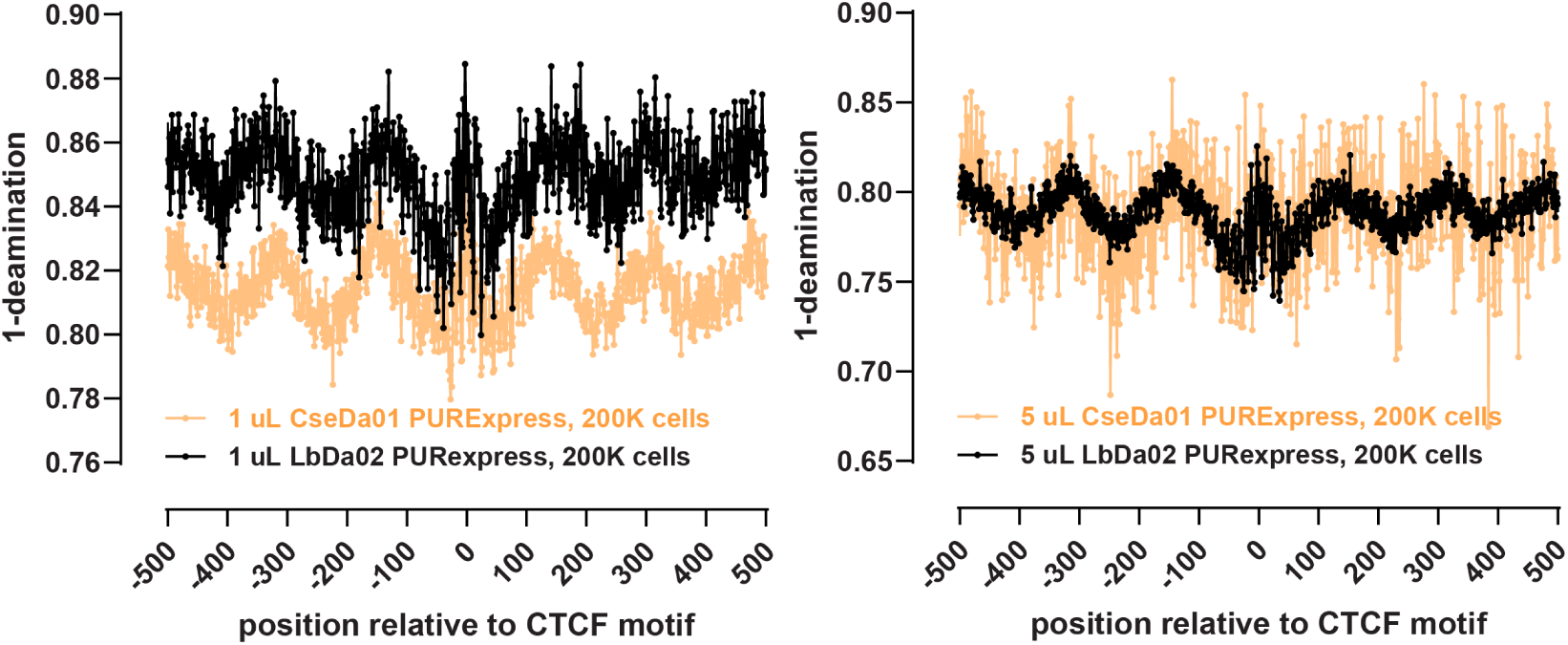
High-resolution chromatin accessibility profiling using the non-specific CseDa01 and LbDa02 dsDNA deaminases. Shown are metaprofiles around CTCF motifs within CTCF ChIP-seq peaks for experiments carried out with PURExpress CseDa01 preps or PURExpress LbDa02 and the indicated amount of enzyme and number of cells (K562 cells, crosslinked).

**Supplementary Figure 3:**
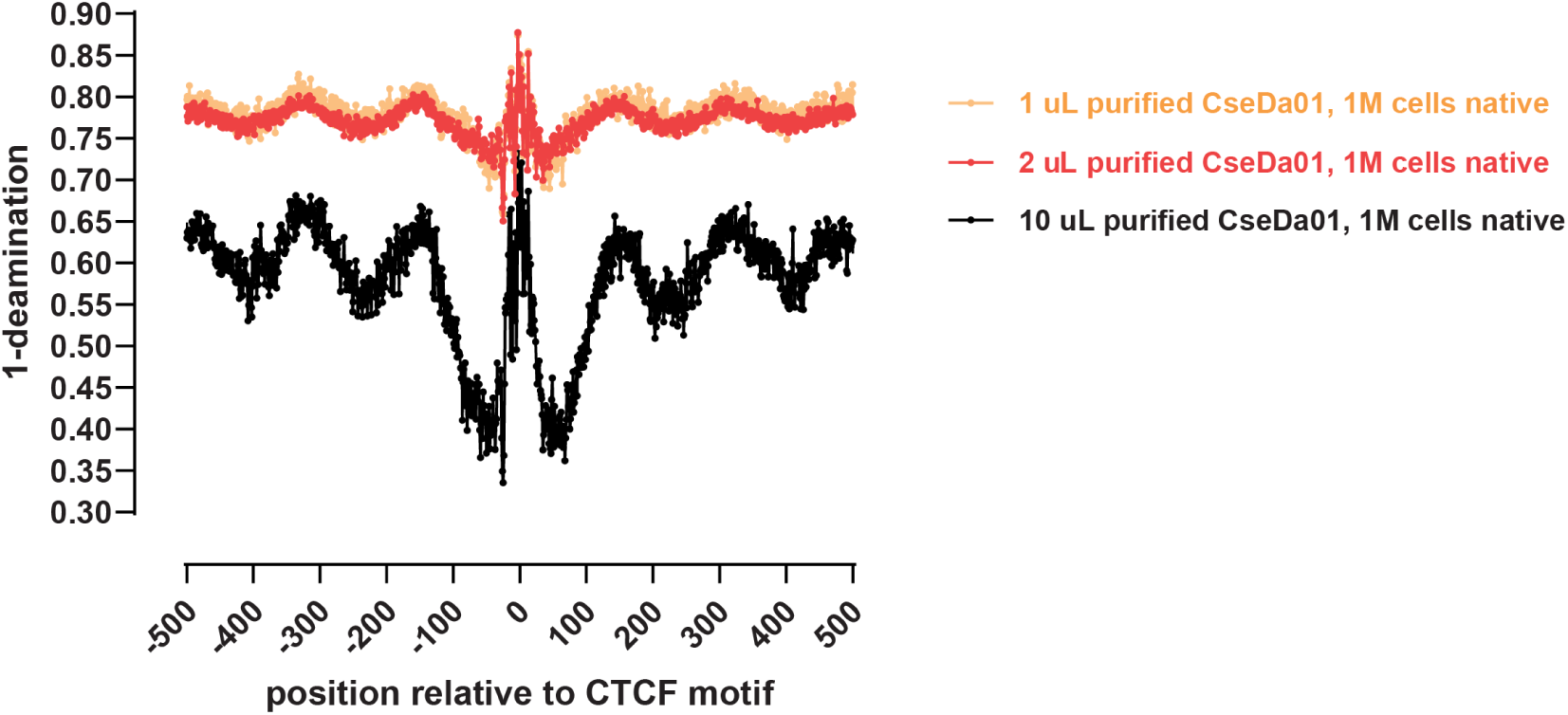
Native high-resolution chromatin accessibility profiling using the non-specific CseDa01 dsDNA deaminase. Shown are metaprofiles around CTCF motifs within CTCF ChIP-seq peaks for experiments carried out with purified CseDa01 and the indicated amount of enzyme and number of cells (K562 cells, native).

**Supplementary Figure 4:**
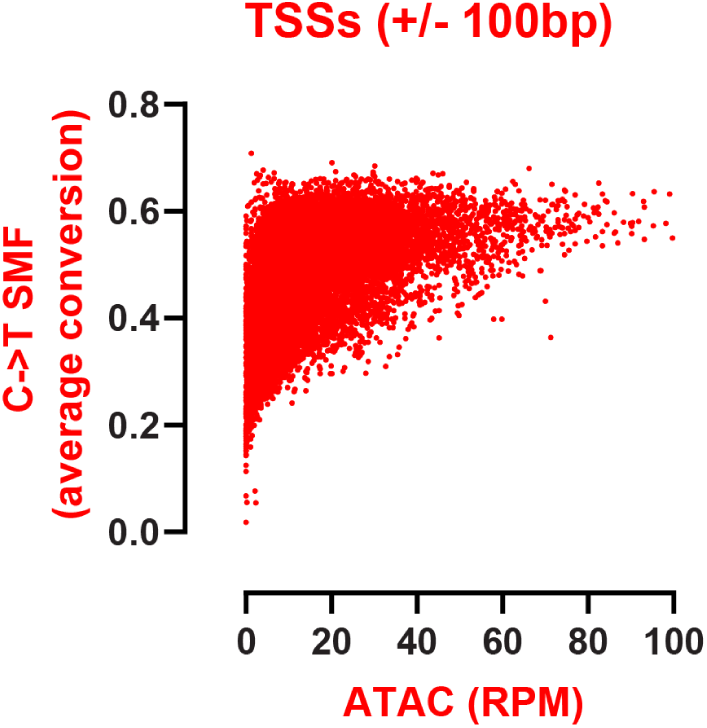
Correspondence between short-read CseDa01 C→T-SMF and ATAC-seq genome-wide profiles. Shown are RPKM values over annotated RefSeq TSSs (±100 bp) and average deamination rates for the same windows.

**Supplementary Figure 5:**
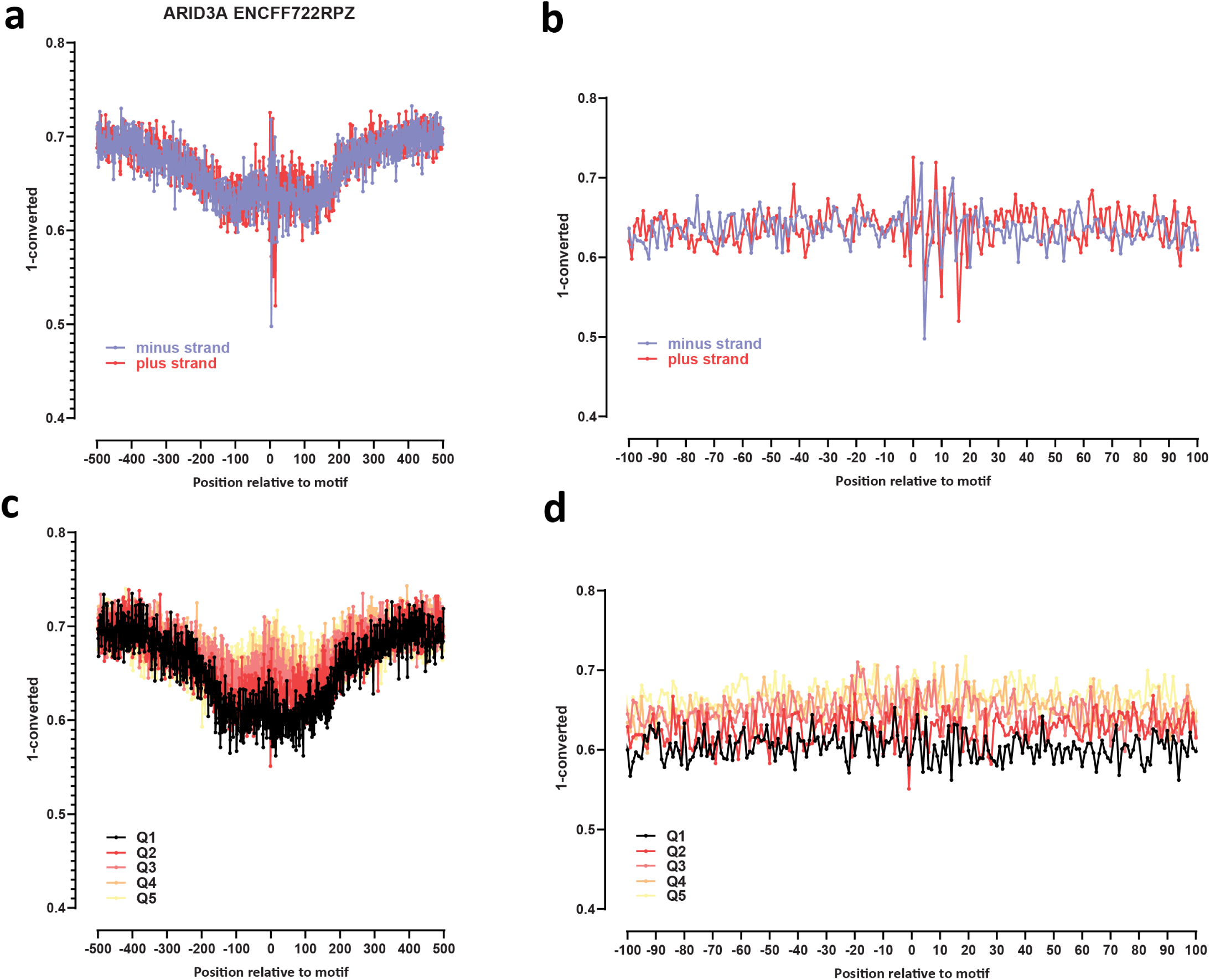
Genome-wide footprint profiles for the ARID3A transcription factor. GM12878 ssDNA-CseDa01-XL-SMF datasets are shown. (a) Genome-wide footprint profiles over the closest motif to ENCODE ChIP-seq peaks (ENCODE dataset ID indicated on top). Profiles are generated in a strand-aware way with respect to the motif orientation in the genome, and are shown separately for forward and reverse-strand reads. (b) Same as in (a) but zoomed in. The transcription factor sequence recognition motif is shown on top. (c) Genome-wide footprint profiles over the closest motif to ENCODE ChIP-seq peaks for the TF divided into quintiles based on ChIP-seq signal strength. (d) Same as in (c) but zoomed in.

**Supplementary Figure 6:**
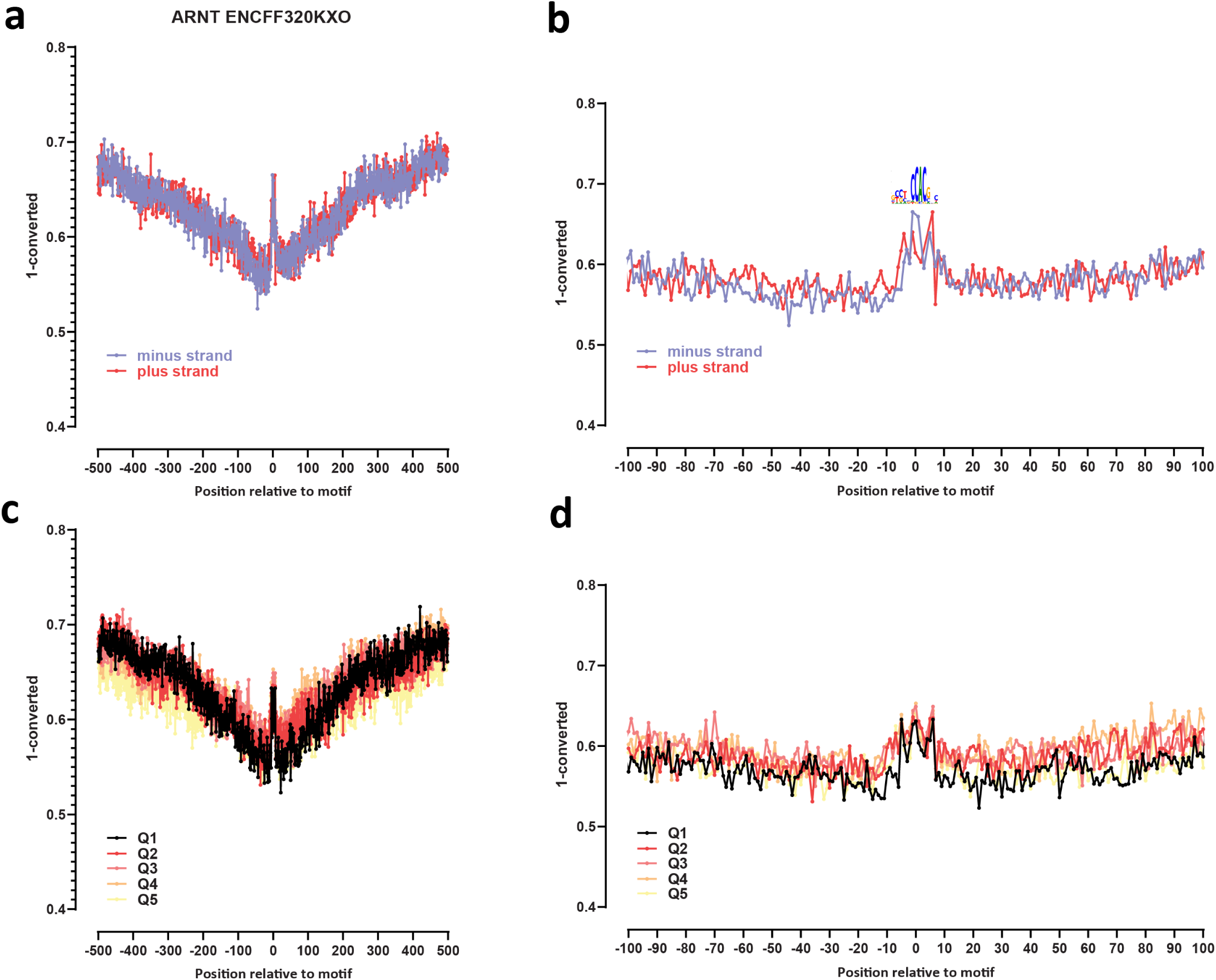
Genome-wide footprint profiles for the ARNT transcription factor. GM12878 ssDNA-CseDa01-XL-SMF datasets are shown. (a) Genome-wide footprint profiles over the closest motif to ENCODE ChIP-seq peaks (ENCODE dataset ID indicated on top). Profiles are generated in a strand-aware way with respect to the motif orientation in the genome, and are shown separately for forward and reverse-strand reads. (b) Same as in (a) but zoomed in. The transcription factor sequence recognition motif is shown on top. (c) Genome-wide footprint profiles over the closest motif to ENCODE ChIP-seq peaks for the TF divided into quintiles based on ChIP-seq signal strength. (d) Same as in (c) but zoomed in.

**Supplementary Figure 7:**
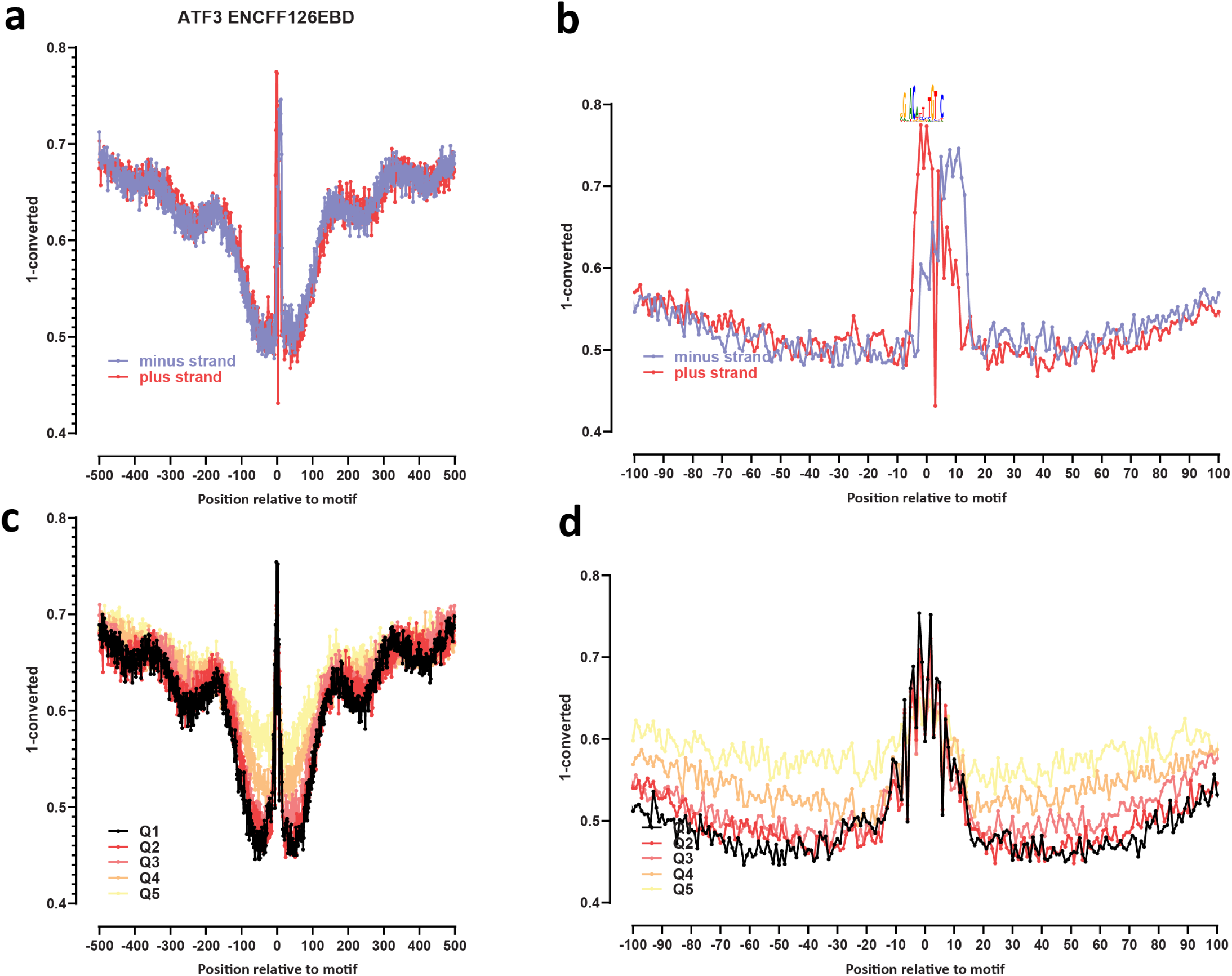
Genome-wide footprint profiles for the ATF3 transcription factor. GM12878 ssDNA-CseDa01-XL-SMF datasets are shown. (a) Genome-wide footprint profiles over the closest motif to ENCODE ChIP-seq peaks (ENCODE dataset ID indicated on top). Profiles are generated in a strand-aware way with respect to the motif orientation in the genome, and are shown separately for forward and reverse-strand reads. (b) Same as in (a) but zoomed in. The transcription factor sequence recognition motif is shown on top. (c) Genome-wide footprint profiles over the closest motif to ENCODE ChIP-seq peaks for the TF divided into quintiles based on ChIP-seq signal strength. (d) Same as in (c) but zoomed in.

**Supplementary Figure 8:**
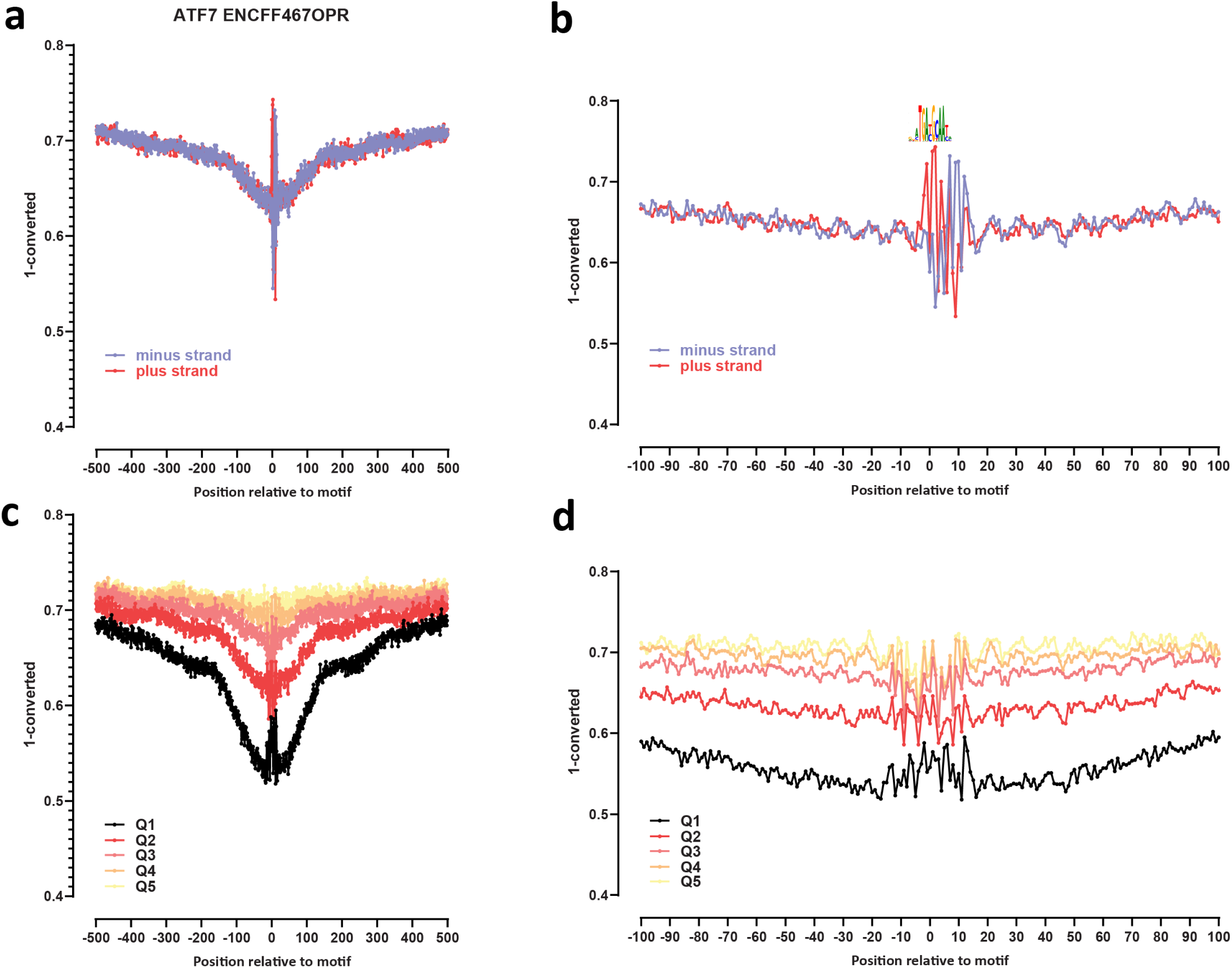
Genome-wide footprint profiles for the ATF7 transcription factor. GM12878 ssDNA-CseDa01-XL-SMF datasets are shown. (a) Genome-wide footprint profiles over the closest motif to ENCODE ChIP-seq peaks (ENCODE dataset ID indicated on top). Profiles are generated in a strand-aware way with respect to the motif orientation in the genome, and are shown separately for forward and reverse-strand reads. (b) Same as in (a) but zoomed in. The transcription factor sequence recognition motif is shown on top. (c) Genome-wide footprint profiles over the closest motif to ENCODE ChIP-seq peaks for the TF divided into quintiles based on ChIP-seq signal strength. (d) Same as in (c) but zoomed in.

**Supplementary Figure 9:**
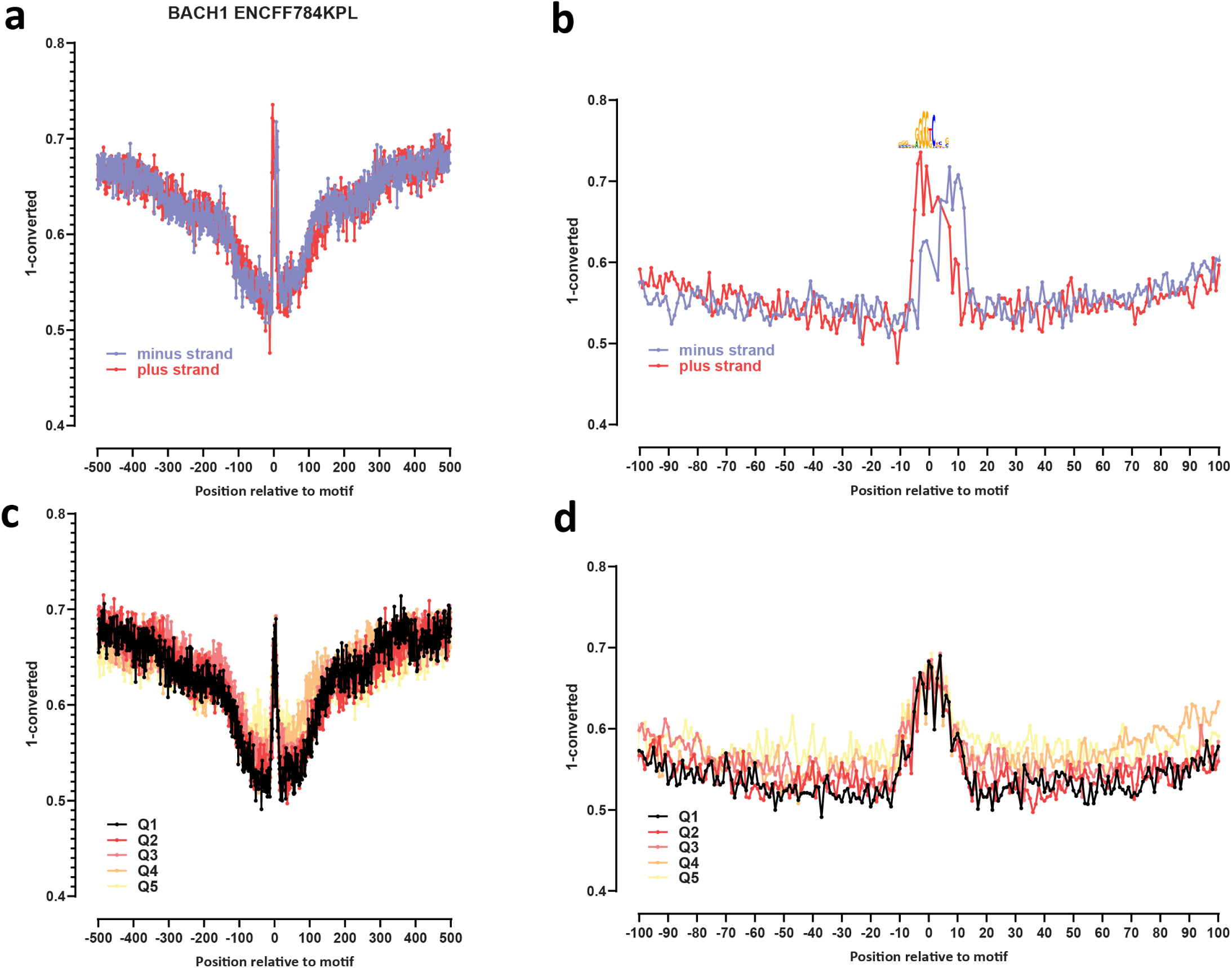
Genome-wide footprint profiles for the BACH1 transcription factor. GM12878 ssDNA-CseDa01-XL-SMF datasets are shown. (a) Genome-wide footprint profiles over the closest motif to ENCODE ChIP-seq peaks (ENCODE dataset ID indicated on top). Profiles are generated in a strand-aware way with respect to the motif orientation in the genome, and are shown separately for forward and reverse-strand reads. (b) Same as in (a) but zoomed in. The transcription factor sequence recognition motif is shown on top. (c) Genome-wide footprint profiles over the closest motif to ENCODE ChIP-seq peaks for the TF divided into quintiles based on ChIP-seq signal strength. (d) Same as in (c) but zoomed in.

**Supplementary Figure 10:**
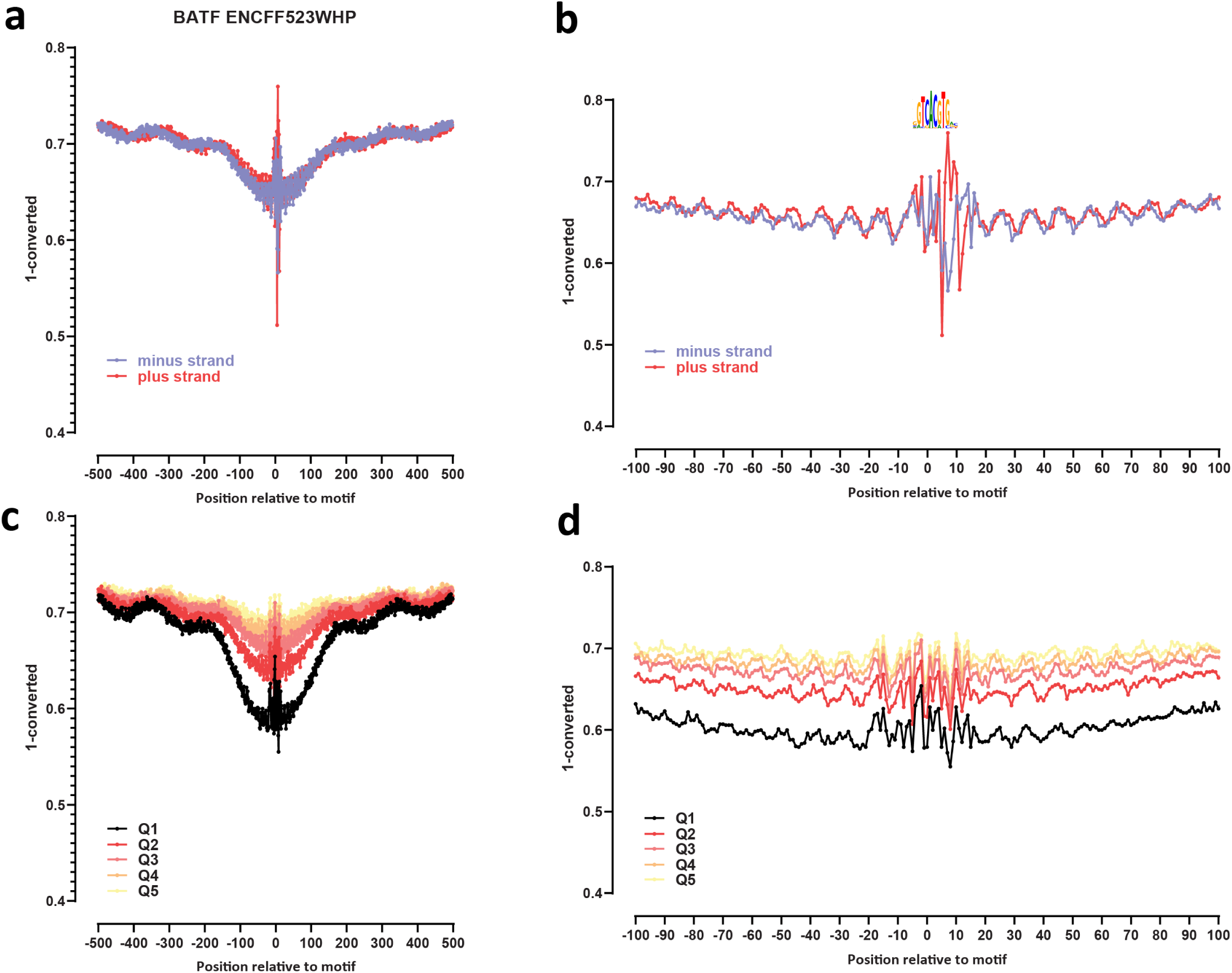
Genome-wide footprint profiles for the BATF transcription factor. GM12878 ssDNA-CseDa01-XL-SMF datasets are shown. (a) Genome-wide footprint profiles over the closest motif to ENCODE ChIP-seq peaks (ENCODE dataset ID indicated on top). Profiles are generated in a strand-aware way with respect to the motif orientation in the genome, and are shown separately for forward and reverse-strand reads. (b) Same as in (a) but zoomed in. The transcription factor sequence recognition motif is shown on top. (c) Genome-wide footprint profiles over the closest motif to ENCODE ChIP-seq peaks for the TF divided into quintiles based on ChIP-seq signal strength. (d) Same as in (c) but zoomed in.

**Supplementary Figure 11:**
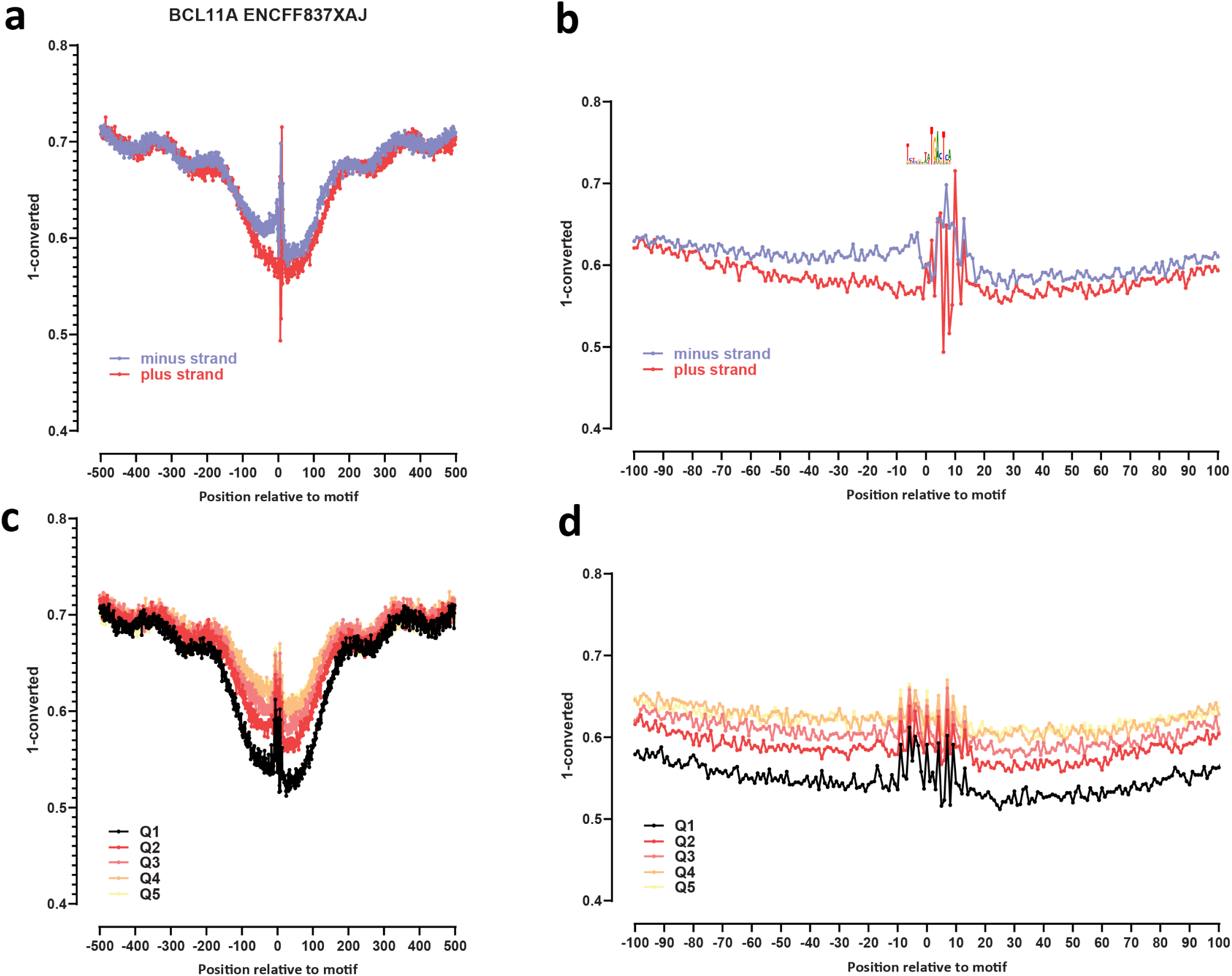
Genome-wide footprint profiles for the BCL11A transcription factor. GM12878 ssDNA-CseDa01-XL-SMF datasets are shown. (a) Genome-wide footprint profiles over the closest motif to ENCODE ChIP-seq peaks (ENCODE dataset ID indicated on top). Profiles are generated in a strand-aware way with respect to the motif orientation in the genome, and are shown separately for forward and reverse-strand reads. (b) Same as in (a) but zoomed in. The transcription factor sequence recognition motif is shown on top. (c) Genome-wide footprint profiles over the closest motif to ENCODE ChIP-seq peaks for the TF divided into quintiles based on ChIP-seq signal strength. (d) Same as in (c) but zoomed in.

**Supplementary Figure 12:**
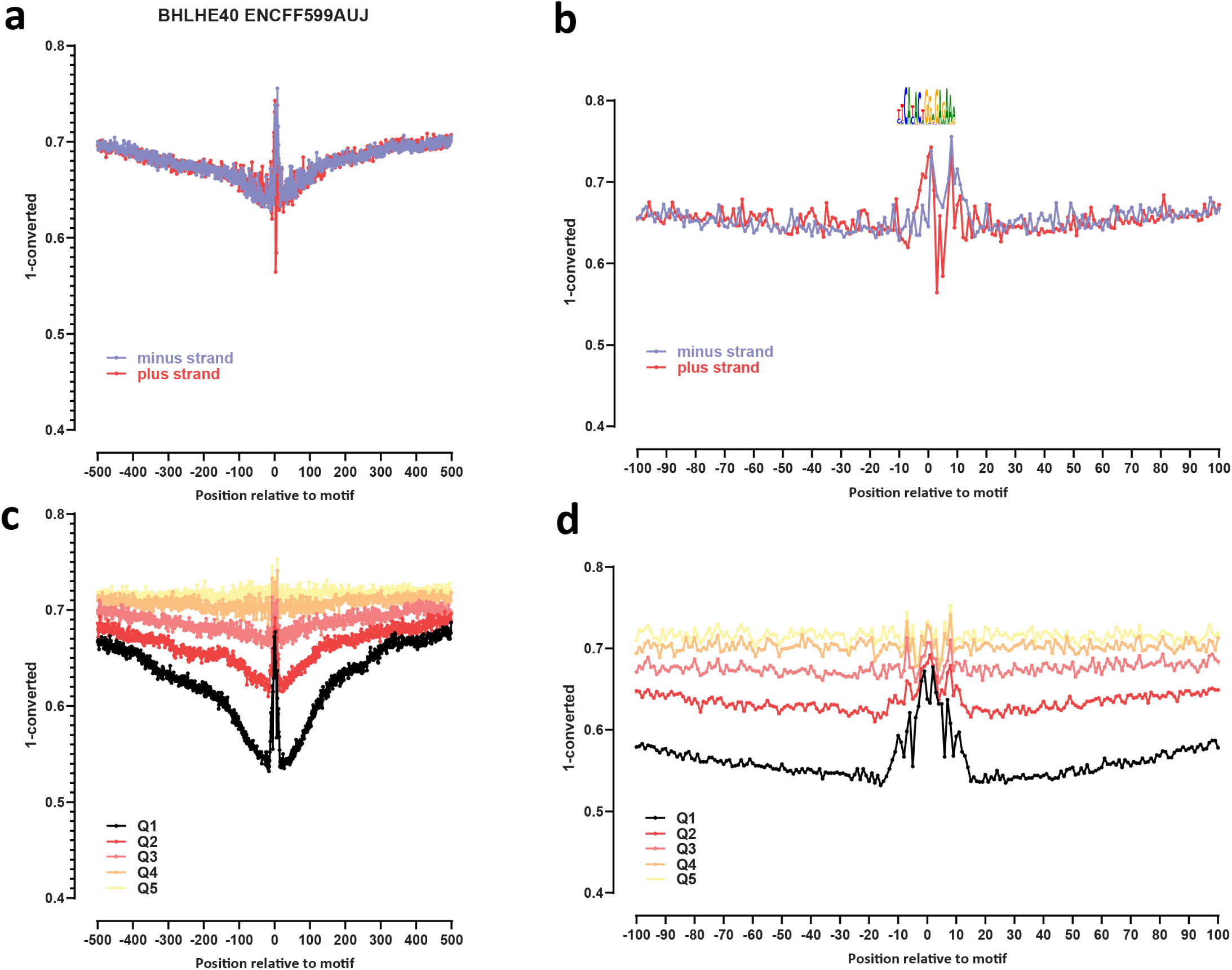
Genome-wide footprint profiles for the BHLHE40 transcription factor. GM12878 ssDNA-CseDa01-XL-SMF datasets are shown. (a) Genome-wide footprint profiles over the closest motif to ENCODE ChIP-seq peaks (ENCODE dataset ID indicated on top). Profiles are generated in a strand-aware way with respect to the motif orientation in the genome, and are shown separately for forward and reverse-strand reads. (b) Same as in (a) but zoomed in. The transcription factor sequence recognition motif is shown on top. (c) Genome-wide footprint profiles over the closest motif to ENCODE ChIP-seq peaks for the TF divided into quintiles based on ChIP-seq signal strength. (d) Same as in (c) but zoomed in.

**Supplementary Figure 13:**
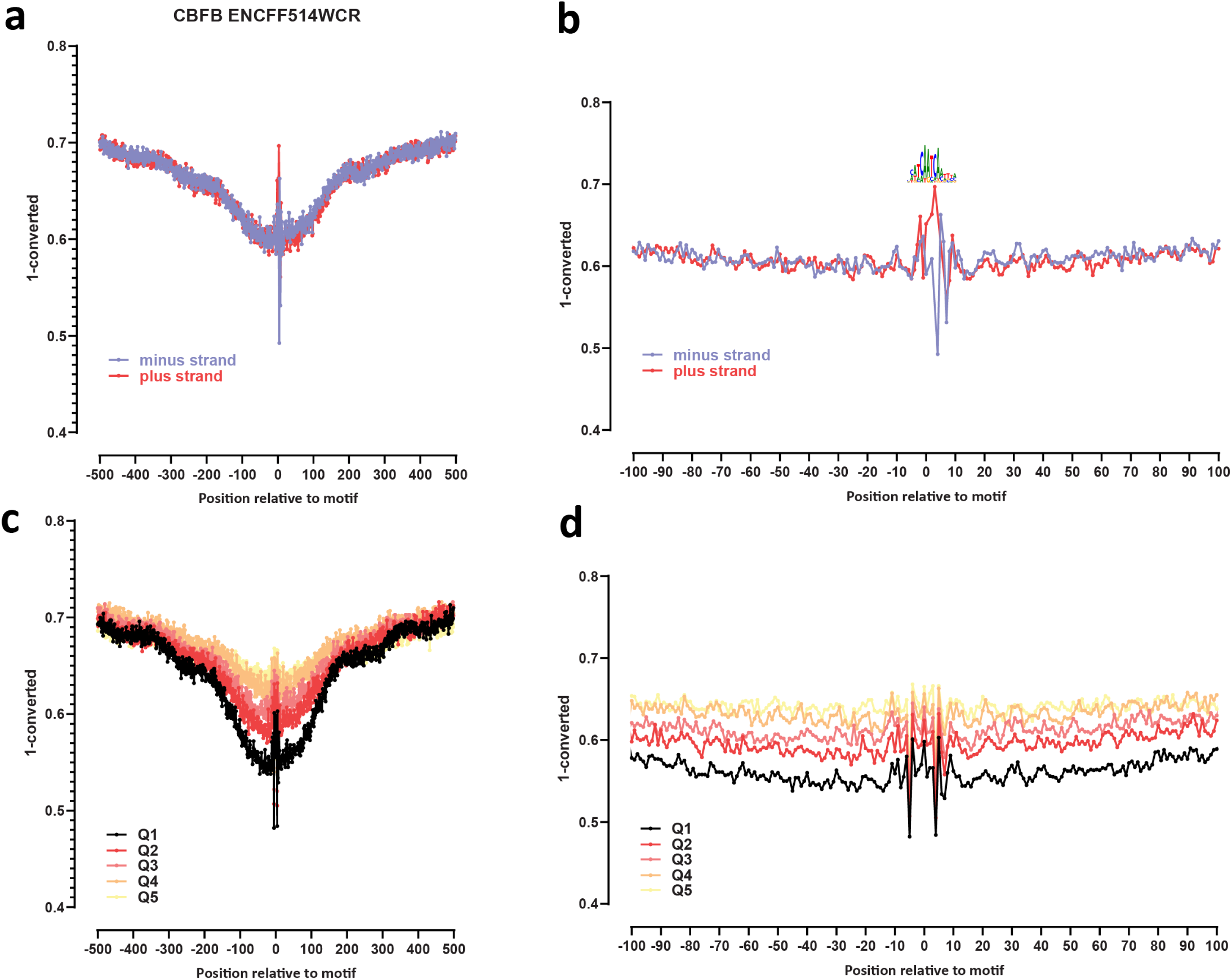
Genome-wide footprint profiles for the CBFB transcription factor. GM12878 ssDNA-CseDa01-XL-SMF datasets are shown. (a) Genome-wide footprint profiles over the closest motif to ENCODE ChIP-seq peaks (ENCODE dataset ID indicated on top). Profiles are generated in a strand-aware way with respect to the motif orientation in the genome, and are shown separately for forward and reverse-strand reads. (b) Same as in (a) but zoomed in. The transcription factor sequence recognition motif is shown on top. (c) Genome-wide footprint profiles over the closest motif to ENCODE ChIP-seq peaks for the TF divided into quintiles based on ChIP-seq signal strength. (d) Same as in (c) but zoomed in.

**Supplementary Figure 14:**
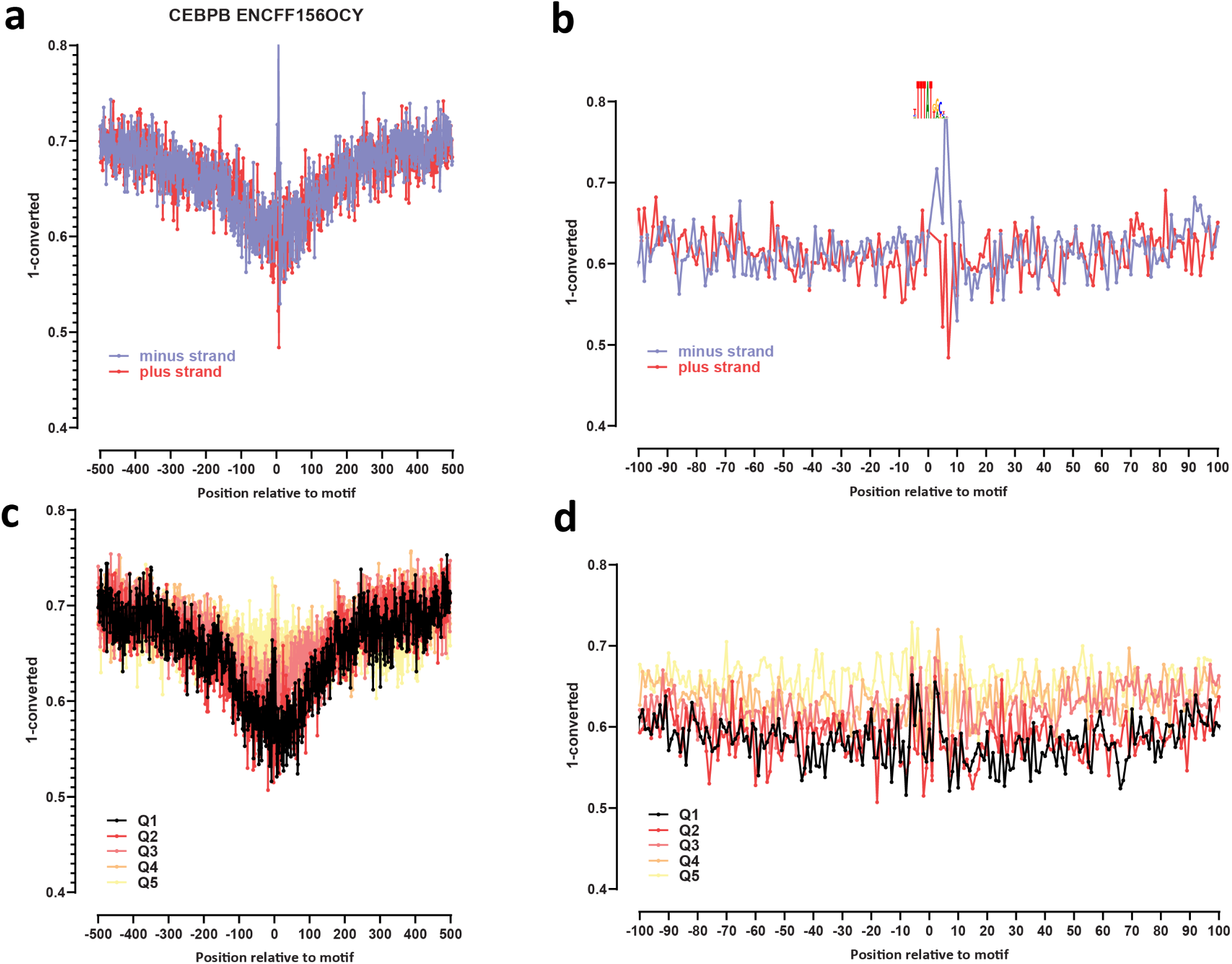
Genome-wide footprint profiles for the CEBPB transcription factor. GM12878 ssDNA-CseDa01-XL-SMF datasets are shown. (a) Genome-wide footprint profiles over the closest motif to ENCODE ChIP-seq peaks (ENCODE dataset ID indicated on top). Profiles are generated in a strand-aware way with respect to the motif orientation in the genome, and are shown separately for forward and reverse-strand reads. (b) Same as in (a) but zoomed in. The transcription factor sequence recognition motif is shown on top. (c) Genome-wide footprint profiles over the closest motif to ENCODE ChIP-seq peaks for the TF divided into quintiles based on ChIP-seq signal strength. (d) Same as in (c) but zoomed in.

**Supplementary Figure 15:**
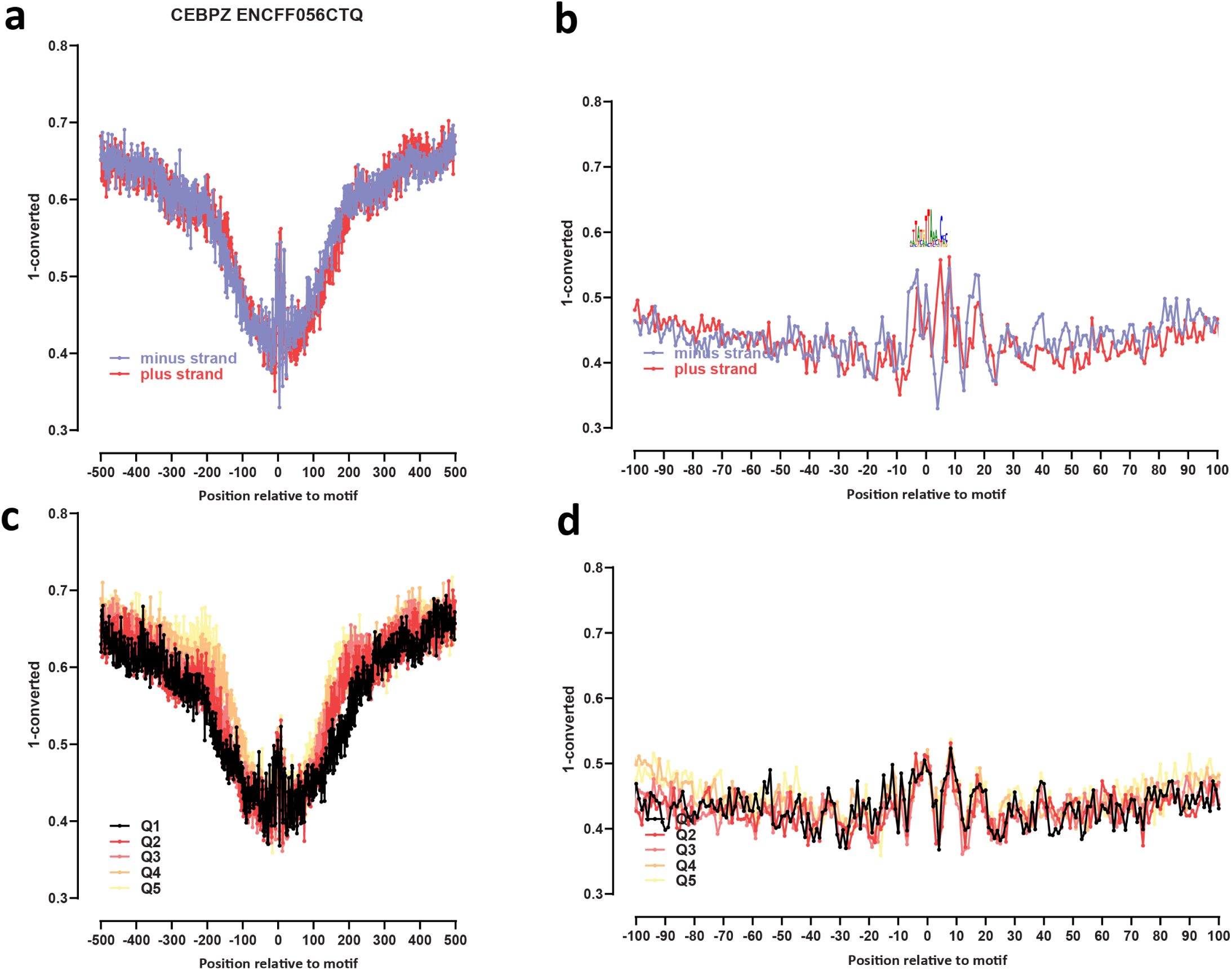
Genome-wide footprint profiles for the CEBPZ transcription factor. GM12878 ssDNA-CseDa01-XL-SMF datasets are shown. (a) Genome-wide footprint profiles over the closest motif to ENCODE ChIP-seq peaks (ENCODE dataset ID indicated on top). Profiles are generated in a strand-aware way with respect to the motif orientation in the genome, and are shown separately for forward and reverse-strand reads. (b) Same as in (a) but zoomed in. The transcription factor sequence recognition motif is shown on top. (c) Genome-wide footprint profiles over the closest motif to ENCODE ChIP-seq peaks for the TF divided into quintiles based on ChIP-seq signal strength. (d) Same as in (c) but zoomed in.

**Supplementary Figure 16:**
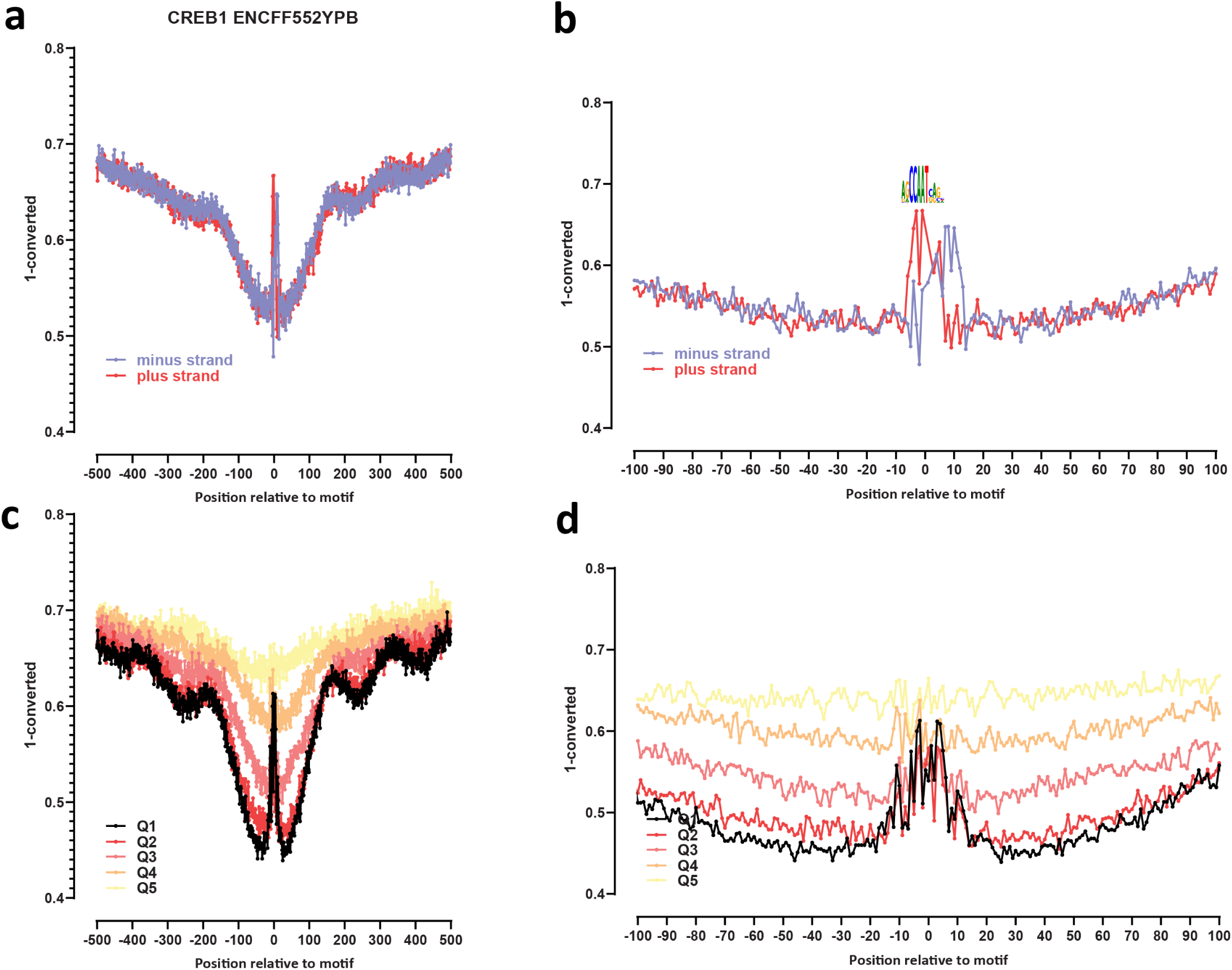
Genome-wide footprint profiles for the CREB1 transcription factor. GM12878 ssDNA-CseDa01-XL-SMF datasets are shown. (a) Genome-wide footprint profiles over the closest motif to ENCODE ChIP-seq peaks (ENCODE dataset ID indicated on top). Profiles are generated in a strand-aware way with respect to the motif orientation in the genome, and are shown separately for forward and reverse-strand reads. (b) Same as in (a) but zoomed in. The transcription factor sequence recognition motif is shown on top. (c) Genome-wide footprint profiles over the closest motif to ENCODE ChIP-seq peaks for the TF divided into quintiles based on ChIP-seq signal strength. (d) Same as in (c) but zoomed in.

**Supplementary Figure 17:**
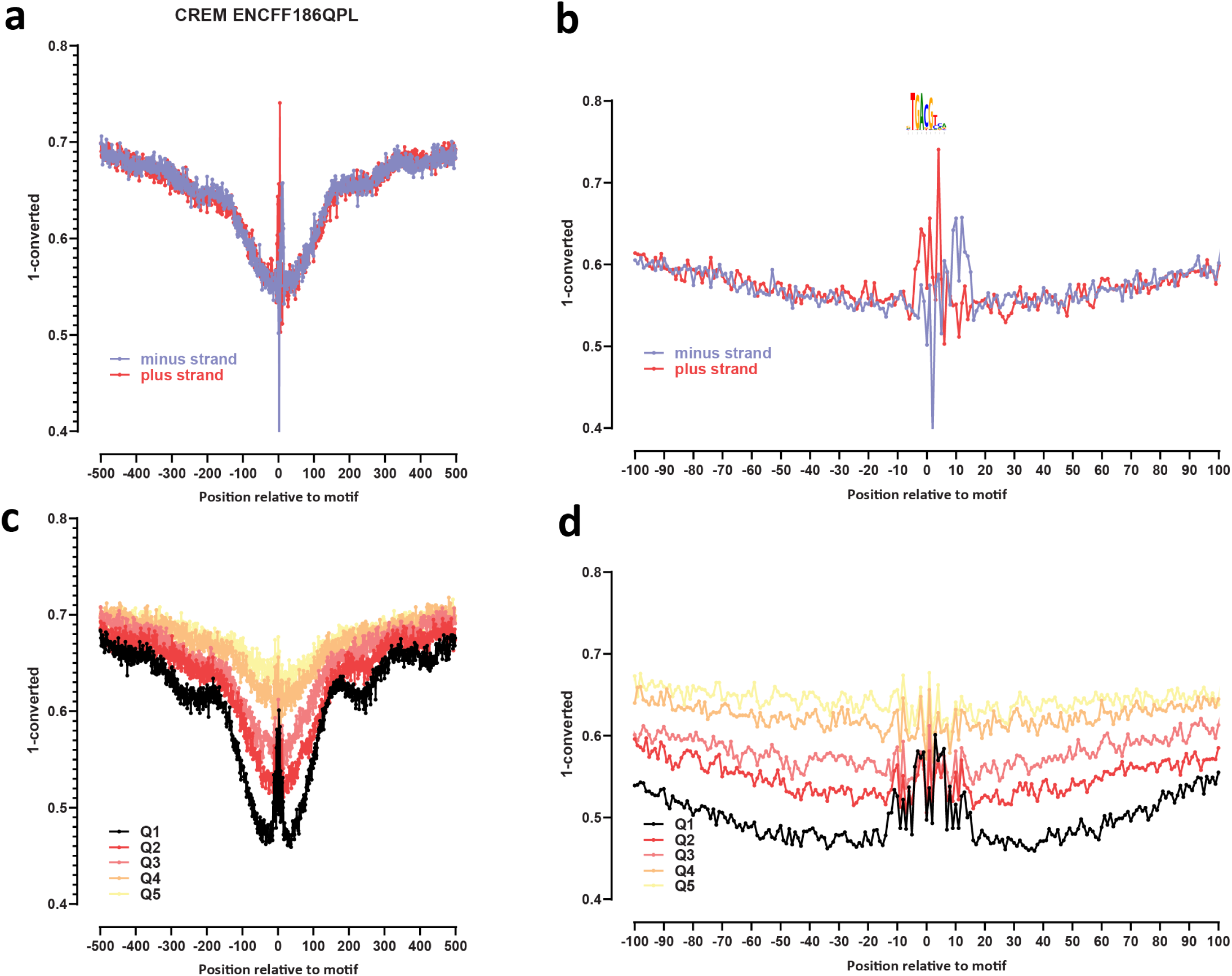
Genome-wide footprint profiles for the CREM transcription factor. GM12878 ssDNA-CseDa01-XL-SMF datasets are shown. (a) Genome-wide footprint profiles over the closest motif to ENCODE ChIP-seq peaks (ENCODE dataset ID indicated on top). Profiles are generated in a strand-aware way with respect to the motif orientation in the genome, and are shown separately for forward and reverse-strand reads. (b) Same as in (a) but zoomed in. The transcription factor sequence recognition motif is shown on top. (c) Genome-wide footprint profiles over the closest motif to ENCODE ChIP-seq peaks for the TF divided into quintiles based on ChIP-seq signal strength. (d) Same as in (c) but zoomed in.

**Supplementary Figure 18:**
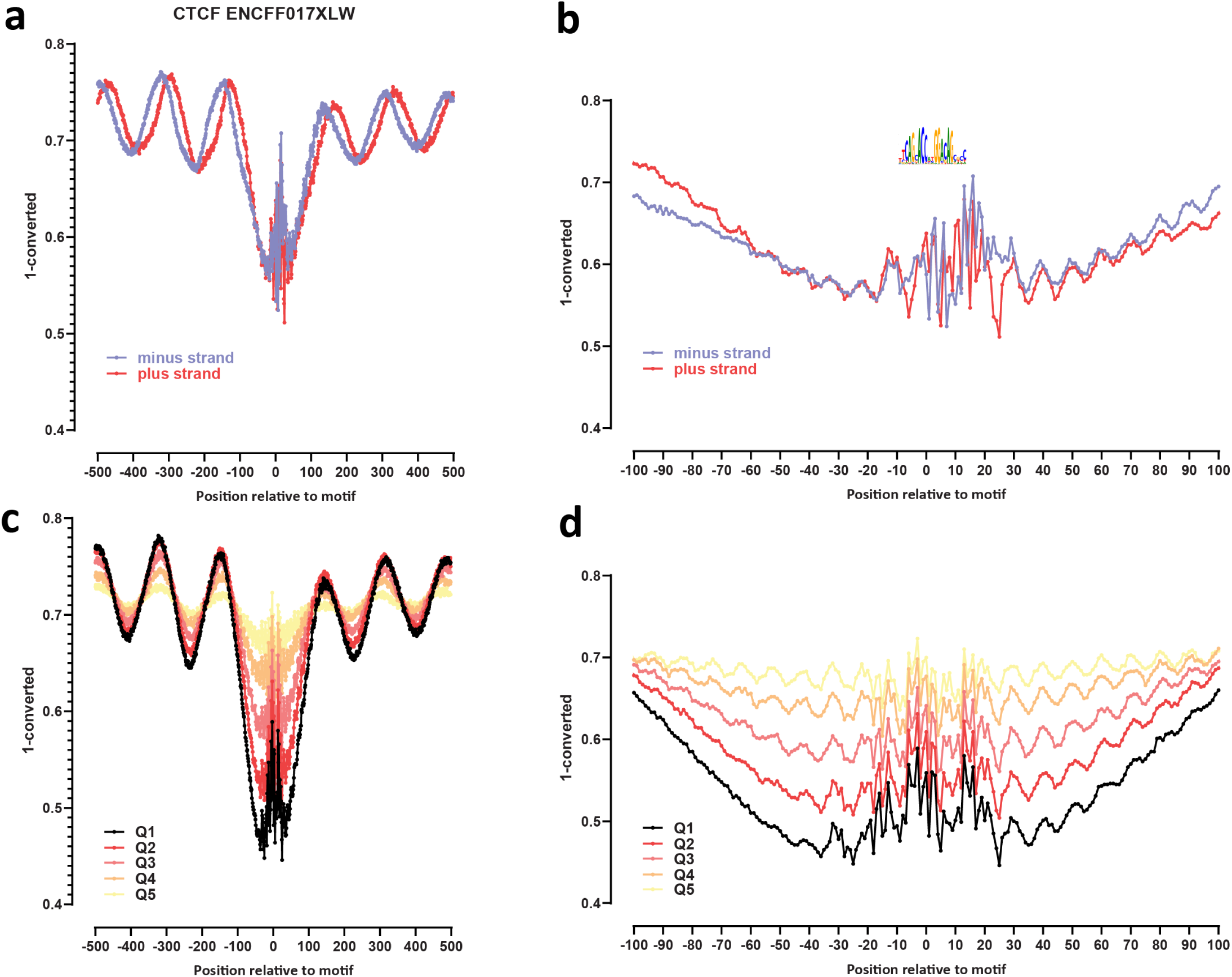
Genome-wide footprint profiles for the CTCF transcription factor. GM12878 ssDNA-CseDa01-XL-SMF datasets are shown. (a) Genome-wide footprint profiles over the closest motif to ENCODE ChIP-seq peaks (ENCODE dataset ID indicated on top). Profiles are generated in a strand-aware way with respect to the motif orientation in the genome, and are shown separately for forward and reverse-strand reads. (b) Same as in (a) but zoomed in. The transcription factor sequence recognition motif is shown on top. (c) Genome-wide footprint profiles over the closest motif to ENCODE ChIP-seq peaks for the TF divided into quintiles based on ChIP-seq signal strength. (d) Same as in (c) but zoomed in.

**Supplementary Figure 19:**
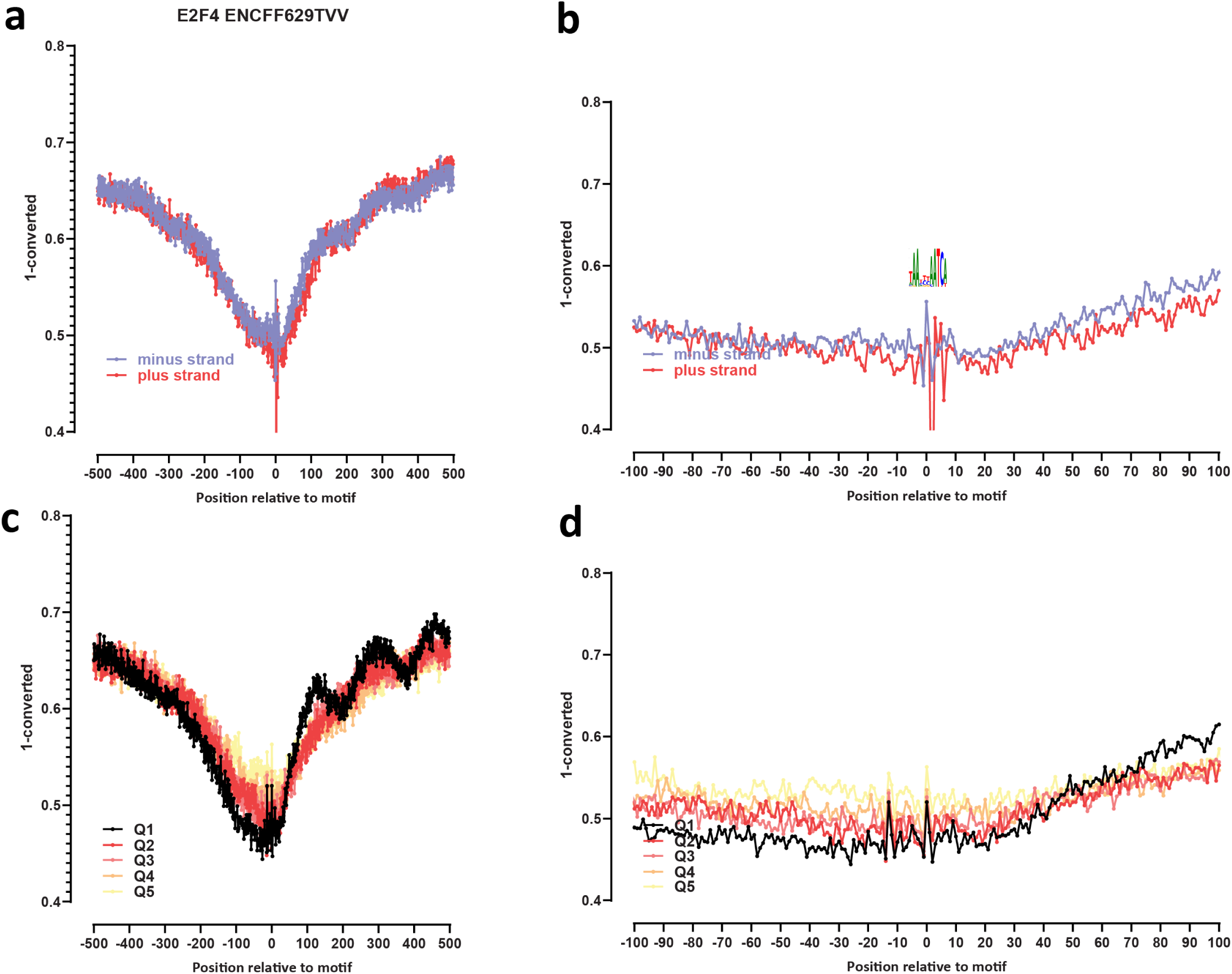
Genome-wide footprint profiles for the E2F4 transcription factor. GM12878 ssDNA-CseDa01-XL-SMF datasets are shown. (a) Genome-wide footprint profiles over the closest motif to ENCODE ChIP-seq peaks (ENCODE dataset ID indicated on top). Profiles are generated in a strand-aware way with respect to the motif orientation in the genome, and are shown separately for forward and reverse-strand reads. (b) Same as in (a) but zoomed in. The transcription factor sequence recognition motif is shown on top. (c) Genome-wide footprint profiles over the closest motif to ENCODE ChIP-seq peaks for the TF divided into quintiles based on ChIP-seq signal strength. (d) Same as in (c) but zoomed in.

**Supplementary Figure 20:**
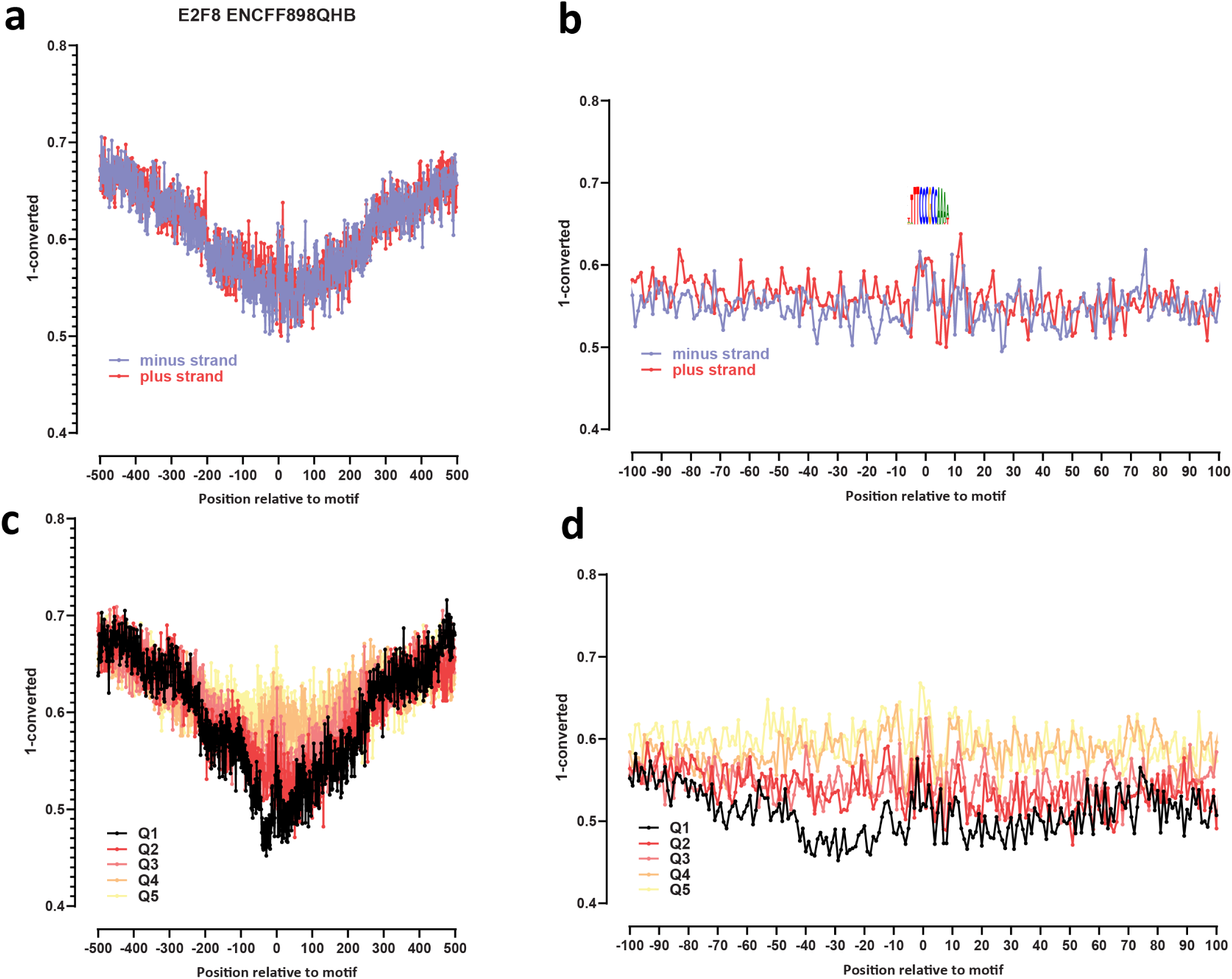
Genome-wide footprint profiles for the E2F8 transcription factor. GM12878 ssDNA-CseDa01-XL-SMF datasets are shown. (a) Genome-wide footprint profiles over the closest motif to ENCODE ChIP-seq peaks (ENCODE dataset ID indicated on top). Profiles are generated in a strand-aware way with respect to the motif orientation in the genome, and are shown separately for forward and reverse-strand reads. (b) Same as in (a) but zoomed in. The transcription factor sequence recognition motif is shown on top. (c) Genome-wide footprint profiles over the closest motif to ENCODE ChIP-seq peaks for the TF divided into quintiles based on ChIP-seq signal strength. (d) Same as in (c) but zoomed in.

**Supplementary Figure 21:**
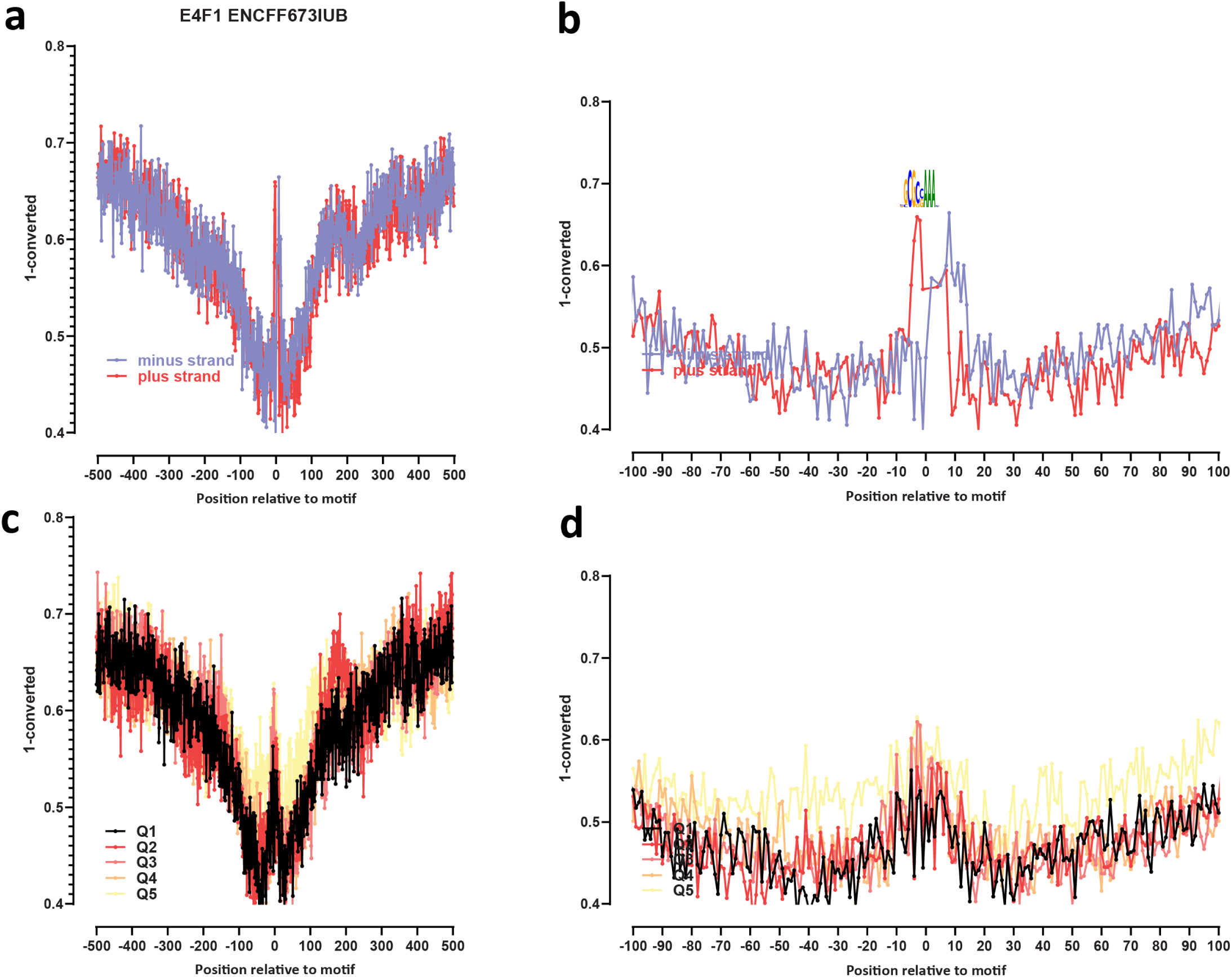
Genome-wide footprint profiles for the E4F1 transcription factor. GM12878 ssDNA-CseDa01-XL-SMF datasets are shown. (a) Genome-wide footprint profiles over the closest motif to ENCODE ChIP-seq peaks (ENCODE dataset ID indicated on top). Profiles are generated in a strand-aware way with respect to the motif orientation in the genome, and are shown separately for forward and reverse-strand reads. (b) Same as in (a) but zoomed in. The transcription factor sequence recognition motif is shown on top. (c) Genome-wide footprint profiles over the closest motif to ENCODE ChIP-seq peaks for the TF divided into quintiles based on ChIP-seq signal strength. (d) Same as in (c) but zoomed in.

**Supplementary Figure 22:**
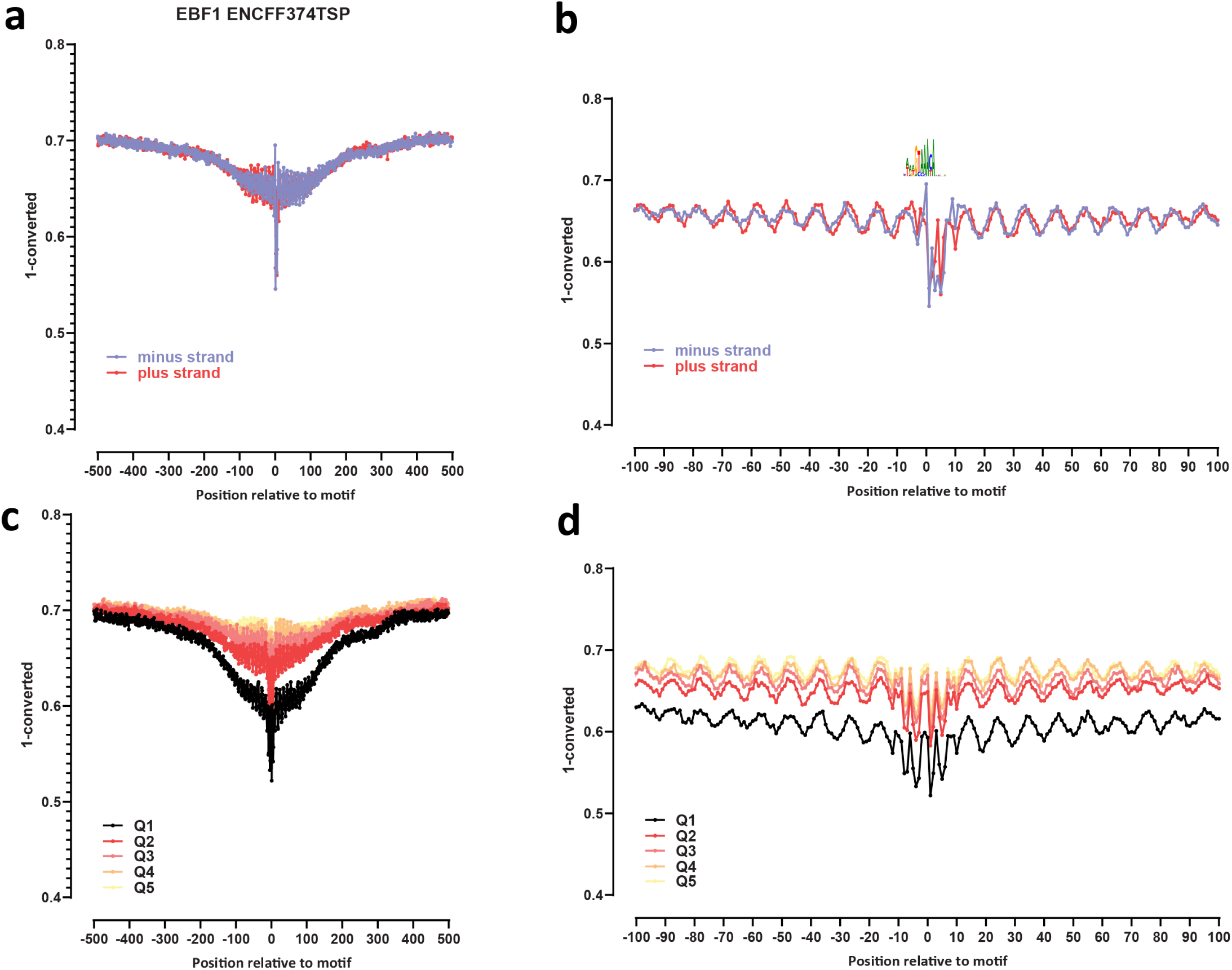
Genome-wide footprint profiles for the EBF1 transcription factor. GM12878 ssDNA-CseDa01-XL-SMF datasets are shown. (a) Genome-wide footprint profiles over the closest motif to ENCODE ChIP-seq peaks (ENCODE dataset ID indicated on top). Profiles are generated in a strand-aware way with respect to the motif orientation in the genome, and are shown separately for forward and reverse-strand reads. (b) Same as in (a) but zoomed in. The transcription factor sequence recognition motif is shown on top. (c) Genome-wide footprint profiles over the closest motif to ENCODE ChIP-seq peaks for the TF divided into quintiles based on ChIP-seq signal strength. (d) Same as in (c) but zoomed in.

**Supplementary Figure 23:**
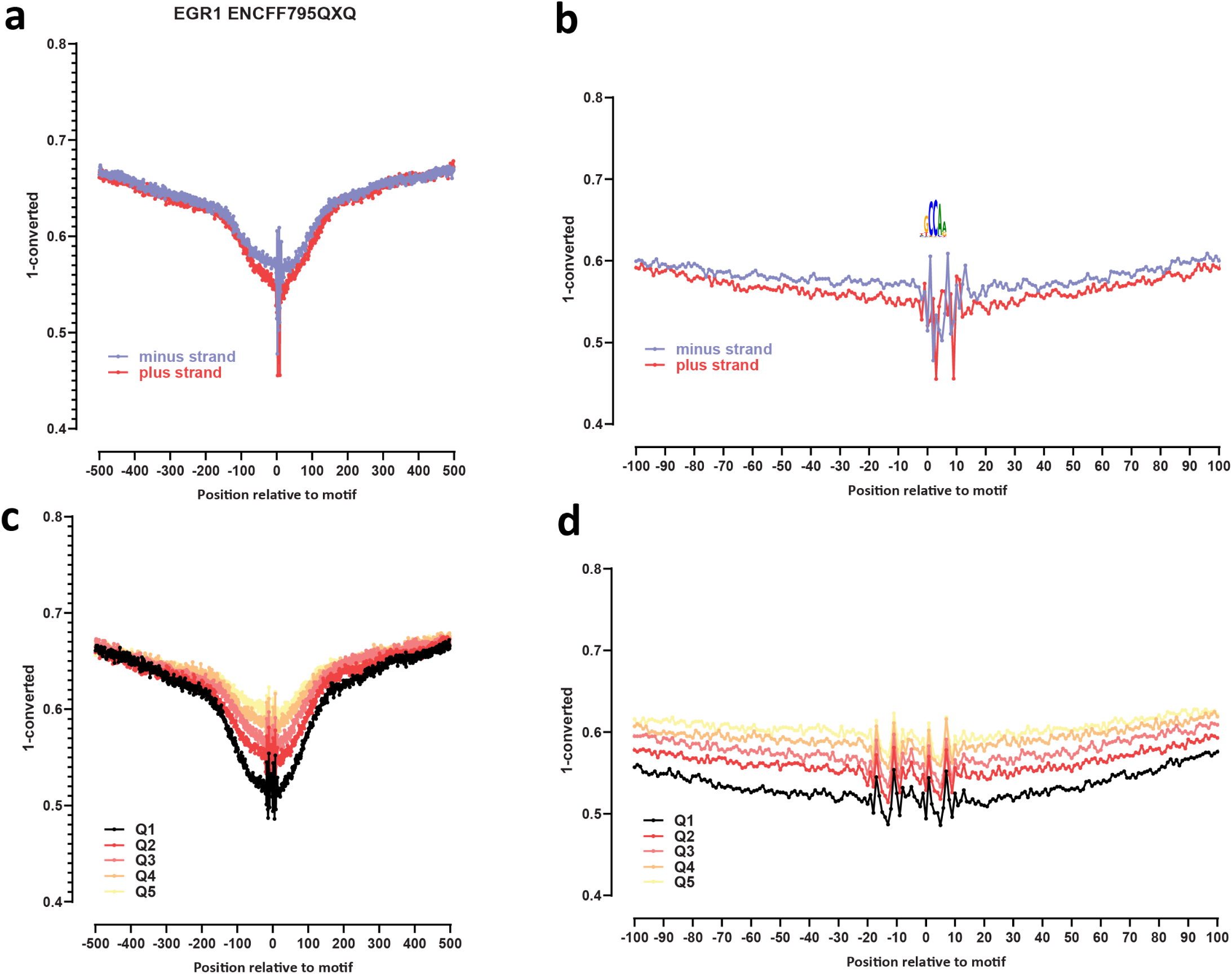
Genome-wide footprint profiles for the EGR1 transcription factor. GM12878 ssDNA-CseDa01-XL-SMF datasets are shown. (a) Genome-wide footprint profiles over the closest motif to ENCODE ChIP-seq peaks (ENCODE dataset ID indicated on top). Profiles are generated in a strand-aware way with respect to the motif orientation in the genome, and are shown separately for forward and reverse-strand reads. (b) Same as in (a) but zoomed in. The transcription factor sequence recognition motif is shown on top. (c) Genome-wide footprint profiles over the closest motif to ENCODE ChIP-seq peaks for the TF divided into quintiles based on ChIP-seq signal strength. (d) Same as in (c) but zoomed in.

**Supplementary Figure 24:**
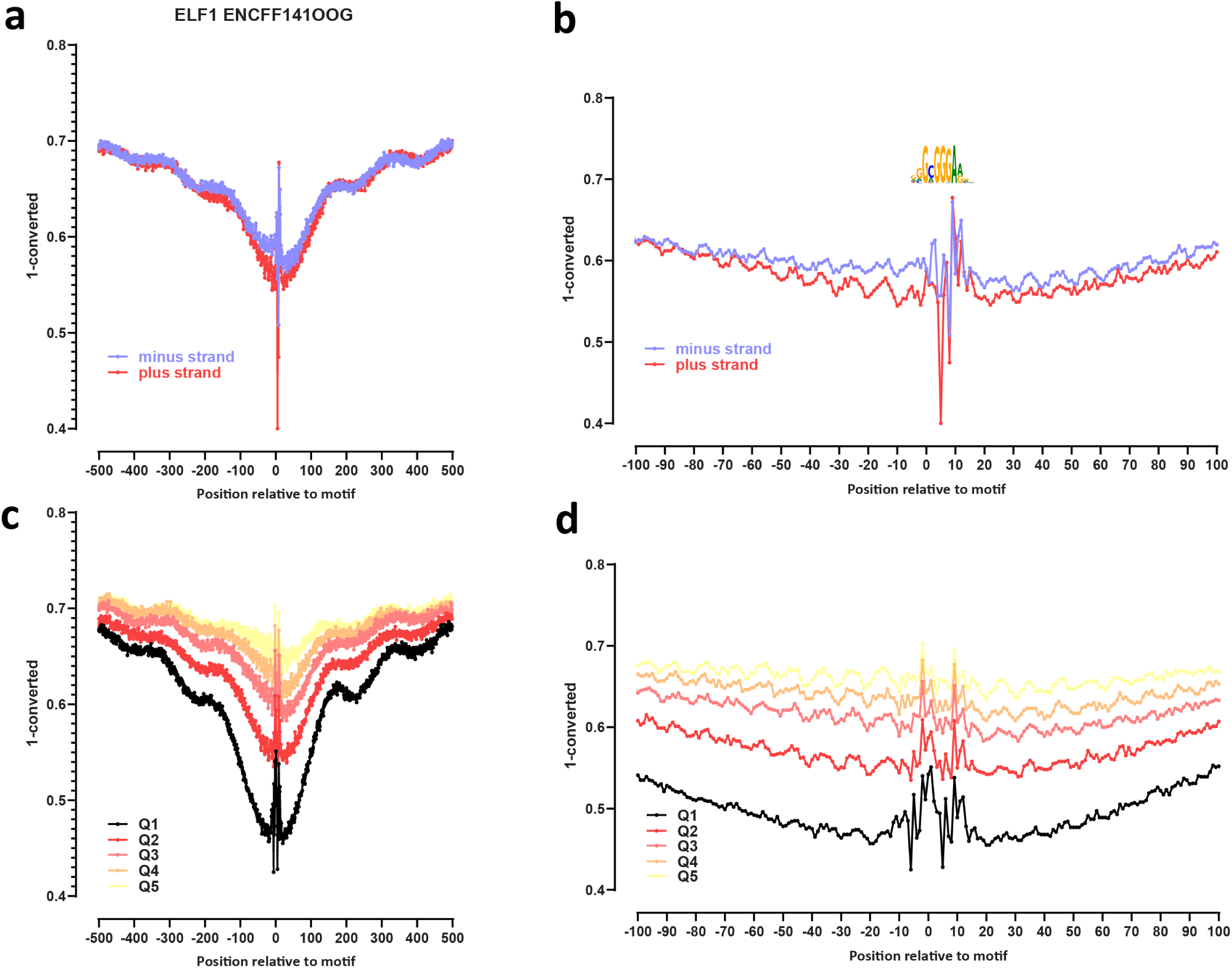
Genome-wide footprint profiles for the ELF1 transcription factor. GM12878 ssDNA-CseDa01-XL-SMF datasets are shown. (a) Genome-wide footprint profiles over the closest motif to ENCODE ChIP-seq peaks (ENCODE dataset ID indicated on top). Profiles are generated in a strand-aware way with respect to the motif orientation in the genome, and are shown separately for forward and reverse-strand reads. (b) Same as in (a) but zoomed in. The transcription factor sequence recognition motif is shown on top. (c) Genome-wide footprint profiles over the closest motif to ENCODE ChIP-seq peaks for the TF divided into quintiles based on ChIP-seq signal strength. (d) Same as in (c) but zoomed in.

**Supplementary Figure 25:**
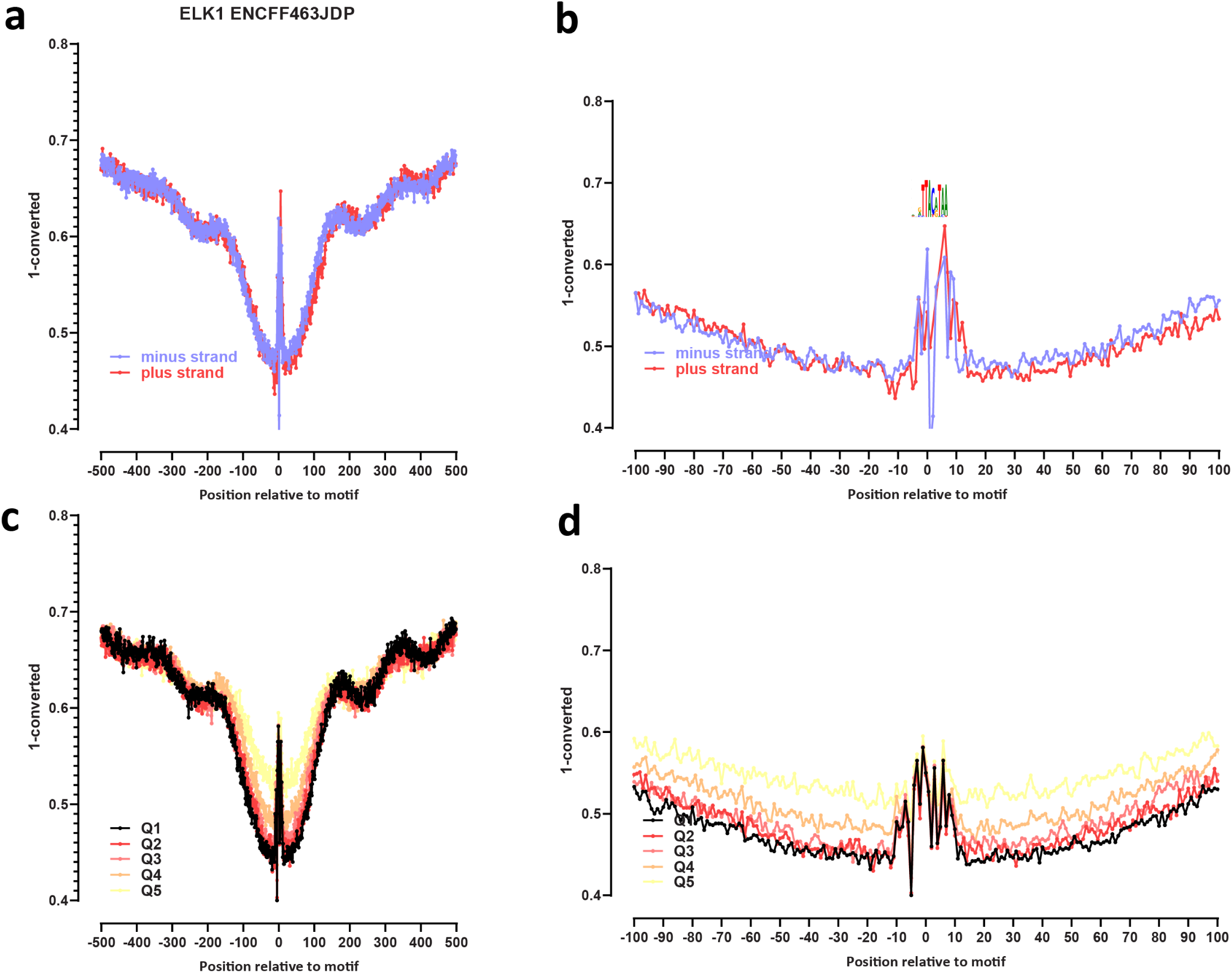
Genome-wide footprint profiles for the ELK1 transcription factor. GM12878 ssDNA-CseDa01-XL-SMF datasets are shown. (a) Genome-wide footprint profiles over the closest motif to ENCODE ChIP-seq peaks (ENCODE dataset ID indicated on top). Profiles are generated in a strand-aware way with respect to the motif orientation in the genome, and are shown separately for forward and reverse-strand reads. (b) Same as in (a) but zoomed in. The transcription factor sequence recognition motif is shown on top. (c) Genome-wide footprint profiles over the closest motif to ENCODE ChIP-seq peaks for the TF divided into quintiles based on ChIP-seq signal strength. (d) Same as in (c) but zoomed in.

**Supplementary Figure 26:**
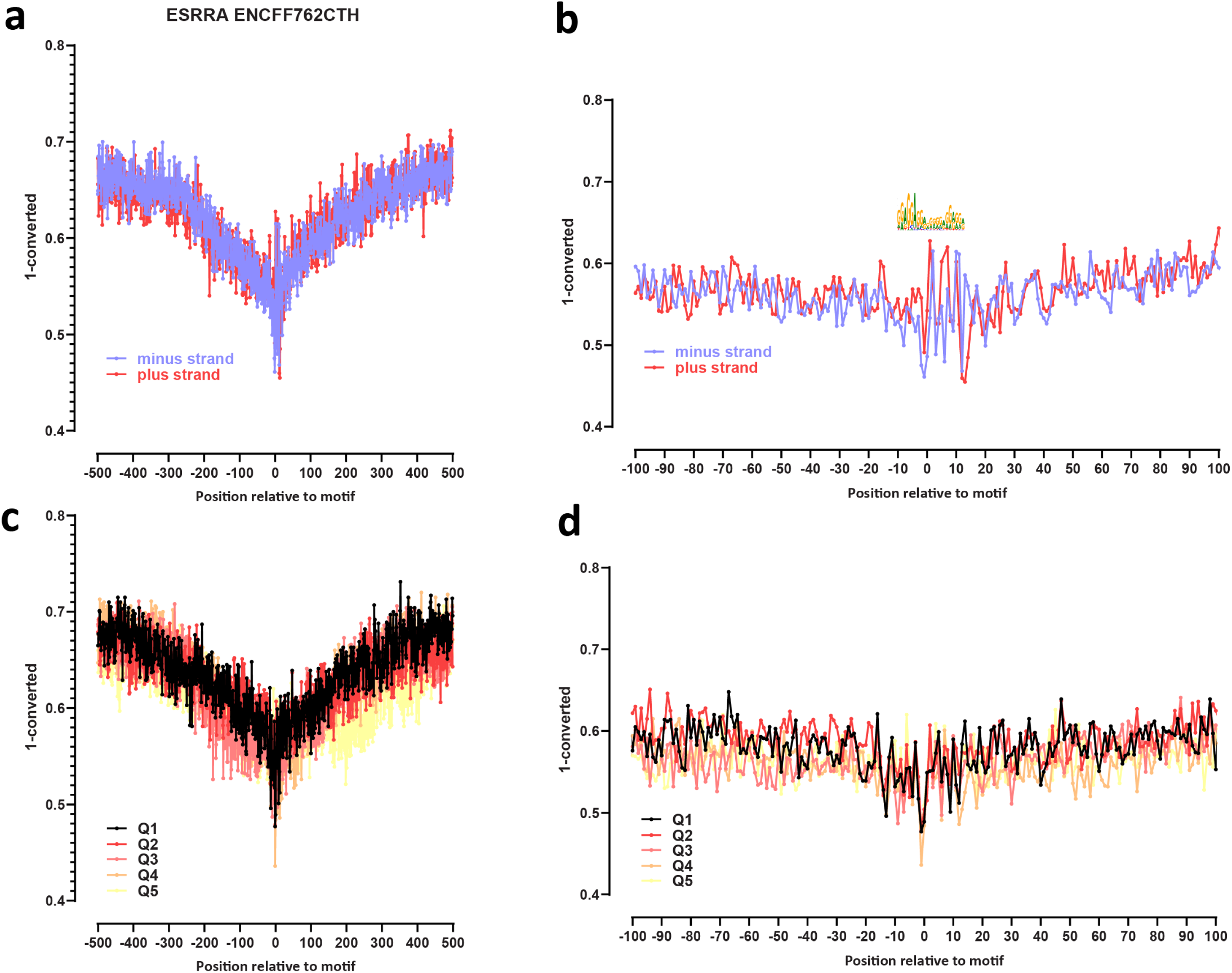
Genome-wide footprint profiles for the ESRRA transcription factor. GM12878 ssDNA-CseDa01-XL-SMF datasets are shown. (a) Genome-wide footprint profiles over the closest motif to ENCODE ChIP-seq peaks (ENCODE dataset ID indicated on top). Profiles are generated in a strand-aware way with respect to the motif orientation in the genome, and are shown separately for forward and reverse-strand reads. (b) Same as in (a) but zoomed in. The transcription factor sequence recognition motif is shown on top. (c) Genome-wide footprint profiles over the closest motif to ENCODE ChIP-seq peaks for the TF divided into quintiles based on ChIP-seq signal strength. (d) Same as in (c) but zoomed in.

**Supplementary Figure 27:**
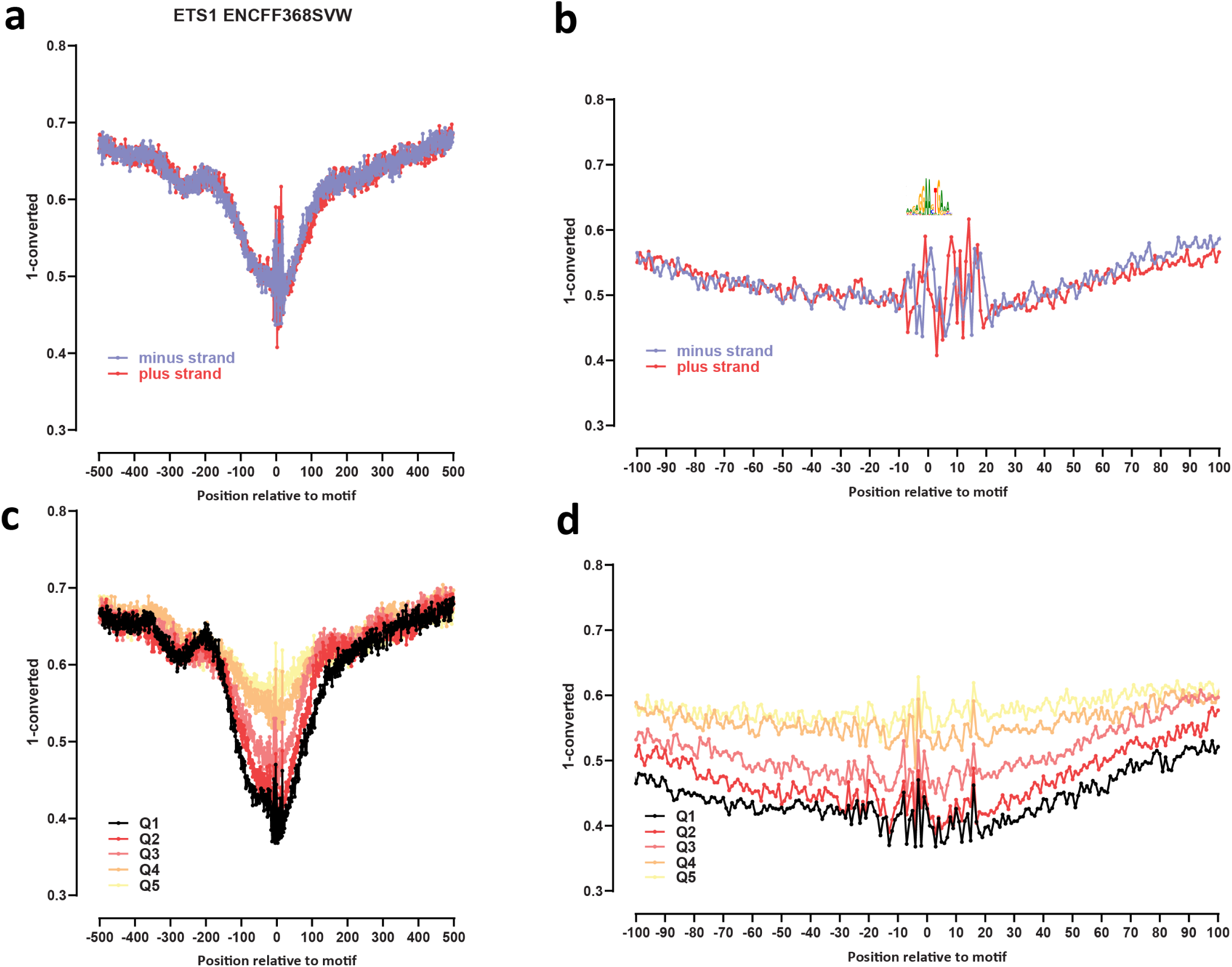
Genome-wide footprint profiles for the ETS1 transcription factor. GM12878 ssDNA-CseDa01-XL-SMF datasets are shown. (a) Genome-wide footprint profiles over the closest motif to ENCODE ChIP-seq peaks (ENCODE dataset ID indicated on top). Profiles are generated in a strand-aware way with respect to the motif orientation in the genome, and are shown separately for forward and reverse-strand reads. (b) Same as in (a) but zoomed in. The transcription factor sequence recognition motif is shown on top. (c) Genome-wide footprint profiles over the closest motif to ENCODE ChIP-seq peaks for the TF divided into quintiles based on ChIP-seq signal strength.(d) Same as in (c) but zoomed in.

**Supplementary Figure 28:**
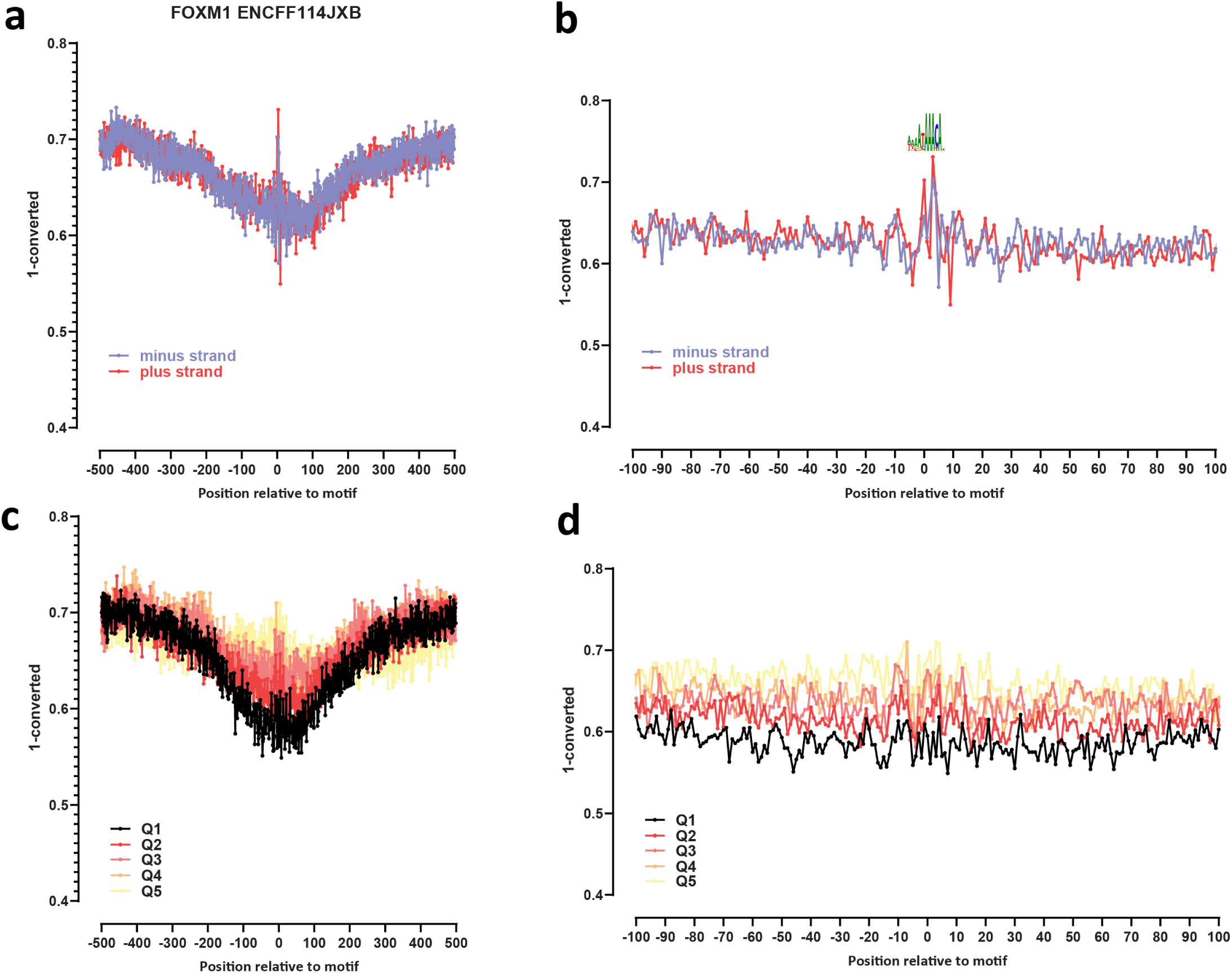
Genome-wide footprint profiles for the FOXM1 transcription factor. GM12878 ssDNA-CseDa01-XL-SMF datasets are shown. (a) Genome-wide footprint profiles over the closest motif to ENCODE ChIP-seq peaks (ENCODE dataset ID indicated on top). Profiles are generated in a strand-aware way with respect to the motif orientation in the genome, and are shown separately for forward and reverse-strand reads. (b) Same as in (a) but zoomed in. The transcription factor sequence recognition motif is shown on top. (c) Genome-wide footprint profiles over the closest motif to ENCODE ChIP-seq peaks for the TF divided into quintiles based on ChIP-seq signal strength. (d) Same as in (c) but zoomed in.

**Supplementary Figure 29:**
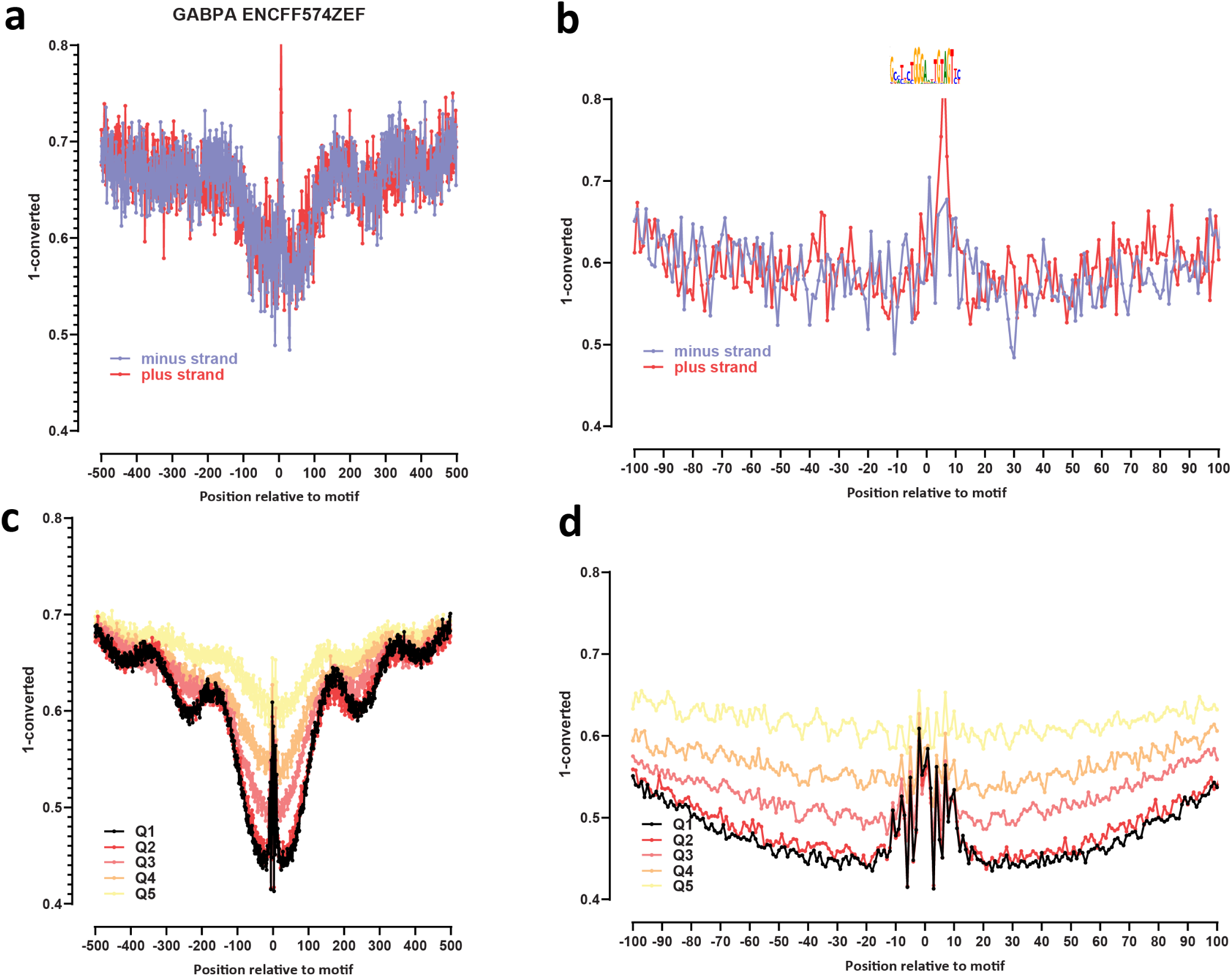
Genome-wide footprint profiles for the GABPA transcription factor. GM12878 ssDNA-CseDa01-XL-SMF datasets are shown. (a) Genome-wide footprint profiles over the closest motif to ENCODE ChIP-seq peaks (ENCODE dataset ID indicated on top). Profiles are generated in a strand-aware way with respect to the motif orientation in the genome, and are shown separately for forward and reverse-strand reads. (b) Same as in (a) but zoomed in. The transcription factor sequence recognition motif is shown on top. (c) Genome-wide footprint profiles over the closest motif to ENCODE ChIP-seq peaks for the TF divided into quintiles based on ChIP-seq signal strength. (d) Same as in (c) but zoomed in.

**Supplementary Figure 30:**
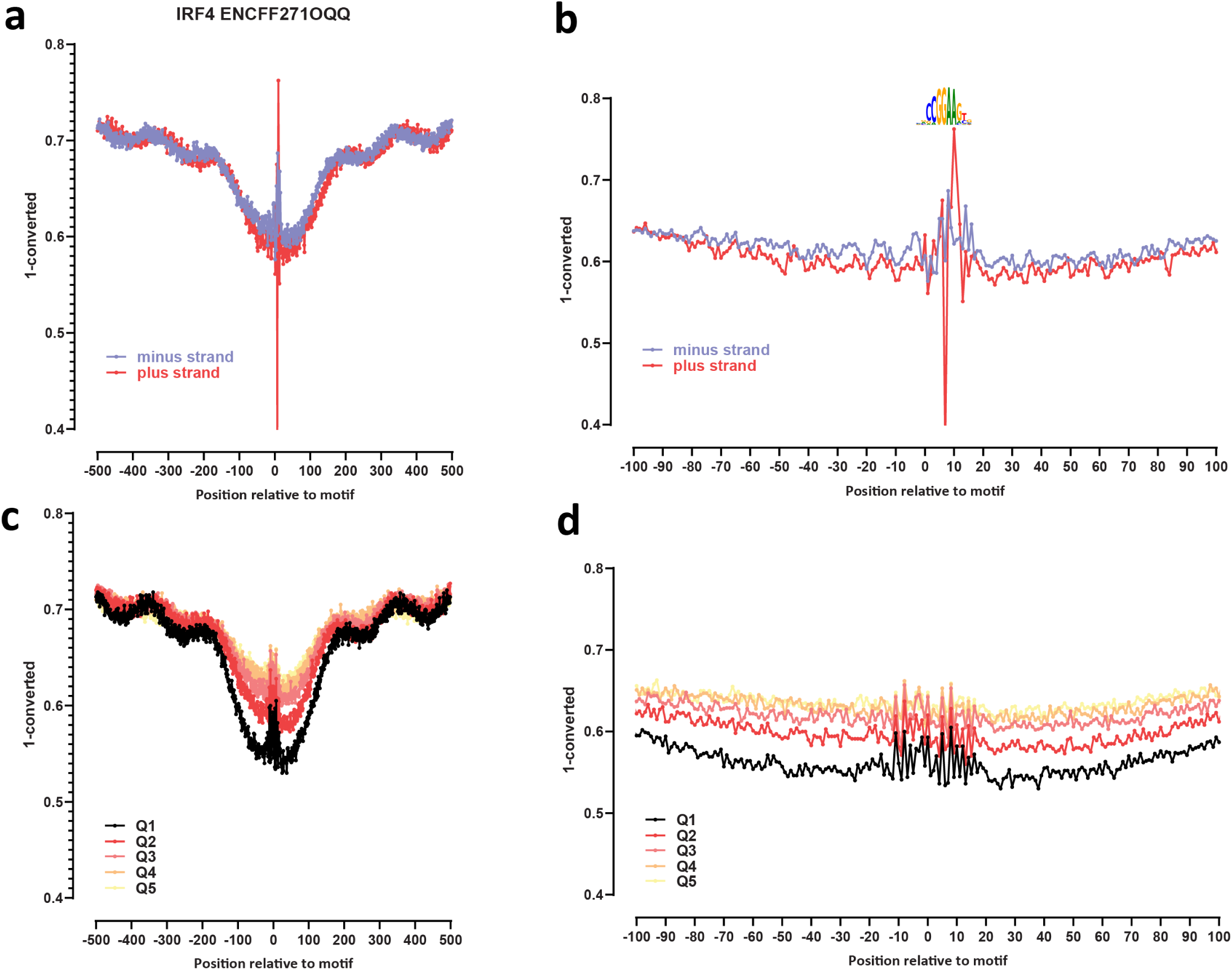
Genome-wide footprint profiles for the IRF4 transcription factor. GM12878 ssDNA-CseDa01-XL-SMF datasets are shown. (a) Genome-wide footprint profiles over the closest motif to ENCODE ChIP-seq peaks (ENCODE dataset ID indicated on top). Profiles are generated in a strand-aware way with respect to the motif orientation in the genome, and are shown separately for forward and reverse-strand reads. (b) Same as in (a) but zoomed in. The transcription factor sequence recognition motif is shown on top. (c) Genome-wide footprint profiles over the closest motif to ENCODE ChIP-seq peaks for the TF divided into quintiles based on ChIP-seq signal strength. (d) Same as in (c) but zoomed in.

**Supplementary Figure 31:**
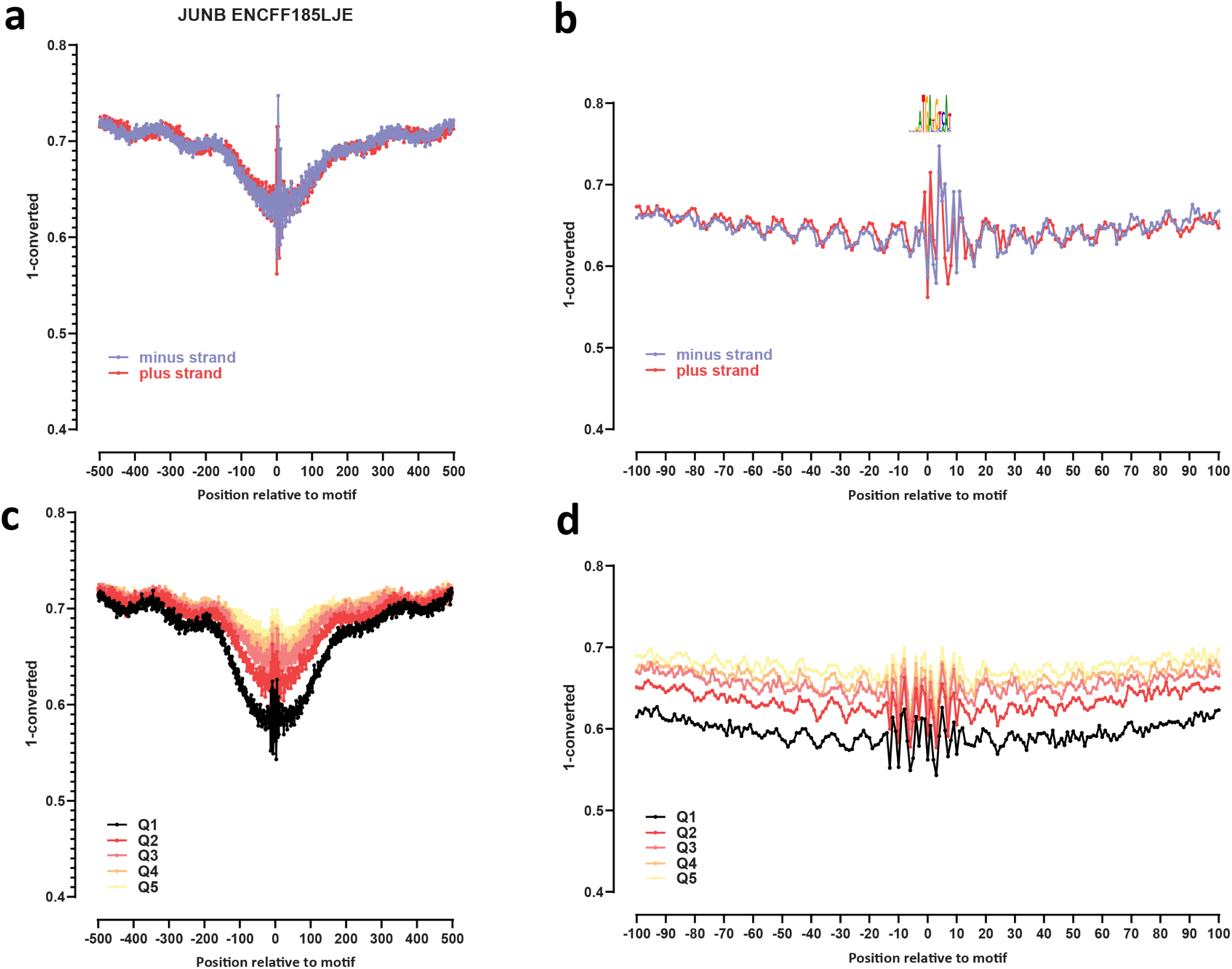
Genome-wide footprint profiles for the JUNB transcription factor. GM12878 ssDNA-CseDa01-XL-SMF datasets are shown. (a) Genome-wide footprint profiles over the closest motif to ENCODE ChIP-seq peaks (ENCODE dataset ID indicated on top). Profiles are generated in a strand-aware way with respect to the motif orientation in the genome, and are shown separately for forward and reverse-strand reads. (b) Same as in (a) but zoomed in. The transcription factor sequence recognition motif is shown on top. (c) Genome-wide footprint profiles over the closest motif to ENCODE ChIP-seq peaks for the TF divided into quintiles based on ChIP-seq signal strength. (d) Same as in (c) but zoomed in.

**Supplementary Figure 32:**
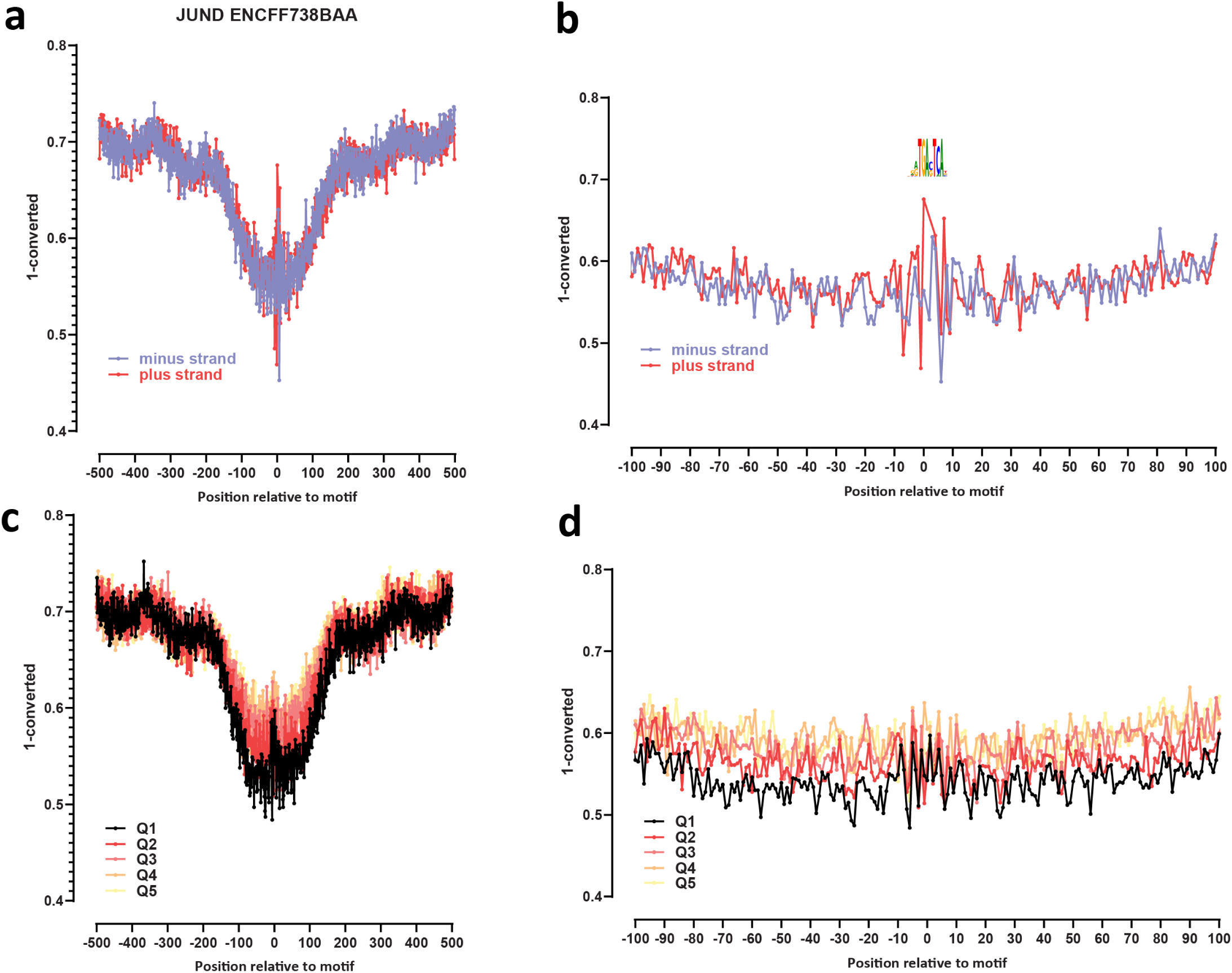
Genome-wide footprint profiles for the JUND transcription factor. GM12878 ssDNA-CseDa01-XL-SMF datasets are shown. (a) Genome-wide footprint profiles over the closest motif to ENCODE ChIP-seq peaks (ENCODE dataset ID indicated on top). Profiles are generated in a strand-aware way with respect to the motif orientation in the genome, and are shown separately for forward and reverse-strand reads. (b) Same as in (a) but zoomed in. The transcription factor sequence recognition motif is shown on top. (c) Genome-wide footprint profiles over the closest motif to ENCODE ChIP-seq peaks for the TF divided into quintiles based on ChIP-seq signal strength. (d) Same as in (c) but zoomed in.

**Supplementary Figure 33:**
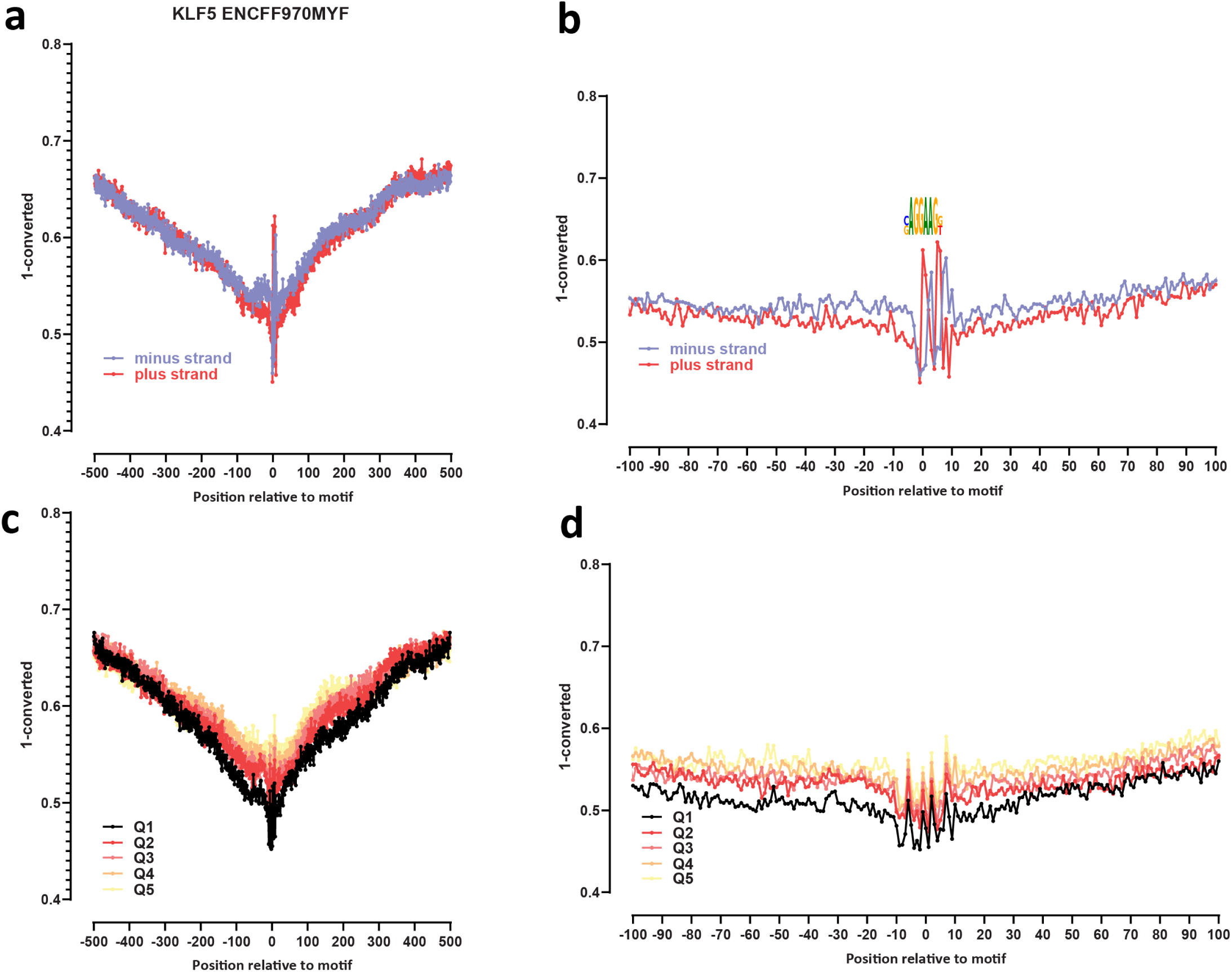
Genome-wide footprint profiles for the KLF5 transcription factor. GM12878 ssDNA-CseDa01-XL-SMF datasets are shown. (a) Genome-wide footprint profiles over the closest motif to ENCODE ChIP-seq peaks (ENCODE dataset ID indicated on top). Profiles are generated in a strand-aware way with respect to the motif orientation in the genome, and are shown separately for forward and reverse-strand reads. (b) Same as in (a) but zoomed in. The transcription factor sequence recognition motif is shown on top. (c) Genome-wide footprint profiles over the closest motif to ENCODE ChIP-seq peaks for the TF divided into quintiles based on ChIP-seq signal strength. (d) Same as in (c) but zoomed in.

**Supplementary Figure 34:**
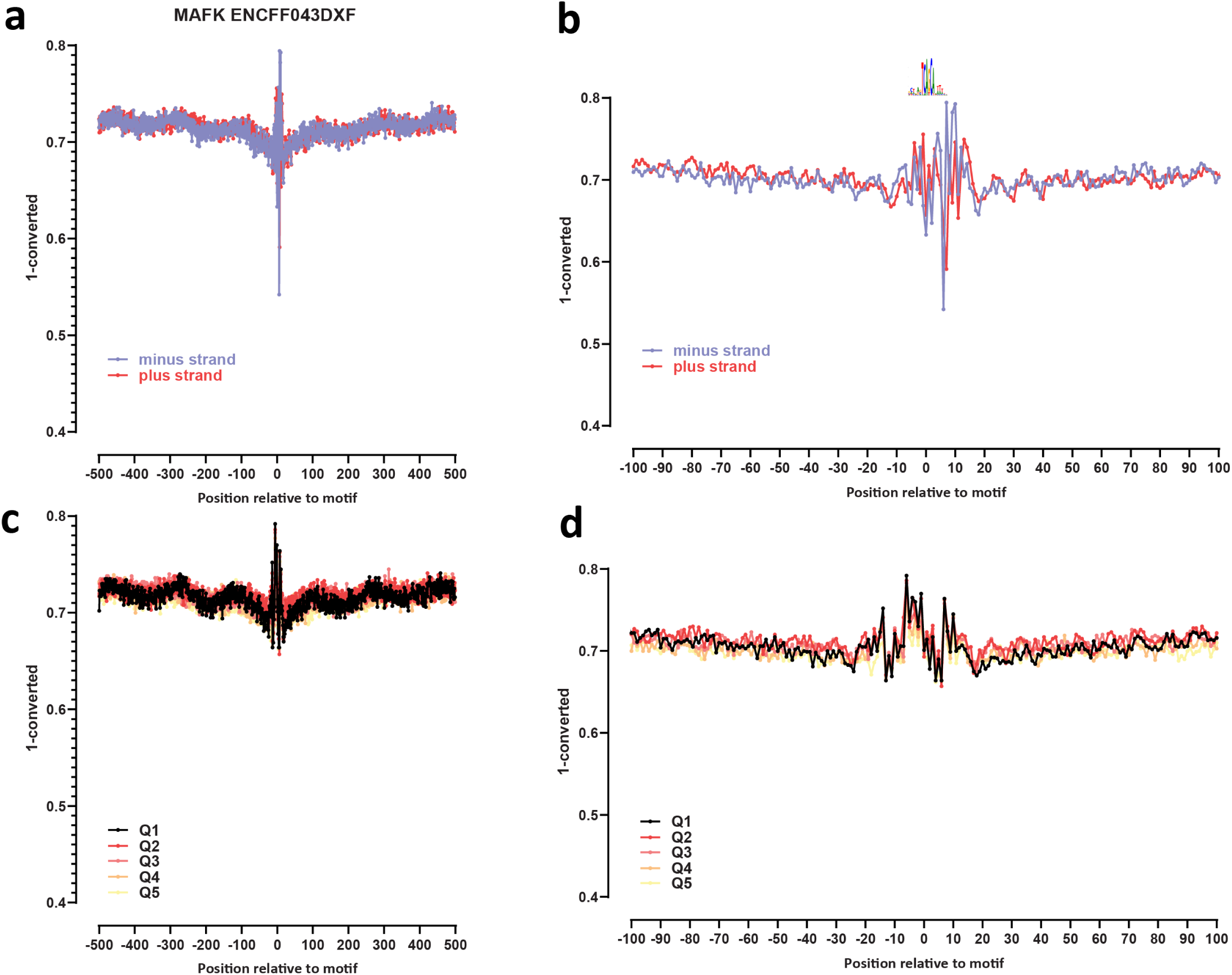
Genome-wide footprint profiles for the MAFK transcription factor. GM12878 ssDNA-CseDa01-XL-SMF datasets are shown. (a) Genome-wide footprint profiles over the closest motif to ENCODE ChIP-seq peaks (ENCODE dataset ID indicated on top). Profiles are generated in a strand-aware way with respect to the motif orientation in the genome, and are shown separately for forward and reverse-strand reads. (b) Same as in (a) but zoomed in. The transcription factor sequence recognition motif is shown on top. (c) Genome-wide footprint profiles over the closest motif to ENCODE ChIP-seq peaks for the TF divided into quintiles based on ChIP-seq signal strength. (d) Same as in (c) but zoomed in.

**Supplementary Figure 35:**
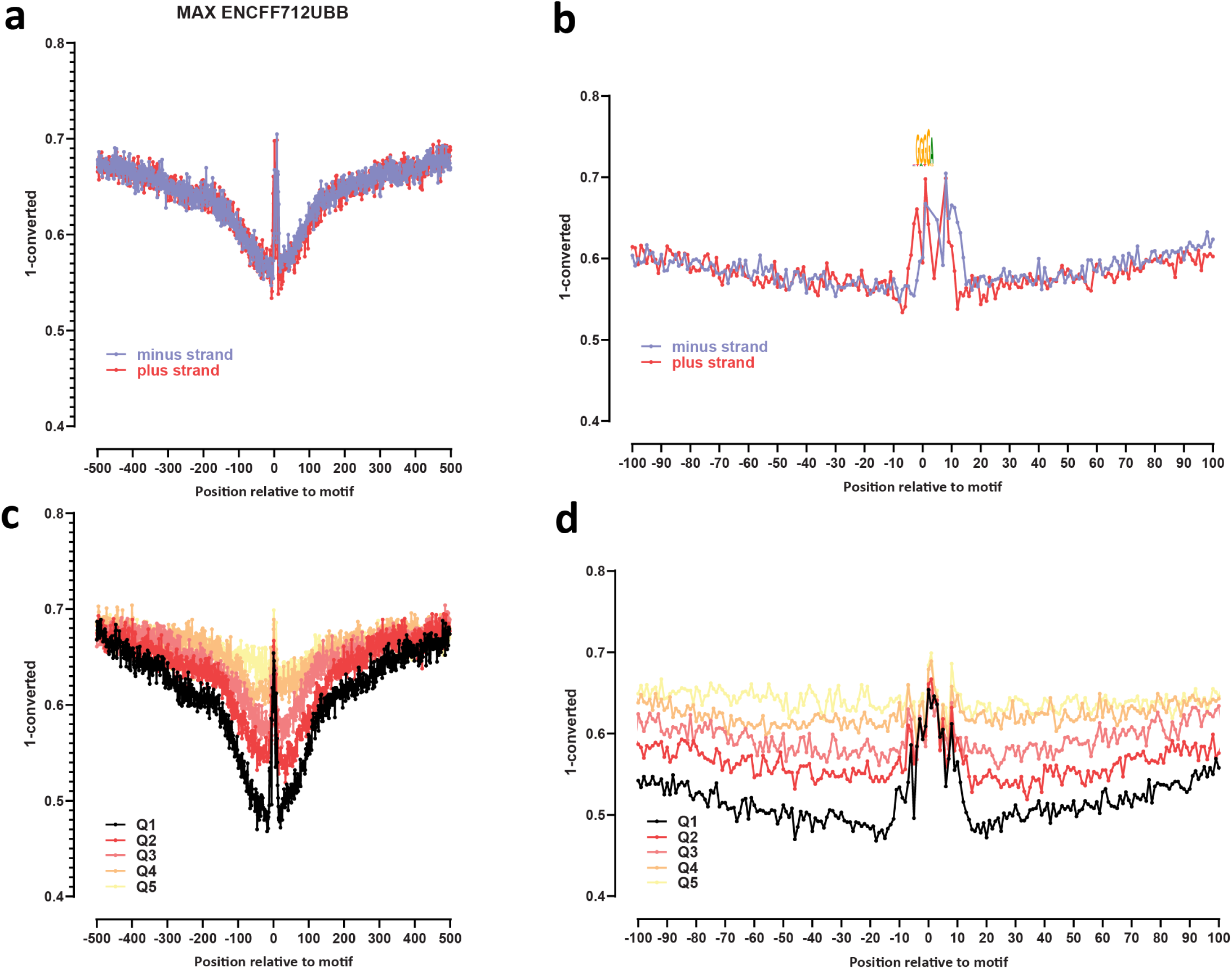
Genome-wide footprint profiles for the MAX transcription factor. GM12878 ssDNA-CseDa01-XL-SMF datasets are shown. (a) Genome-wide footprint profiles over the closest motif to ENCODE ChIP-seq peaks (ENCODE dataset ID indicated on top). Profiles are generated in a strand-aware way with respect to the motif orientation in the genome, and are shown separately for forward and reverse-strand reads. (b) Same as in (a) but zoomed in. The transcription factor sequence recognition motif is shown on top. (c) Genome-wide footprint profiles over the closest motif to ENCODE ChIP-seq peaks for the TF divided into quintiles based on ChIP-seq signal strength. (d) Same as in (c) but zoomed in.

**Supplementary Figure 36:**
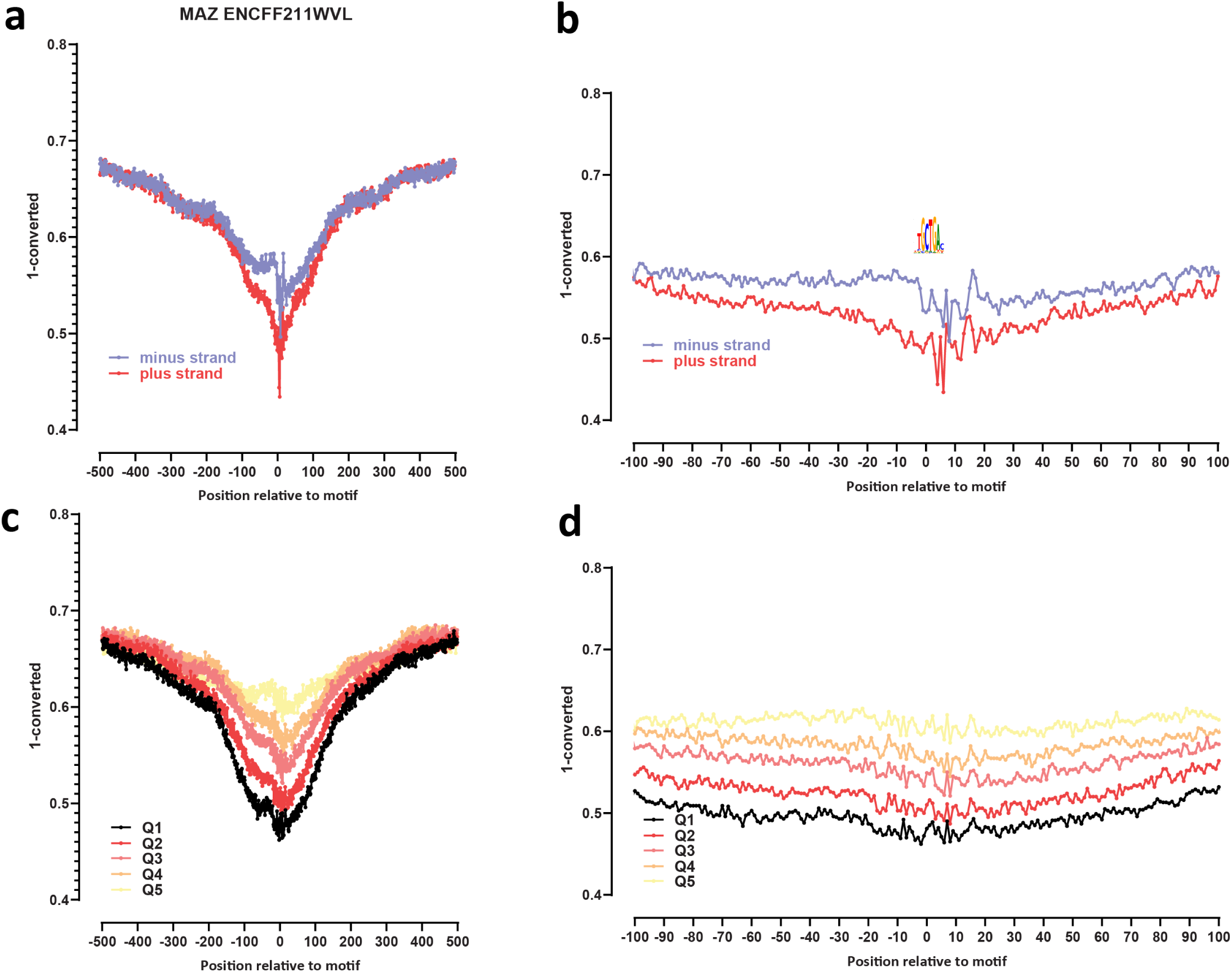
Genome-wide footprint profiles for the MAZ transcription factor. GM12878 ssDNA-CseDa01-XL-SMF datasets are shown. (a) Genome-wide footprint profiles over the closest motif to ENCODE ChIP-seq peaks (ENCODE dataset ID indicated on top). Profiles are generated in a strand-aware way with respect to the motif orientation in the genome, and are shown separately for forward and reverse-strand reads. (b) Same as in (a) but zoomed in. The transcription factor sequence recognition motif is shown on top. (c) Genome-wide footprint profiles over the closest motif to ENCODE ChIP-seq peaks for the TF divided into quintiles based on ChIP-seq signal strength. (d) Same as in (c) but zoomed in.

**Supplementary Figure 37:**
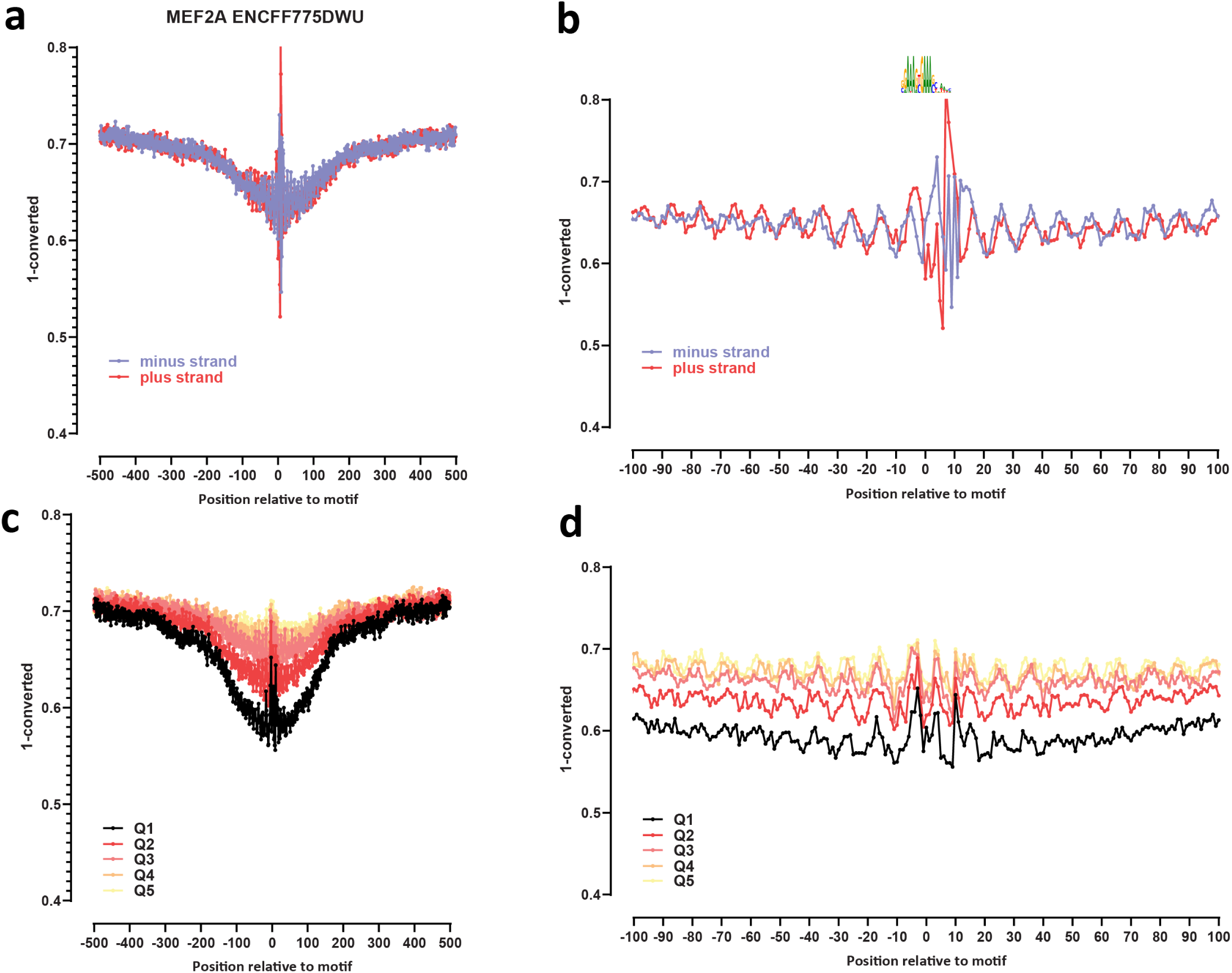
Genome-wide footprint profiles for the MEF2A transcription factor. GM12878 ssDNA-CseDa01-XL-SMF datasets are shown. (a) Genome-wide footprint profiles over the closest motif to ENCODE ChIP-seq peaks (ENCODE dataset ID indicated on top). Profiles are generated in a strand-aware way with respect to the motif orientation in the genome, and are shown separately for forward and reverse-strand reads. (b) Same as in (a) but zoomed in. The transcription factor sequence recognition motif is shown on top. (c) Genome-wide footprint profiles over the closest motif to ENCODE ChIP-seq peaks for the TF divided into quintiles based on ChIP-seq signal strength. (d) Same as in (c) but zoomed in.

**Supplementary Figure 38:**
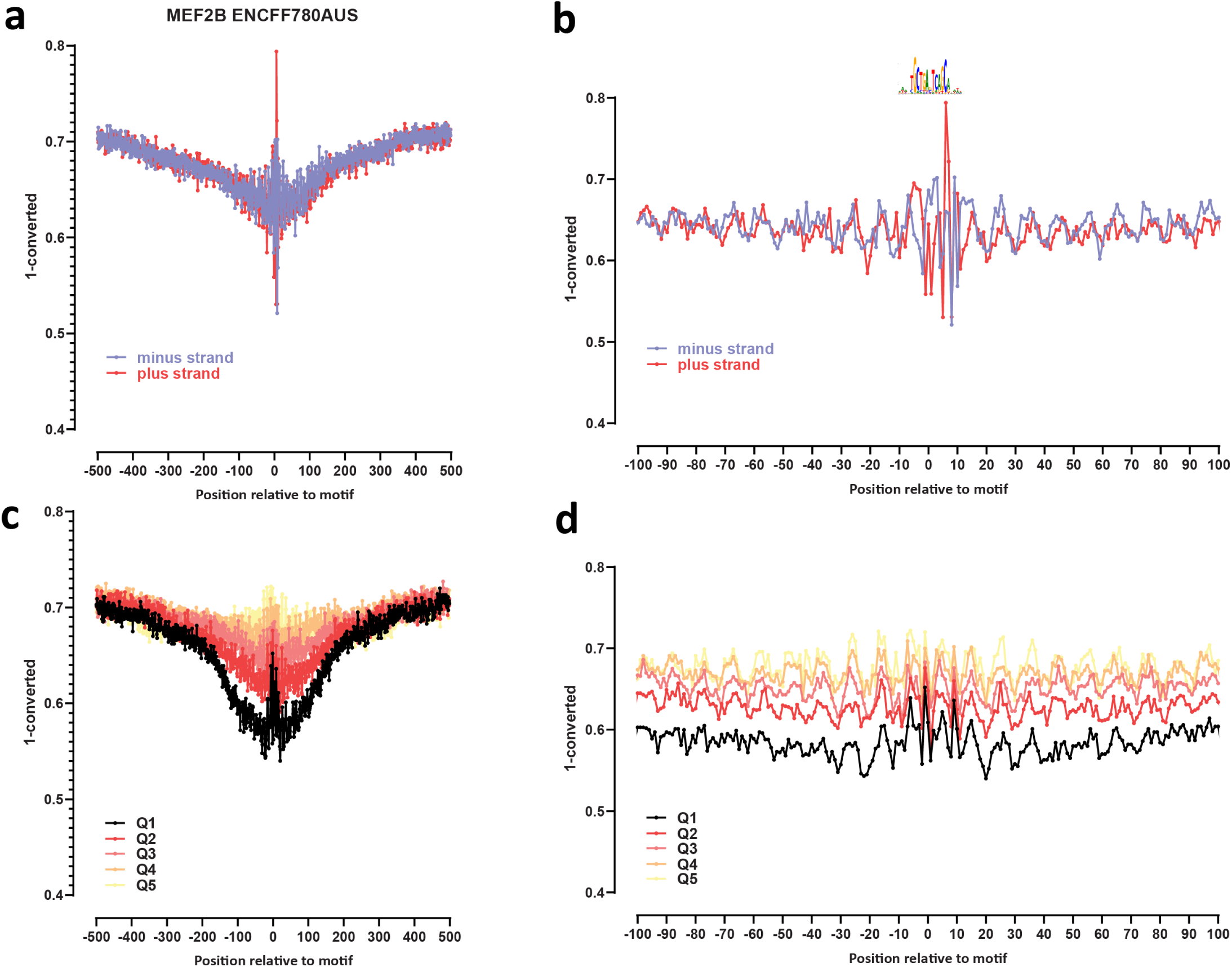
Genome-wide footprint profiles for the MEF2B transcription factor. GM12878 ssDNA-CseDa01-XL-SMF datasets are shown. (a) Genome-wide footprint profiles over the closest motif to ENCODE ChIP-seq peaks (ENCODE dataset ID indicated on top). Profiles are generated in a strand-aware way with respect to the motif orientation in the genome, and are shown separately for forward and reverse-strand reads. (b) Same as in (a) but zoomed in. The transcription factor sequence recognition motif is shown on top. (c) Genome-wide footprint profiles over the closest motif to ENCODE ChIP-seq peaks for the TF divided into quintiles based on ChIP-seq signal strength. (d) Same as in (c) but zoomed in.

**Supplementary Figure 39:**
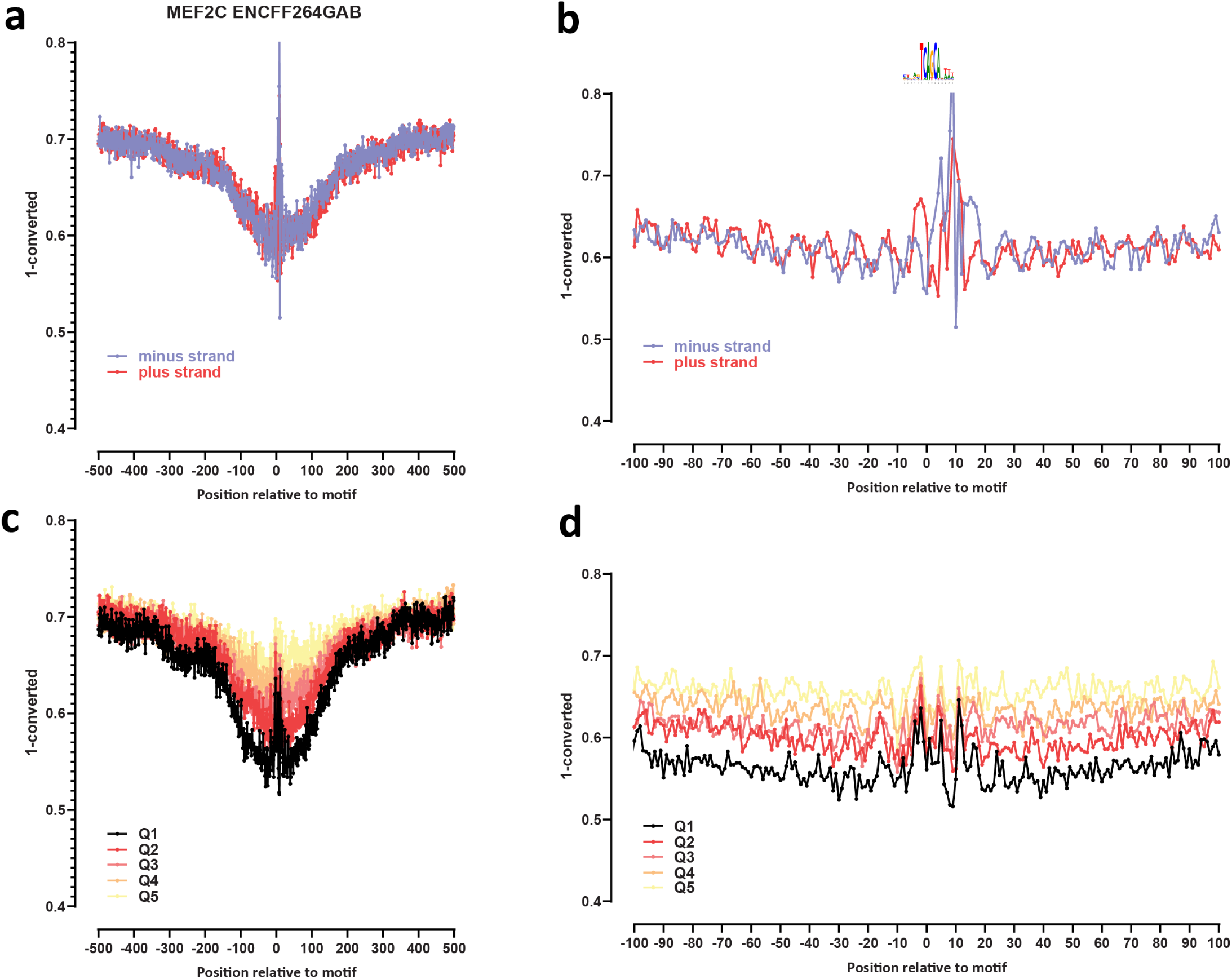
Genome-wide footprint profiles for the MEF2C transcription factor. GM12878 ssDNA-CseDa01-XL-SMF datasets are shown. (a) Genome-wide footprint profiles over the closest motif to ENCODE ChIP-seq peaks (ENCODE dataset ID indicated on top). Profiles are generated in a strand-aware way with respect to the motif orientation in the genome, and are shown separately for forward and reverse-strand reads. (b) Same as in (a) but zoomed in. The transcription factor sequence recognition motif is shown on top. (c) Genome-wide footprint profiles over the closest motif to ENCODE ChIP-seq peaks for the TF divided into quintiles based on ChIP-seq signal strength. (d) Same as in (c) but zoomed in.

**Supplementary Figure 40:**
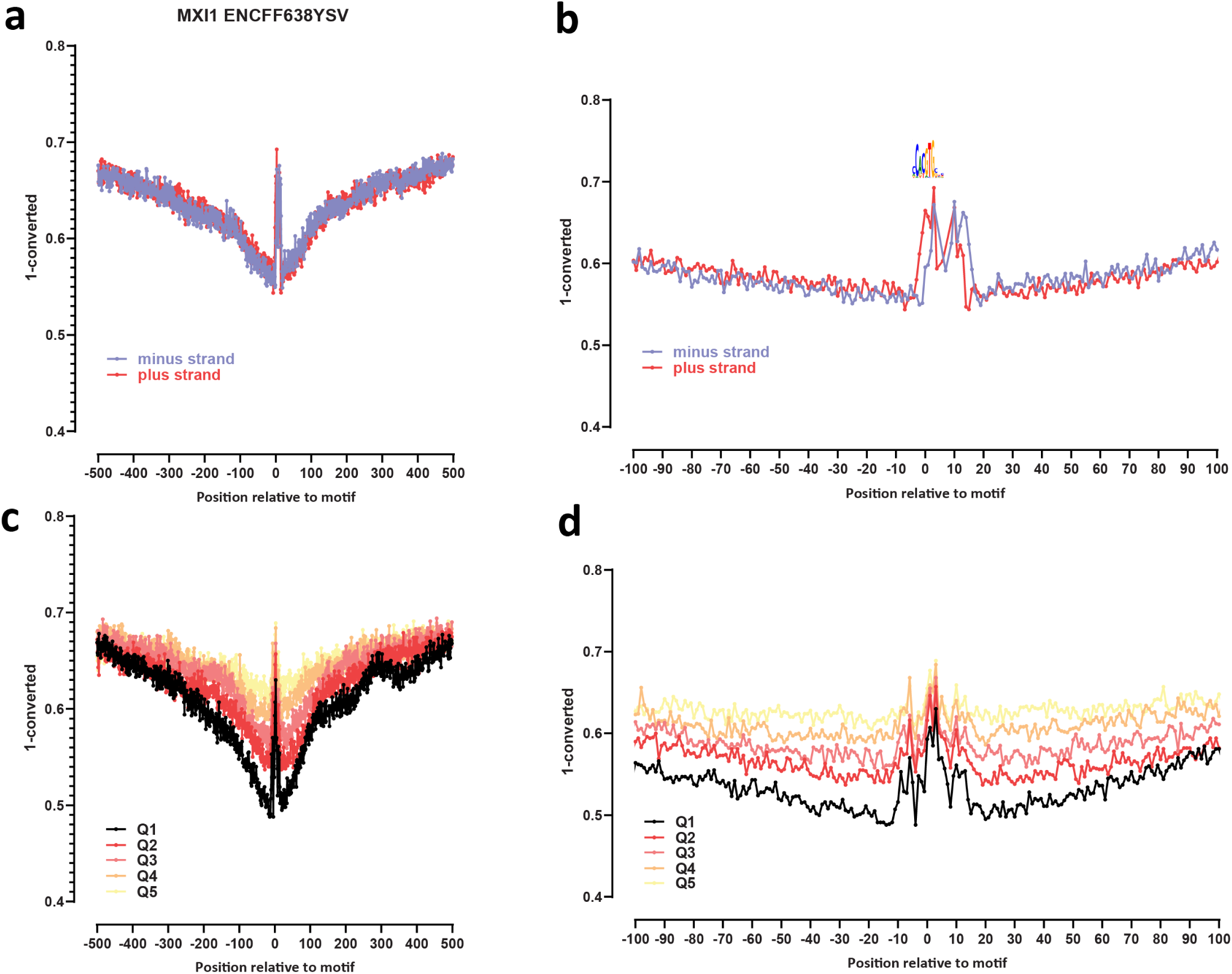
Genome-wide footprint profiles for the MXI1 transcription factor. GM12878 ssDNA-CseDa01-XL-SMF datasets are shown. (a) Genome-wide footprint profiles over the closest motif to ENCODE ChIP-seq peaks (ENCODE dataset ID indicated on top). Profiles are generated in a strand-aware way with respect to the motif orientation in the genome, and are shown separately for forward and reverse-strand reads. (b) Same as in (a) but zoomed in. The transcription factor sequence recognition motif is shown on top. (c) Genome-wide footprint profiles over the closest motif to ENCODE ChIP-seq peaks for the TF divided into quintiles based on ChIP-seq signal strength. (d) Same as in (c) but zoomed in.

**Supplementary Figure 41:**
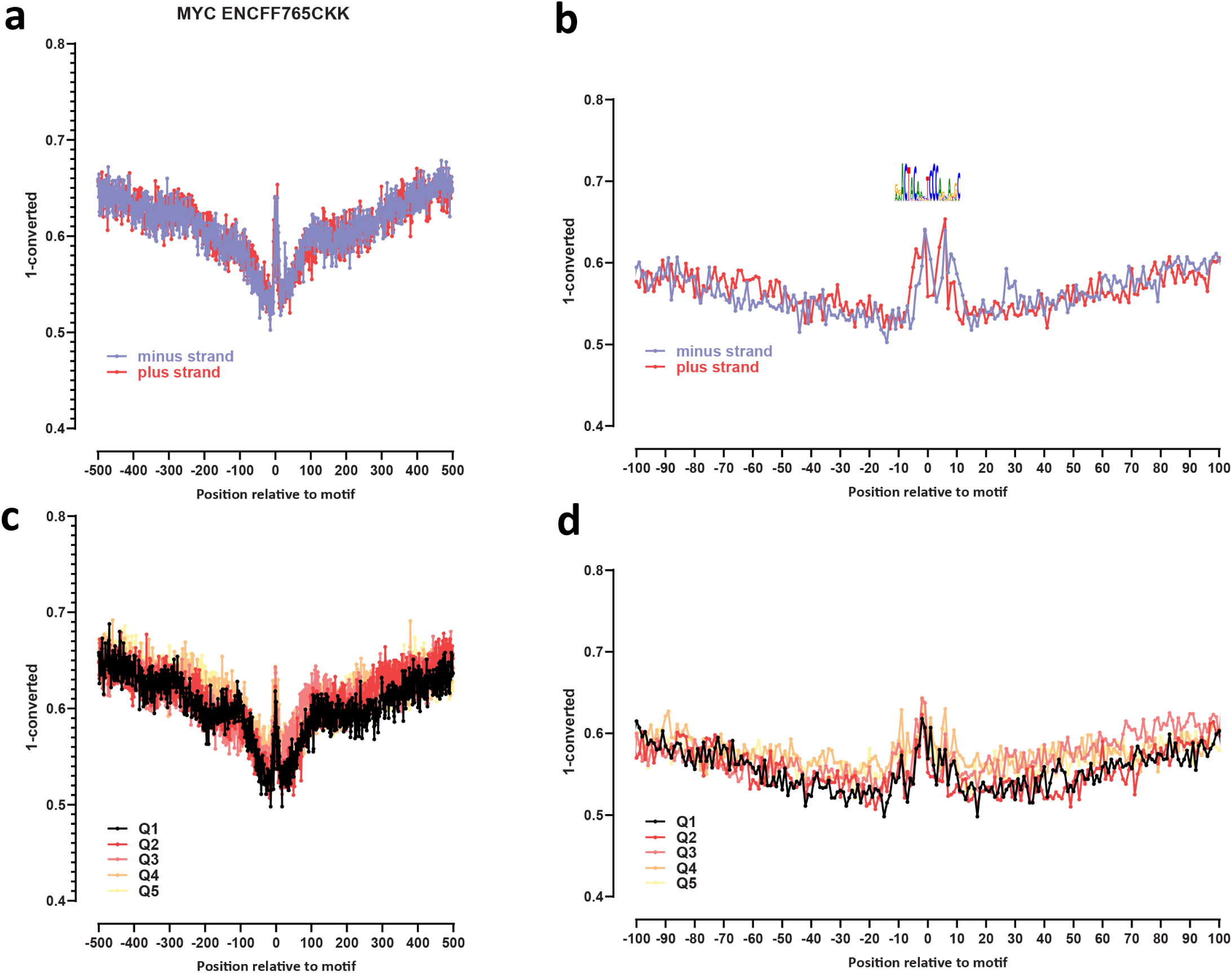
Genome-wide footprint profiles for the MYC transcription factor. GM12878 ssDNA-CseDa01-XL-SMF datasets are shown. (a) Genome-wide footprint profiles over the closest motif to ENCODE ChIP-seq peaks (ENCODE dataset ID indicated on top). Profiles are generated in a strand-aware way with respect to the motif orientation in the genome, and are shown separately for forward and reverse-strand reads. (b) Same as in (a) but zoomed in. The transcription factor sequence recognition motif is shown on top. (c) Genome-wide footprint profiles over the closest motif to ENCODE ChIP-seq peaks for the TF divided into quintiles based on ChIP-seq signal strength. (d) Same as in (c) but zoomed in.

**Supplementary Figure 42:**
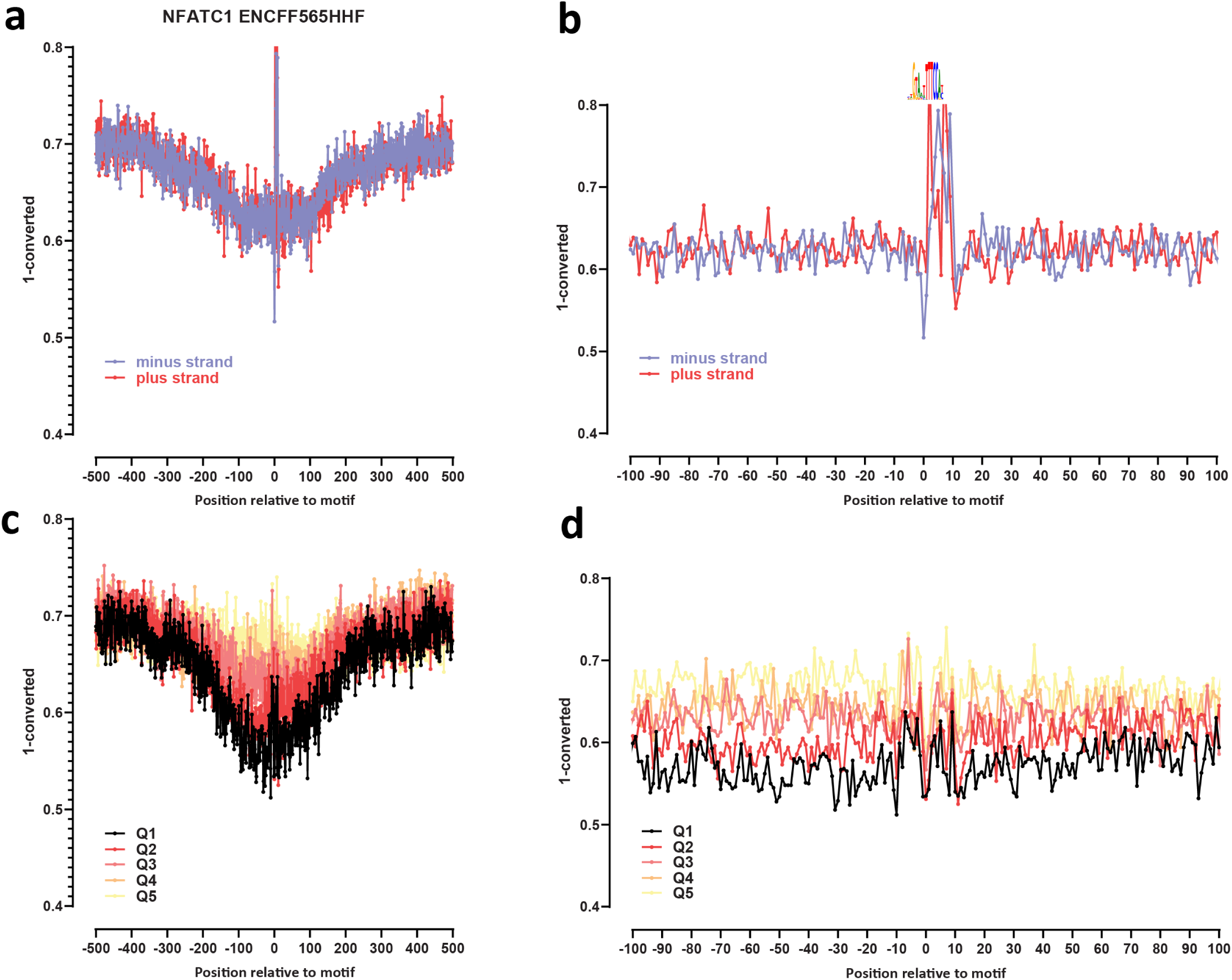
Genome-wide footprint profiles for the NFATC1 transcription factor. GM12878 ssDNA-CseDa01-XL-SMF datasets are shown. (a) Genome-wide footprint profiles over the closest motif to ENCODE ChIP-seq peaks (ENCODE dataset ID indicated on top). Profiles are generated in a strand-aware way with respect to the motif orientation in the genome, and are shown separately for forward and reverse-strand reads. (b) Same as in (a) but zoomed in. The transcription factor sequence recognition motif is shown on top. (c) Genome-wide footprint profiles over the closest motif to ENCODE ChIP-seq peaks for the TF divided into quintiles based on ChIP-seq signal strength. (d) Same as in (c) but zoomed in.

**Supplementary Figure 43:**
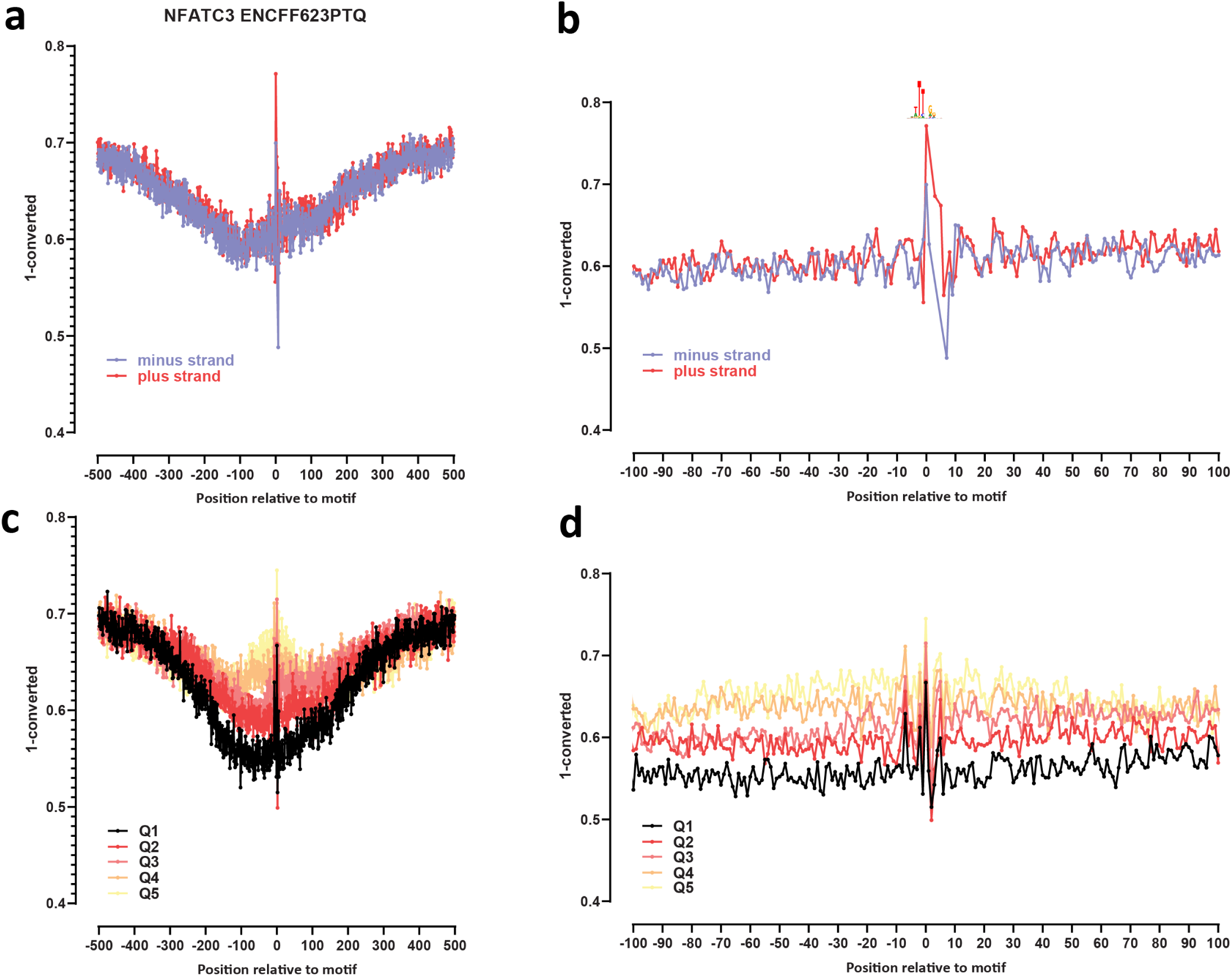
Genome-wide footprint profiles for the NFATC3 transcription factor. GM12878 ssDNA-CseDa01-XL-SMF datasets are shown. (a) Genome-wide footprint profiles over the closest motif to ENCODE ChIP-seq peaks (ENCODE dataset ID indicated on top). Profiles are generated in a strand-aware way with respect to the motif orientation in the genome, and are shown separately for forward and reverse-strand reads. (b) Same as in (a) but zoomed in. The transcription factor sequence recognition motif is shown on top. (c) Genome-wide footprint profiles over the closest motif to ENCODE ChIP-seq peaks for the TF divided into quintiles based on ChIP-seq signal strength. (d) Same as in (c) but zoomed in.

**Supplementary Figure 44:**
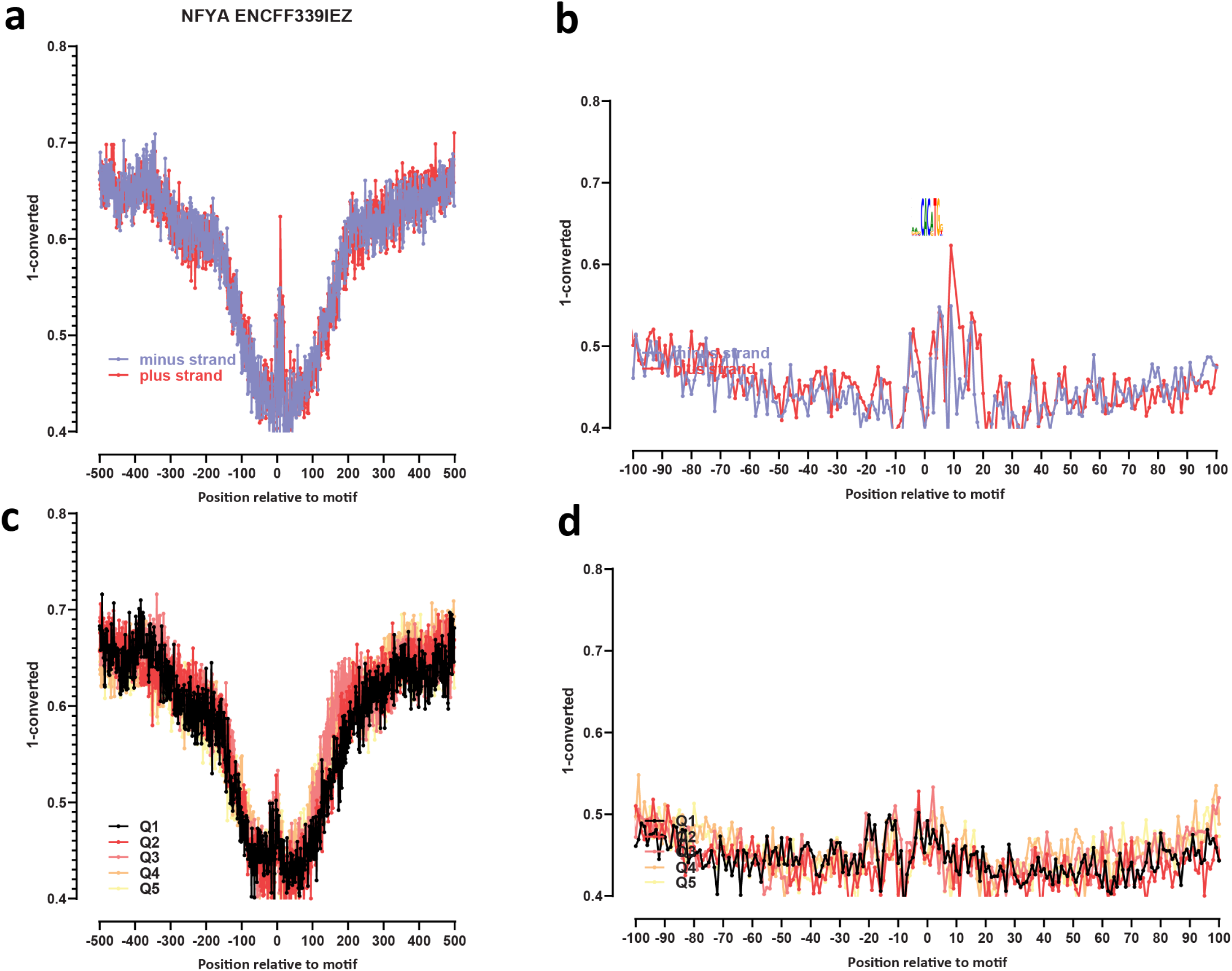
Genome-wide footprint profiles for the NFYA transcription factor. GM12878 ssDNA-CseDa01-XL-SMF datasets are shown. (a) Genome-wide footprint profiles over the closest motif to ENCODE ChIP-seq peaks (ENCODE dataset ID indicated on top). Profiles are generated in a strand-aware way with respect to the motif orientation in the genome, and are shown separately for forward and reverse-strand reads. (b) Same as in (a) but zoomed in. The transcription factor sequence recognition motif is shown on top. (c) Genome-wide footprint profiles over the closest motif to ENCODE ChIP-seq peaks for the TF divided into quintiles based on ChIP-seq signal strength. (d) Same as in (c) but zoomed in.

**Supplementary Figure 45:**
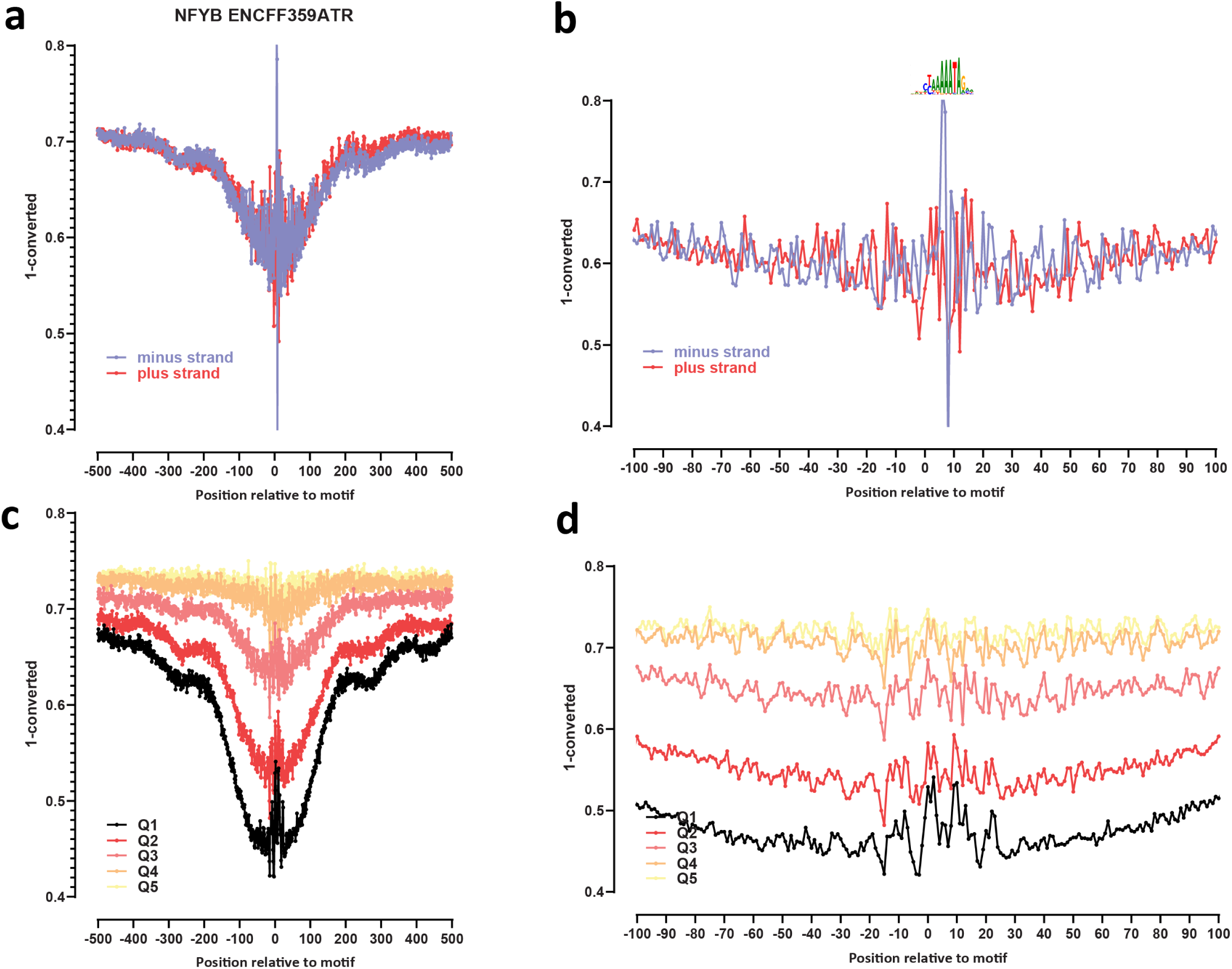
Genome-wide footprint profiles for the NFYB transcription factor. GM12878 ssDNA-CseDa01-XL-SMF datasets are shown. (a) Genome-wide footprint profiles over the closest motif to ENCODE ChIP-seq peaks (ENCODE dataset ID indicated on top). Profiles are generated in a strand-aware way with respect to the motif orientation in the genome, and are shown separately for forward and reverse-strand reads. (b) Same as in (a) but zoomed in. The transcription factor sequence recognition motif is shown on top. (c) Genome-wide footprint profiles over the closest motif to ENCODE ChIP-seq peaks for the TF divided into quintiles based on ChIP-seq signal strength. (d) Same as in (c) but zoomed in.

**Supplementary Figure 46:**
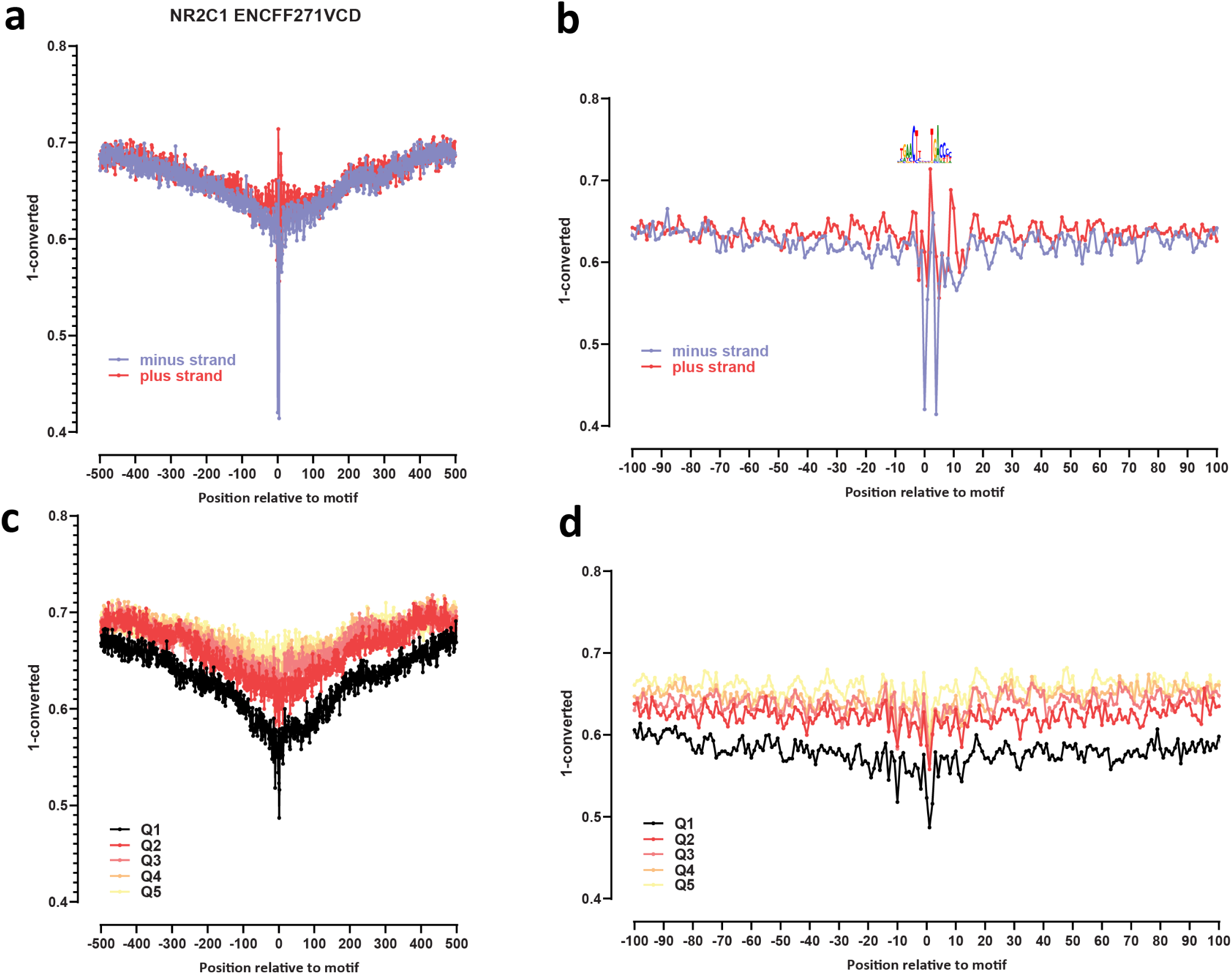
Genome-wide footprint profiles for the NR2C1 transcription factor. GM12878 ssDNA-CseDa01-XL-SMF datasets are shown. (a) Genome-wide footprint profiles over the closest motif to ENCODE ChIP-seq peaks (ENCODE dataset ID indicated on top). Profiles are generated in a strand-aware way with respect to the motif orientation in the genome, and are shown separately for forward and reverse-strand reads. (b) Same as in (a) but zoomed in. The transcription factor sequence recognition motif is shown on top. (c) Genome-wide footprint profiles over the closest motif to ENCODE ChIP-seq peaks for the TF divided into quintiles based on ChIP-seq signal strength. (d) Same as in (c) but zoomed in.

**Supplementary Figure 47:**
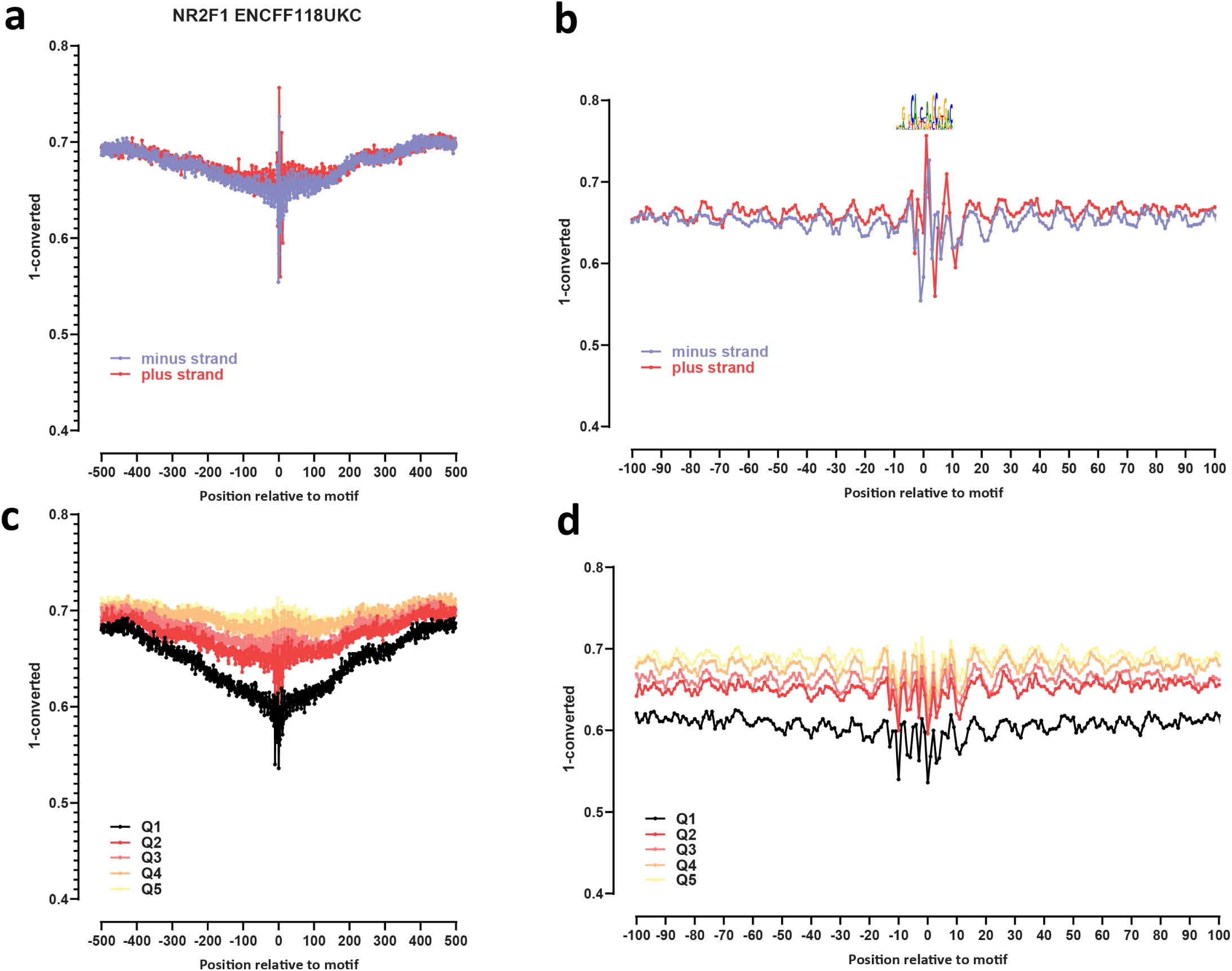
Genome-wide footprint profiles for the NR2F1 transcription factor. GM12878 ssDNA-CseDa01-XL-SMF datasets are shown. (a) Genome-wide footprint profiles over the closest motif to ENCODE ChIP-seq peaks (ENCODE dataset ID indicated on top). Profiles are generated in a strand-aware way with respect to the motif orientation in the genome, and are shown separately for forward and reverse-strand reads. (b) Same as in (a) but zoomed in. The transcription factor sequence recognition motif is shown on top. (c) Genome-wide footprint profiles over the closest motif to ENCODE ChIP-seq peaks for the TF divided into quintiles based on ChIP-seq signal strength. (d) Same as in (c) but zoomed in.

**Supplementary Figure 48:**
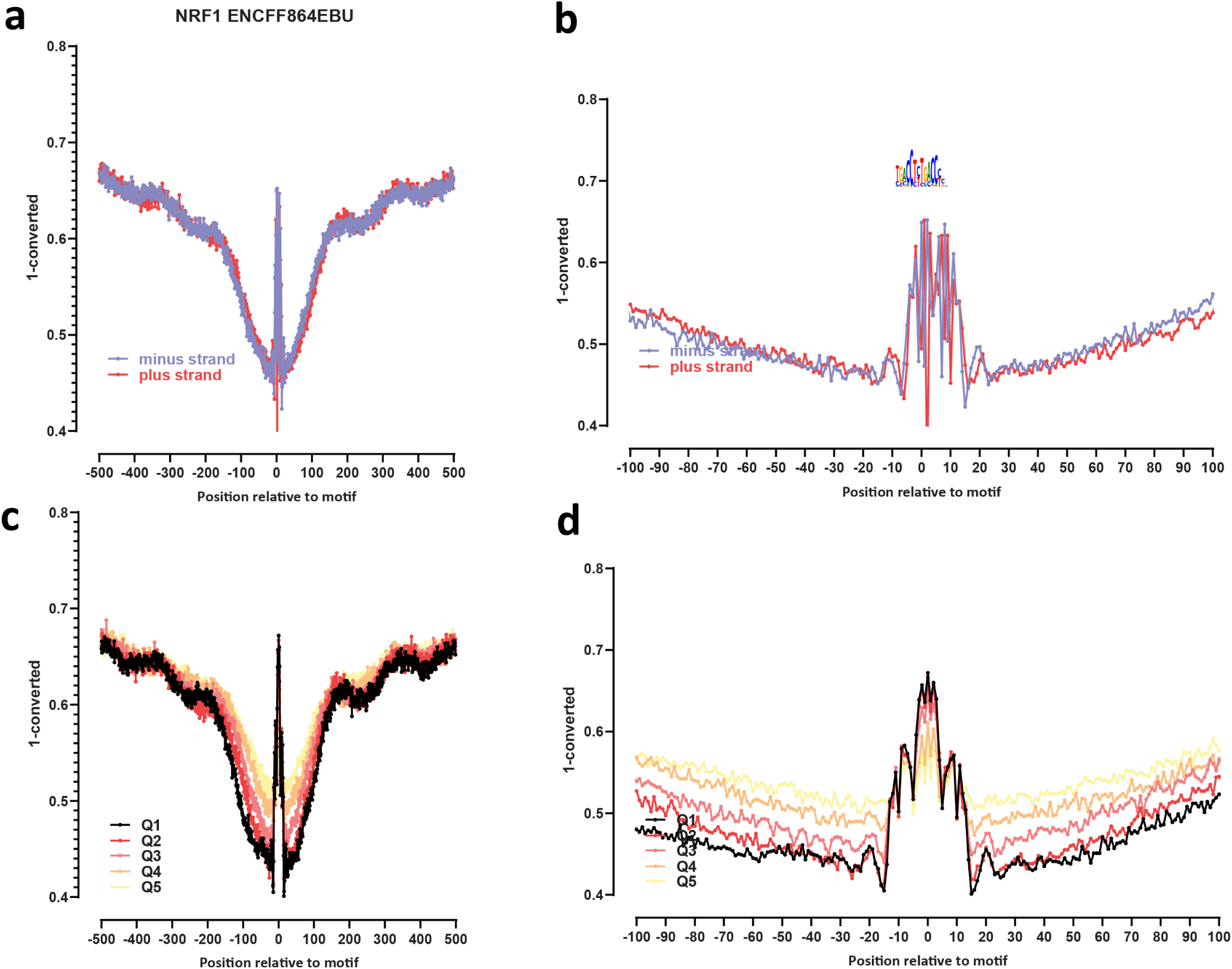
Genome-wide footprint profiles for the NRF1 transcription factor. GM12878 ssDNA-CseDa01-XL-SMF datasets are shown. (a) Genome-wide footprint profiles over the closest motif to ENCODE ChIP-seq peaks (ENCODE dataset ID indicated on top). Profiles are generated in a strand-aware way with respect to the motif orientation in the genome, and are shown separately for forward and reverse-strand reads. (b) Same as in (a) but zoomed in. The transcription factor sequence recognition motif is shown on top. (c) Genome-wide footprint profiles over the closest motif to ENCODE ChIP-seq peaks for the TF divided into quintiles based on ChIP-seq signal strength. (d) Same as in (c) but zoomed in.

**Supplementary Figure 49:**
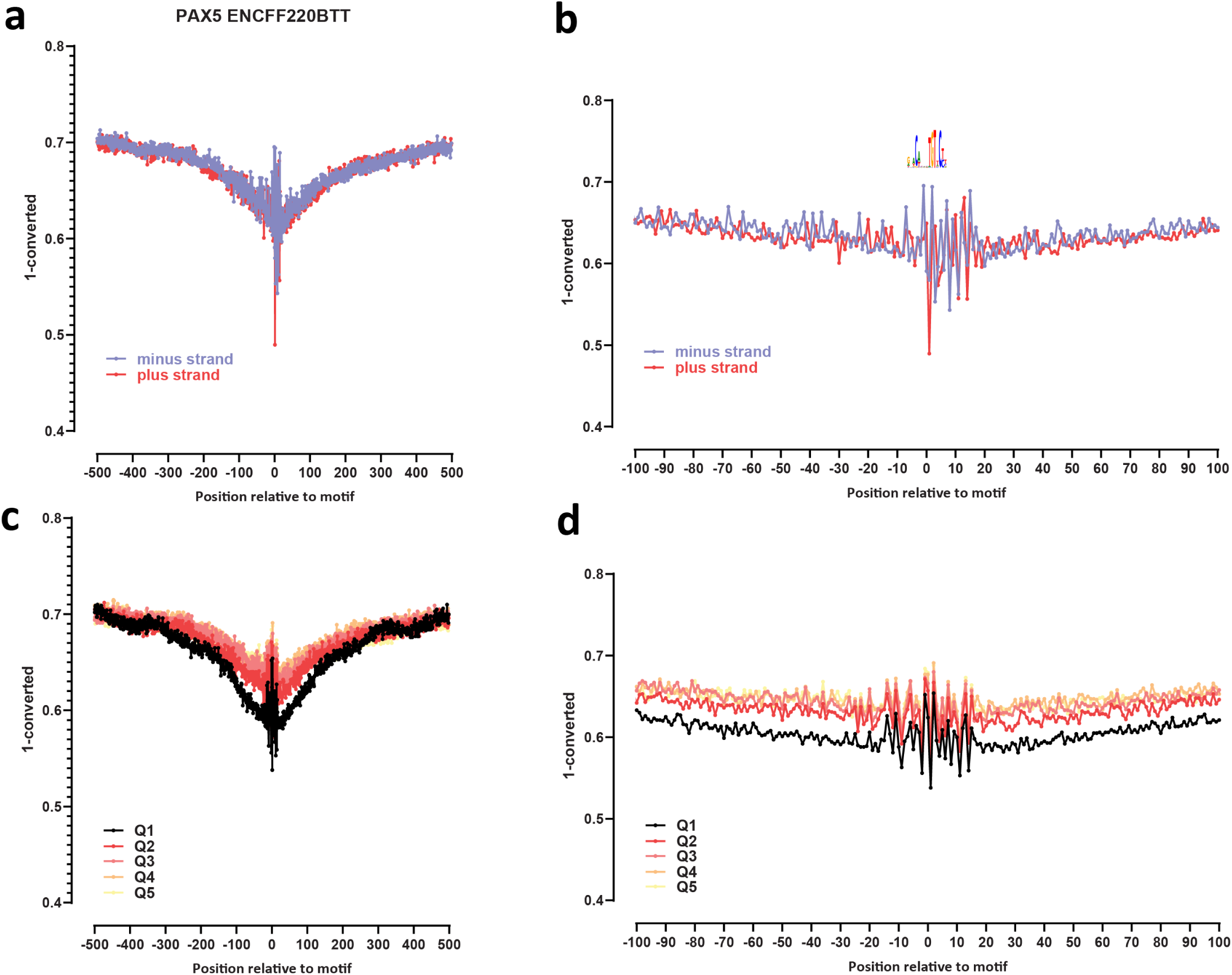
Genome-wide footprint profiles for the PAX5 transcription factor. GM12878 ssDNA-CseDa01-XL-SMF datasets are shown. (a) Genome-wide footprint profiles over the closest motif to ENCODE ChIP-seq peaks (ENCODE dataset ID indicated on top). Profiles are generated in a strand-aware way with respect to the motif orientation in the genome, and are shown separately for forward and reverse-strand reads. (b) Same as in (a) but zoomed in. The transcription factor sequence recognition motif is shown on top. (c) Genome-wide footprint profiles over the closest motif to ENCODE ChIP-seq peaks for the TF divided into quintiles based on ChIP-seq signal strength. (d) Same as in (c) but zoomed in.

**Supplementary Figure 50:**
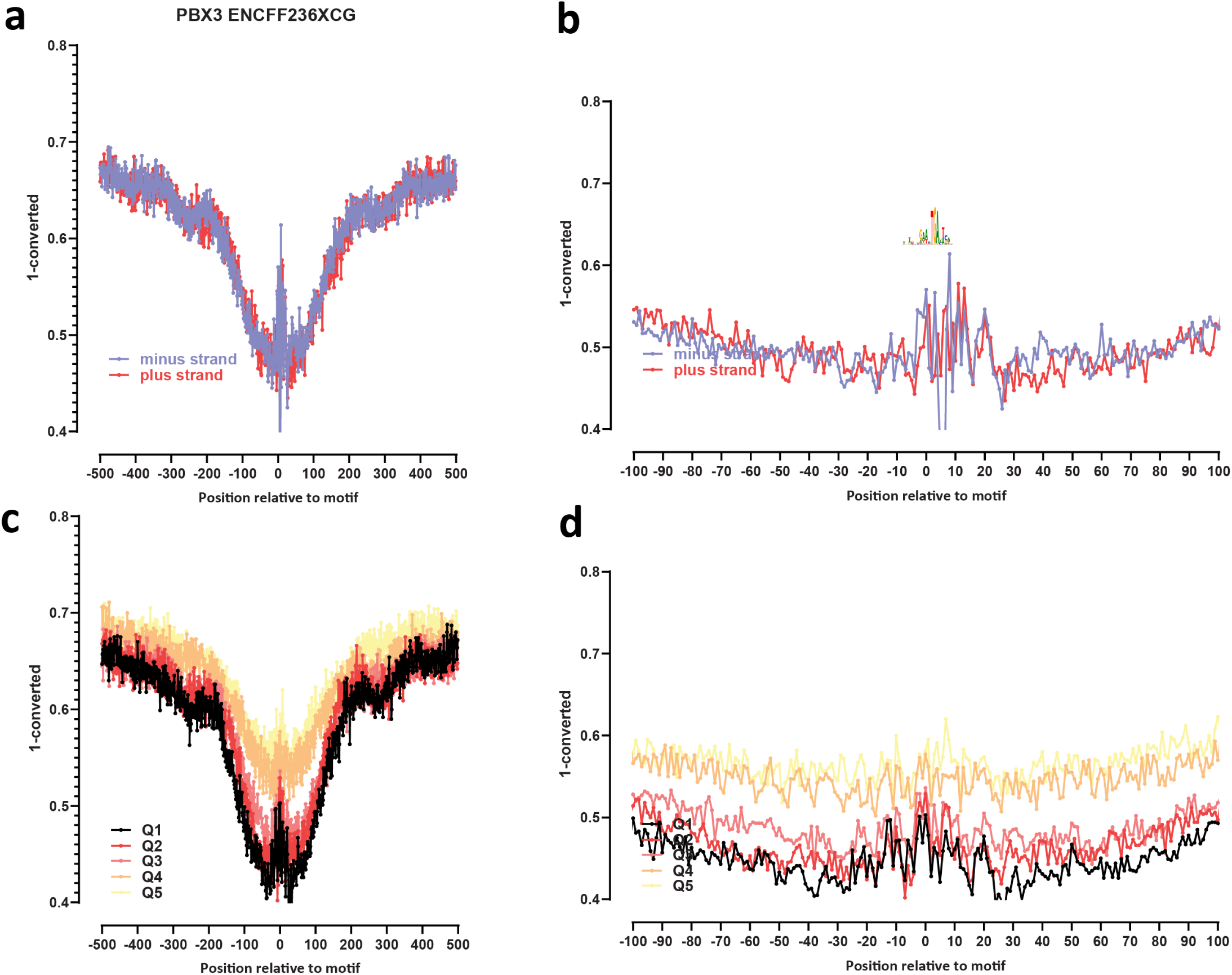
Genome-wide footprint profiles for the PBX3 transcription factor. GM12878 ssDNA-CseDa01-XL-SMF datasets are shown. (a) Genome-wide footprint profiles over the closest motif to ENCODE ChIP-seq peaks (ENCODE dataset ID indicated on top). Profiles are generated in a strand-aware way with respect to the motif orientation in the genome, and are shown separately for forward and reverse-strand reads. (b) Same as in (a) but zoomed in. The transcription factor sequence recognition motif is shown on top. (c) Genome-wide footprint profiles over the closest motif to ENCODE ChIP-seq peaks for the TF divided into quintiles based on ChIP-seq signal strength. (d) Same as in (c) but zoomed in.

**Supplementary Figure 51:**
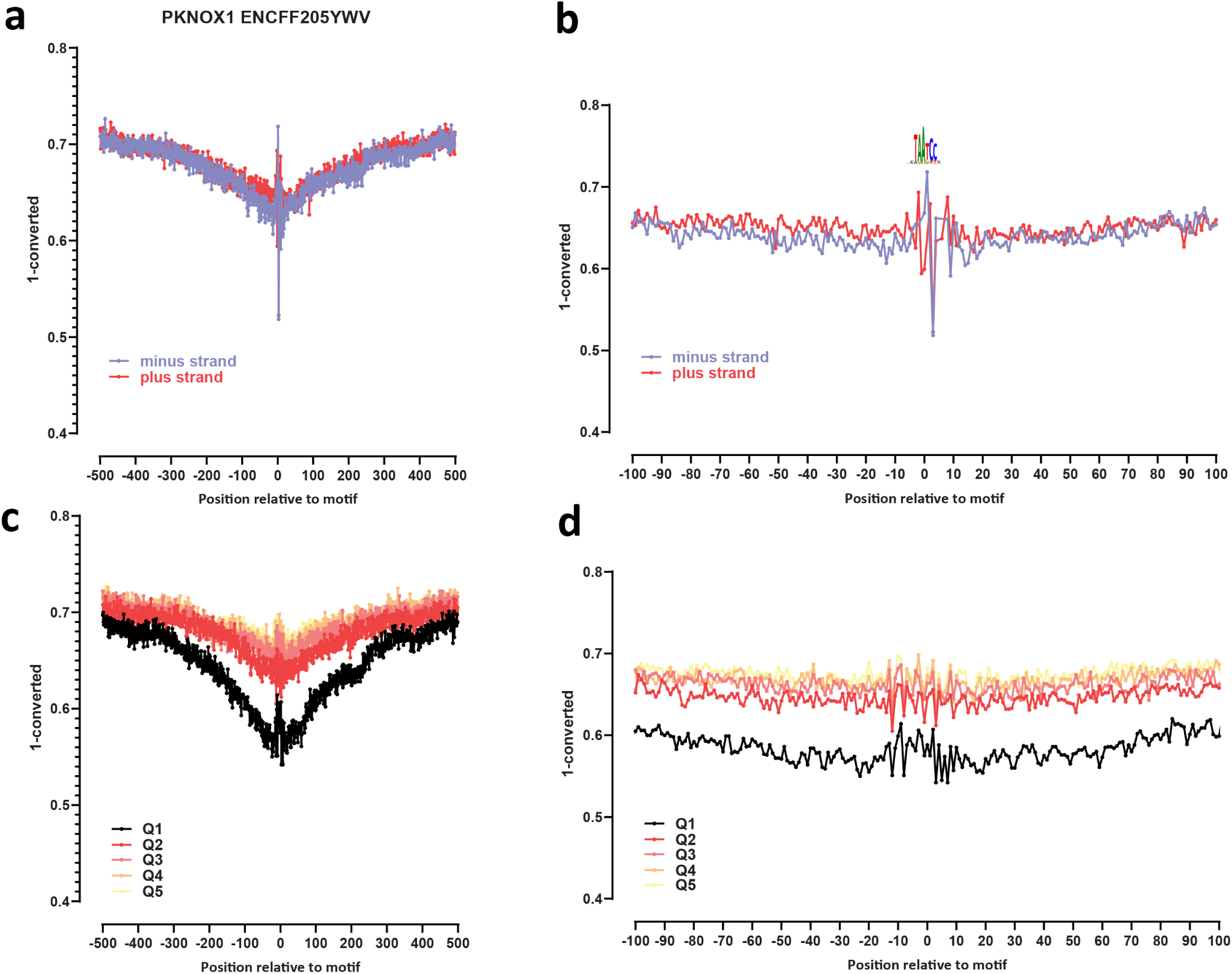
Genome-wide footprint profiles for the PKNOX1 transcription factor. GM12878 ssDNA-CseDa01-XL-SMF datasets are shown. (a) Genome-wide footprint profiles over the closest motif to ENCODE ChIP-seq peaks (ENCODE dataset ID indicated on top). Profiles are generated in a strand-aware way with respect to the motif orientation in the genome, and are shown separately for forward and reverse-strand reads. (b) Same as in (a) but zoomed in. The transcription factor sequence recognition motif is shown on top. (c) Genome-wide footprint profiles over the closest motif to ENCODE ChIP-seq peaks for the TF divided into quintiles based on ChIP-seq signal strength. (d) Same as in (c) but zoomed in.

**Supplementary Figure 52:**
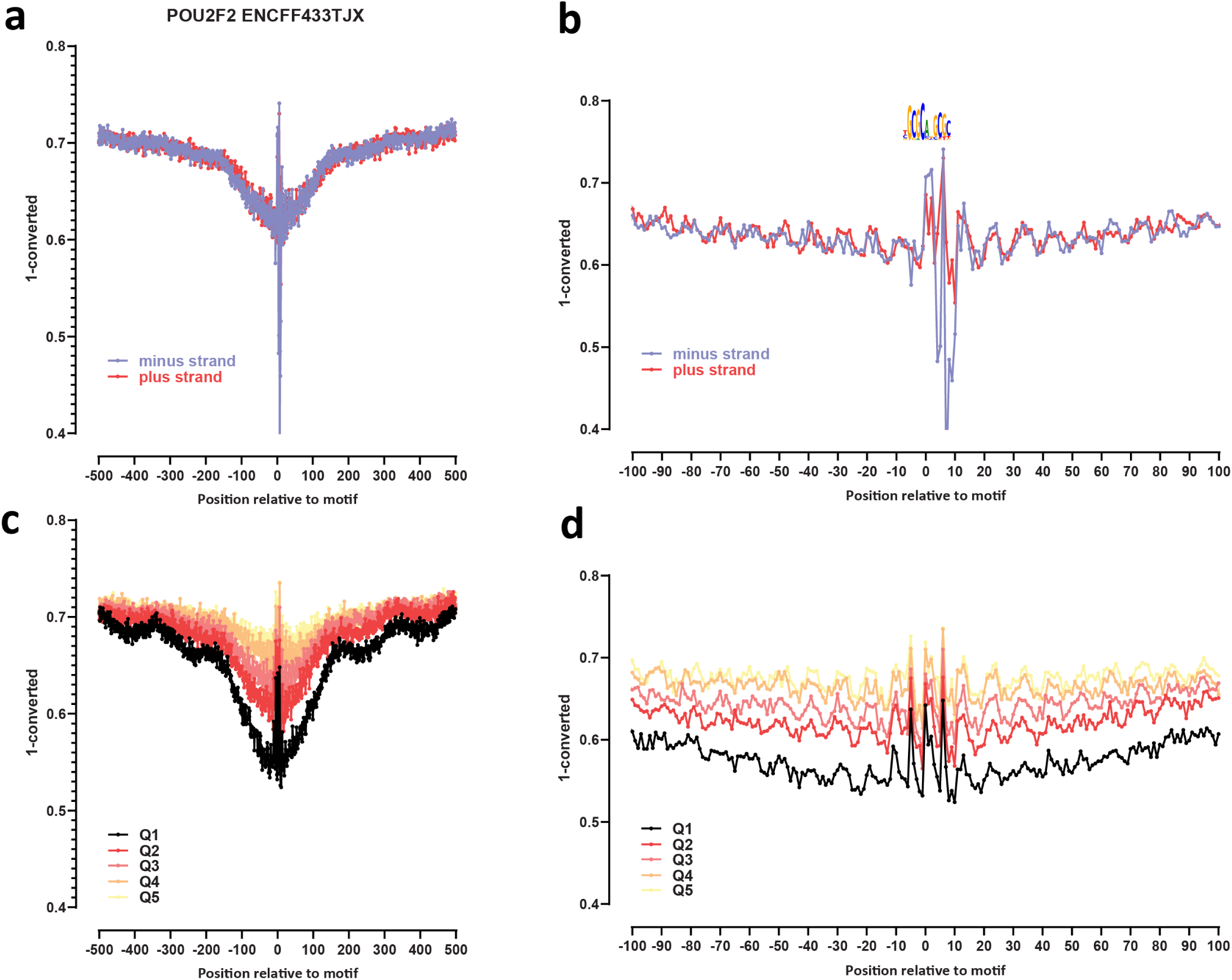
Genome-wide footprint profiles for the POU2F2 transcription factor. GM12878 ssDNA-CseDa01-XL-SMF datasets are shown. (a) Genome-wide footprint profiles over the closest motif to ENCODE ChIP-seq peaks (ENCODE dataset ID indicated on top). Profiles are generated in a strand-aware way with respect to the motif orientation in the genome, and are shown separately for forward and reverse-strand reads. (b) Same as in (a) but zoomed in. The transcription factor sequence recognition motif is shown on top. (c) Genome-wide footprint profiles over the closest motif to ENCODE ChIP-seq peaks for the TF divided into quintiles based on ChIP-seq signal strength. (d) Same as in (c) but zoomed in.

**Supplementary Figure 53:**
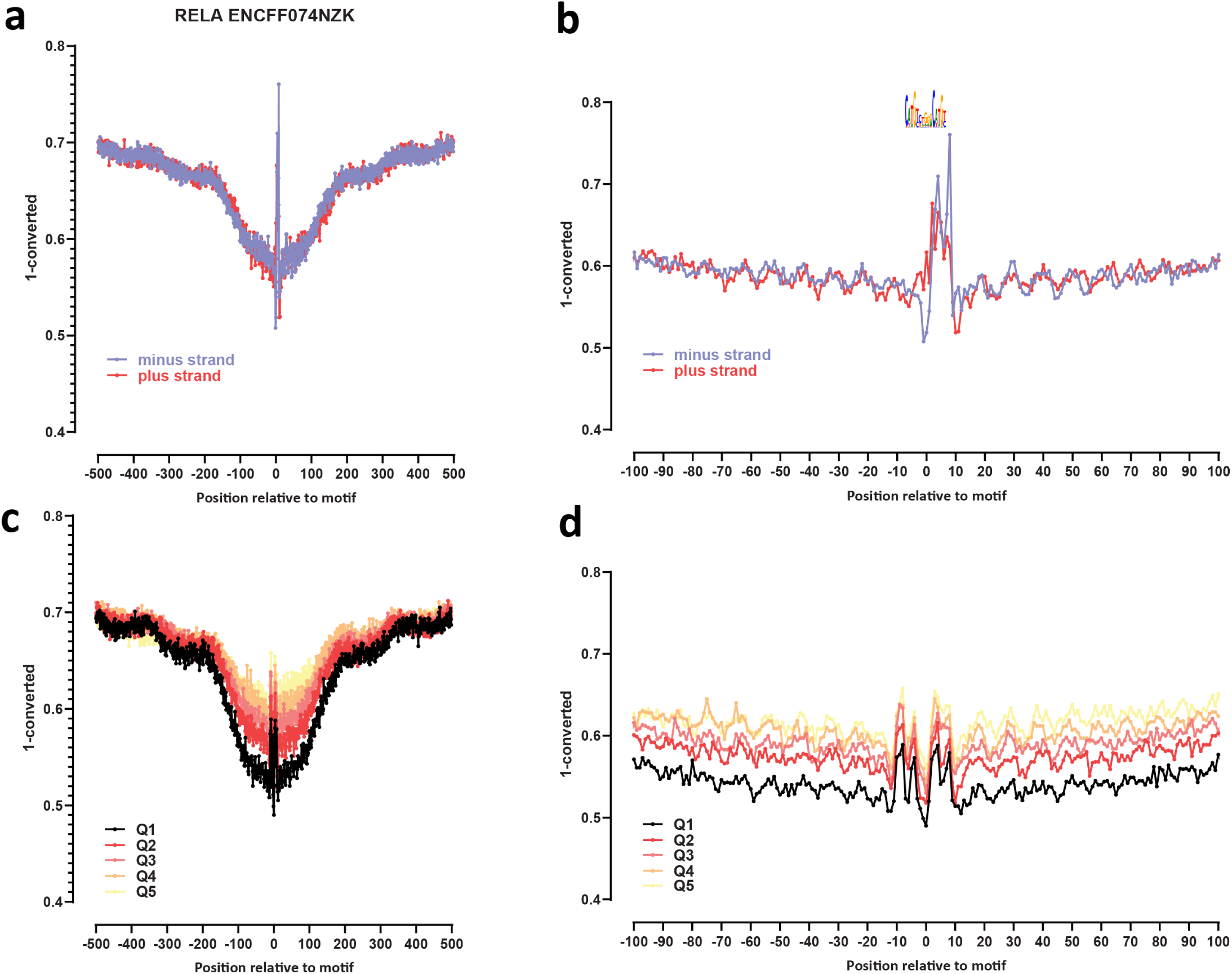
Genome-wide footprint profiles for the RELA transcription factor. GM12878 ssDNA-CseDa01-XL-SMF datasets are shown. (a) Genome-wide footprint profiles over the closest motif to ENCODE ChIP-seq peaks (ENCODE dataset ID indicated on top). Profiles are generated in a strand-aware way with respect to the motif orientation in the genome, and are shown separately for forward and reverse-strand reads. (b) Same as in (a) but zoomed in. The transcription factor sequence recognition motif is shown on top. (c) Genome-wide footprint profiles over the closest motif to ENCODE ChIP-seq peaks for the TF divided into quintiles based on ChIP-seq signal strength. (d) Same as in (c) but zoomed in.

**Supplementary Figure 54:**
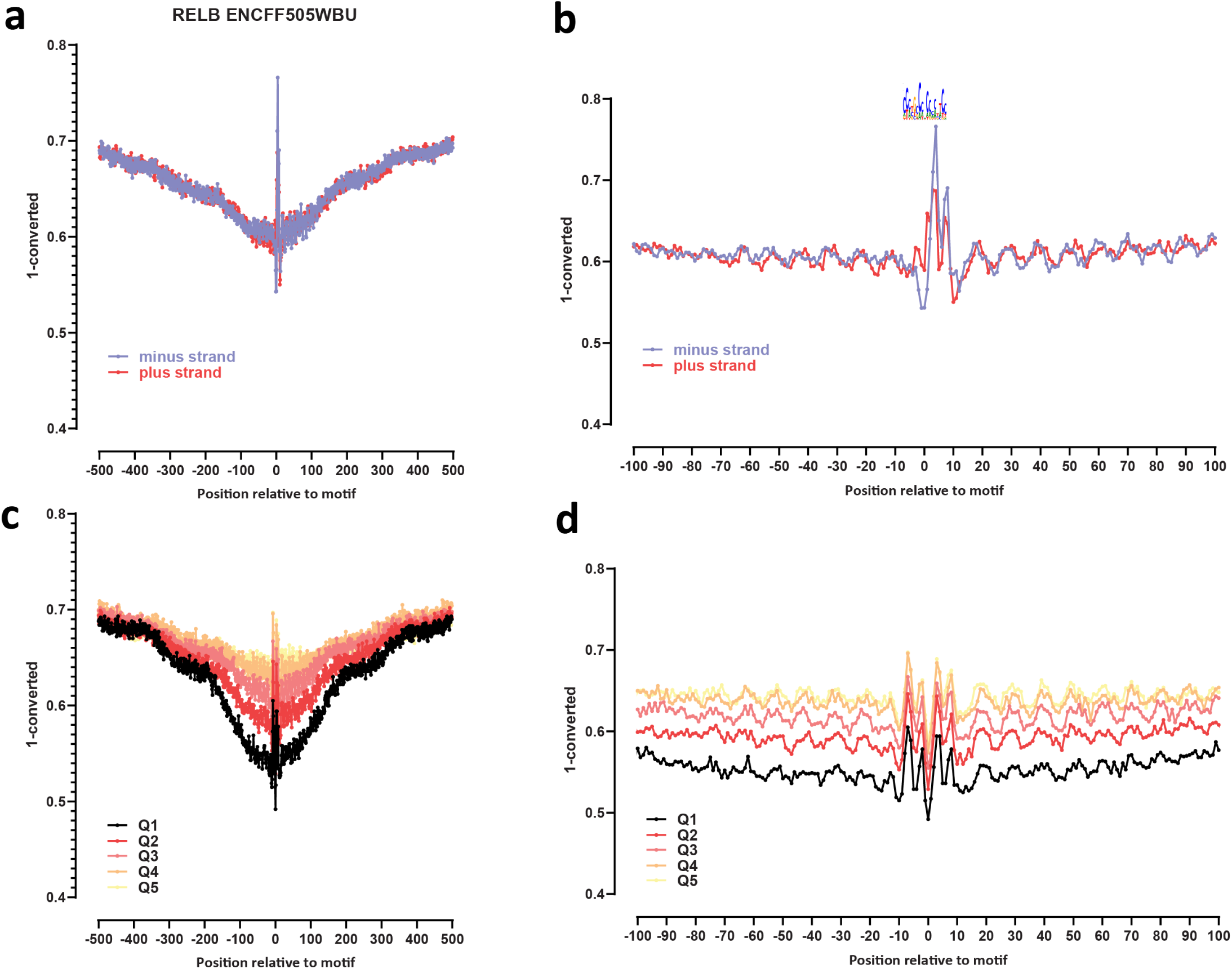
Genome-wide footprint profiles for the RELB transcription factor. GM12878 ssDNA-CseDa01-XL-SMF datasets are shown. (a) Genome-wide footprint profiles over the closest motif to ENCODE ChIP-seq peaks (ENCODE dataset ID indicated on top). Profiles are generated in a strand-aware way with respect to the motif orientation in the genome, and are shown separately for forward and reverse-strand reads. (b) Same as in (a) but zoomed in. The transcription factor sequence recognition motif is shown on top. (c) Genome-wide footprint profiles over the closest motif to ENCODE ChIP-seq peaks for the TF divided into quintiles based on ChIP-seq signal strength. (d) Same as in (c) but zoomed in.

**Supplementary Figure 55:**
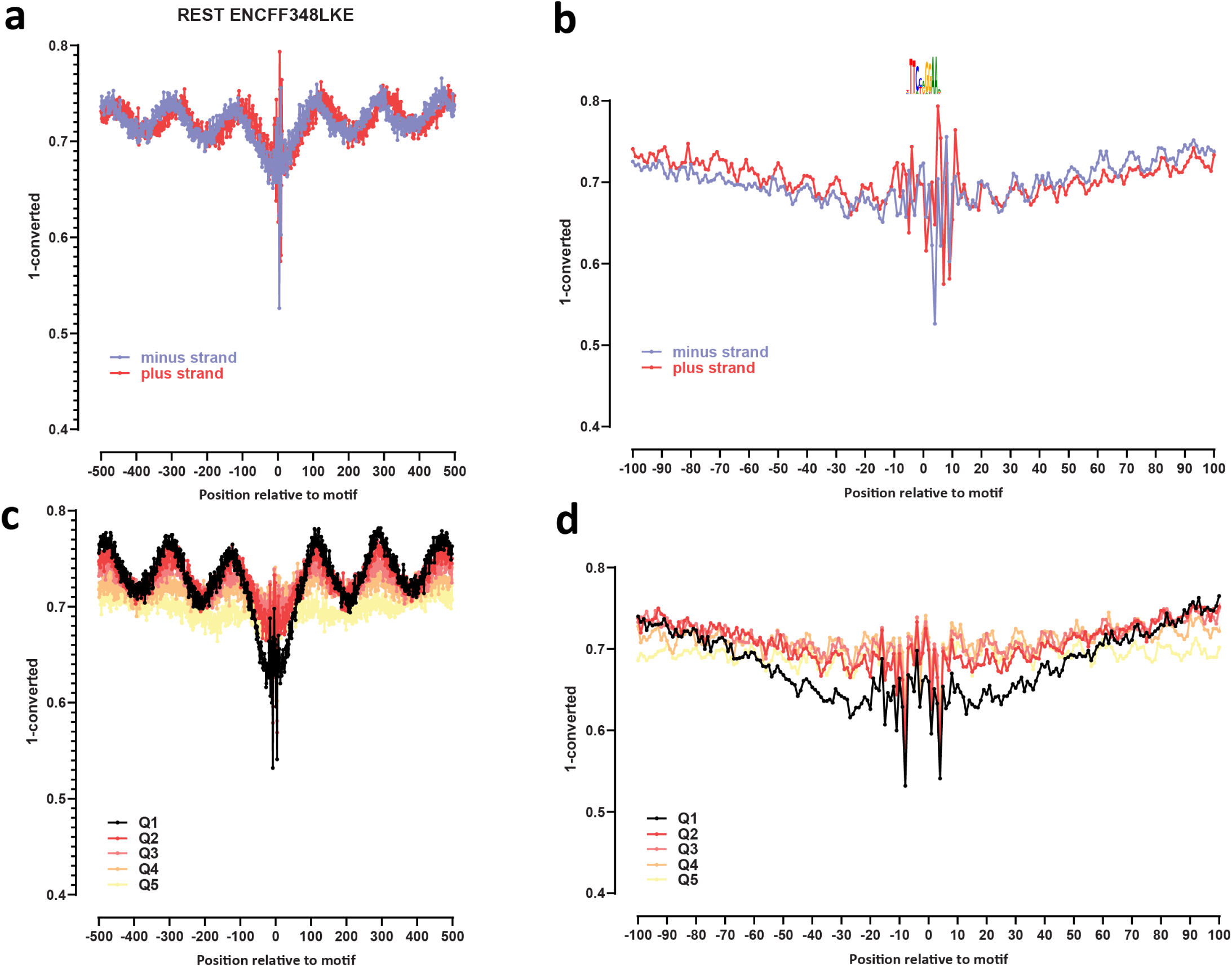
Genome-wide footprint profiles for the REST/NRSF transcription factor. GM12878 ssDNA-CseDa01-XL-SMF datasets are shown. (a) Genome-wide footprint profiles over the closest motif to ENCODE ChIP-seq peaks (ENCODE dataset ID indicated on top). Profiles are generated in a strand-aware way with respect to the motif orientation in the genome, and are shown separately for forward and reverse-strand reads. (b) Same as in (a) but zoomed in. The transcription factor sequence recognition motif is shown on top. (c) Genome-wide footprint profiles over the closest motif to ENCODE ChIP-seq peaks for the TF divided into quintiles based on ChIP-seq signal strength. (d) Same as in (c) but zoomed in.

**Supplementary Figure 56:**
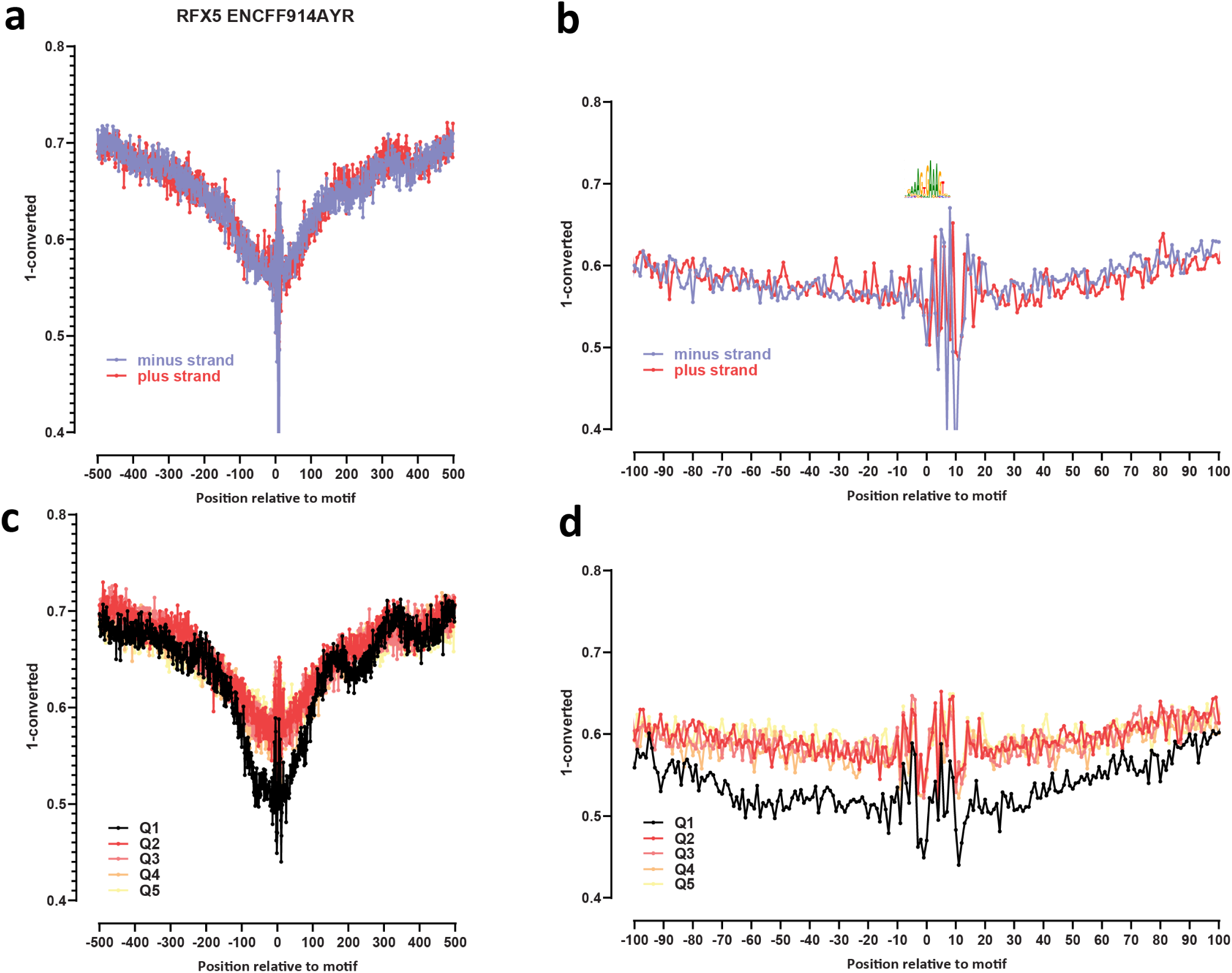
Genome-wide footprint profiles for the RFX5 transcription factor. GM12878 ssDNA-CseDa01-XL-SMF datasets are shown. (a) Genome-wide footprint profiles over the closest motif to ENCODE ChIP-seq peaks (ENCODE dataset ID indicated on top). Profiles are generated in a strand-aware way with respect to the motif orientation in the genome, and are shown separately for forward and reverse-strand reads. (b) Same as in (a) but zoomed in. The transcription factor sequence recognition motif is shown on top. (c) Genome-wide footprint profiles over the closest motif to ENCODE ChIP-seq peaks for the TF divided into quintiles based on ChIP-seq signal strength. (d) Same as in (c) but zoomed in.

**Supplementary Figure 57:**
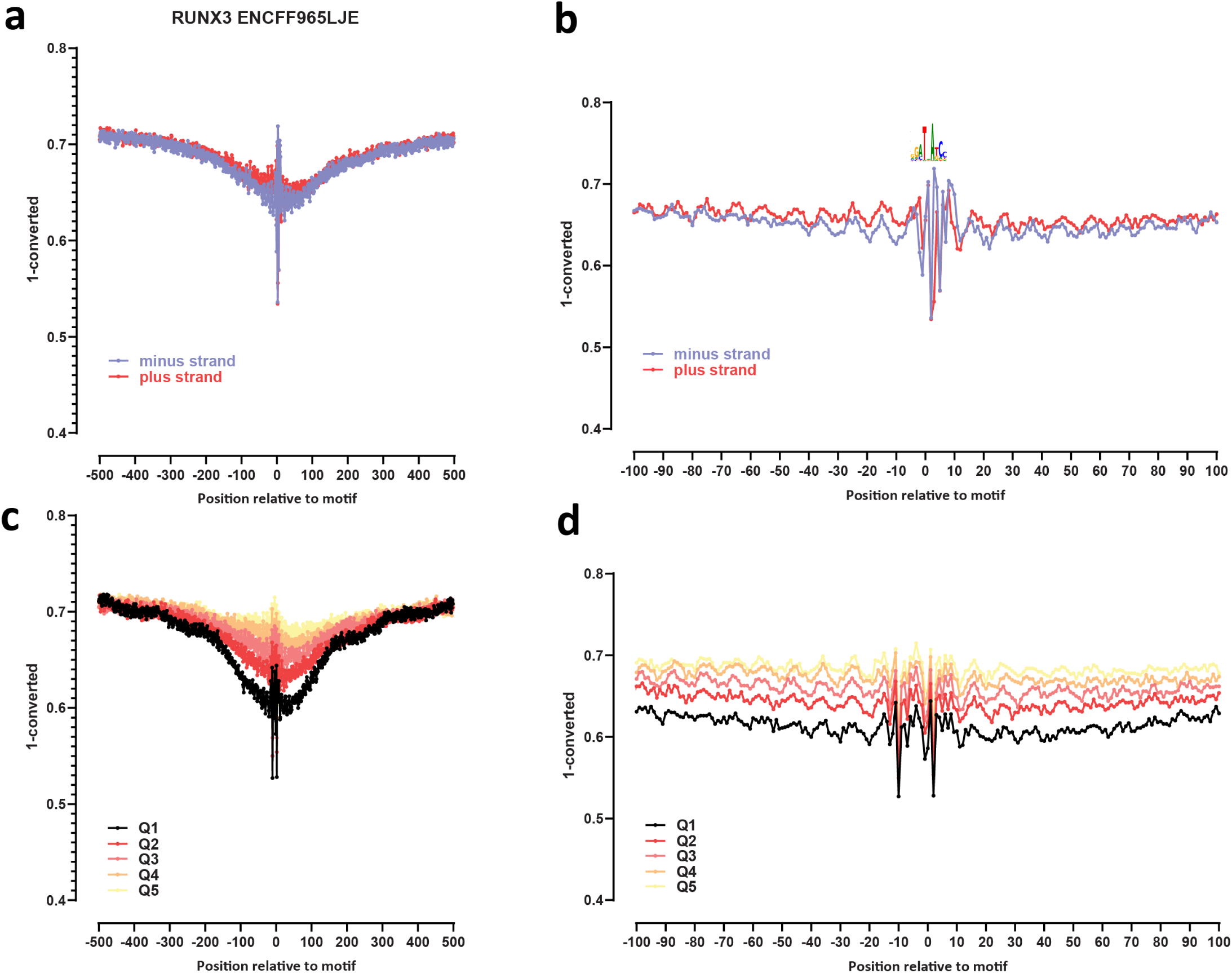
Genome-wide footprint profiles for the RUNX3 transcription factor. GM12878 ssDNA-CseDa01-XL-SMF datasets are shown. (a) Genome-wide footprint profiles over the closest motif to ENCODE ChIP-seq peaks (ENCODE dataset ID indicated on top). Profiles are generated in a strand-aware way with respect to the motif orientation in the genome, and are shown separately for forward and reverse-strand reads. (b) Same as in (a) but zoomed in. The transcription factor sequence recognition motif is shown on top. (c) Genome-wide footprint profiles over the closest motif to ENCODE ChIP-seq peaks for the TF divided into quintiles based on ChIP-seq signal strength. (d) Same as in (c) but zoomed in.

**Supplementary Figure 58:**
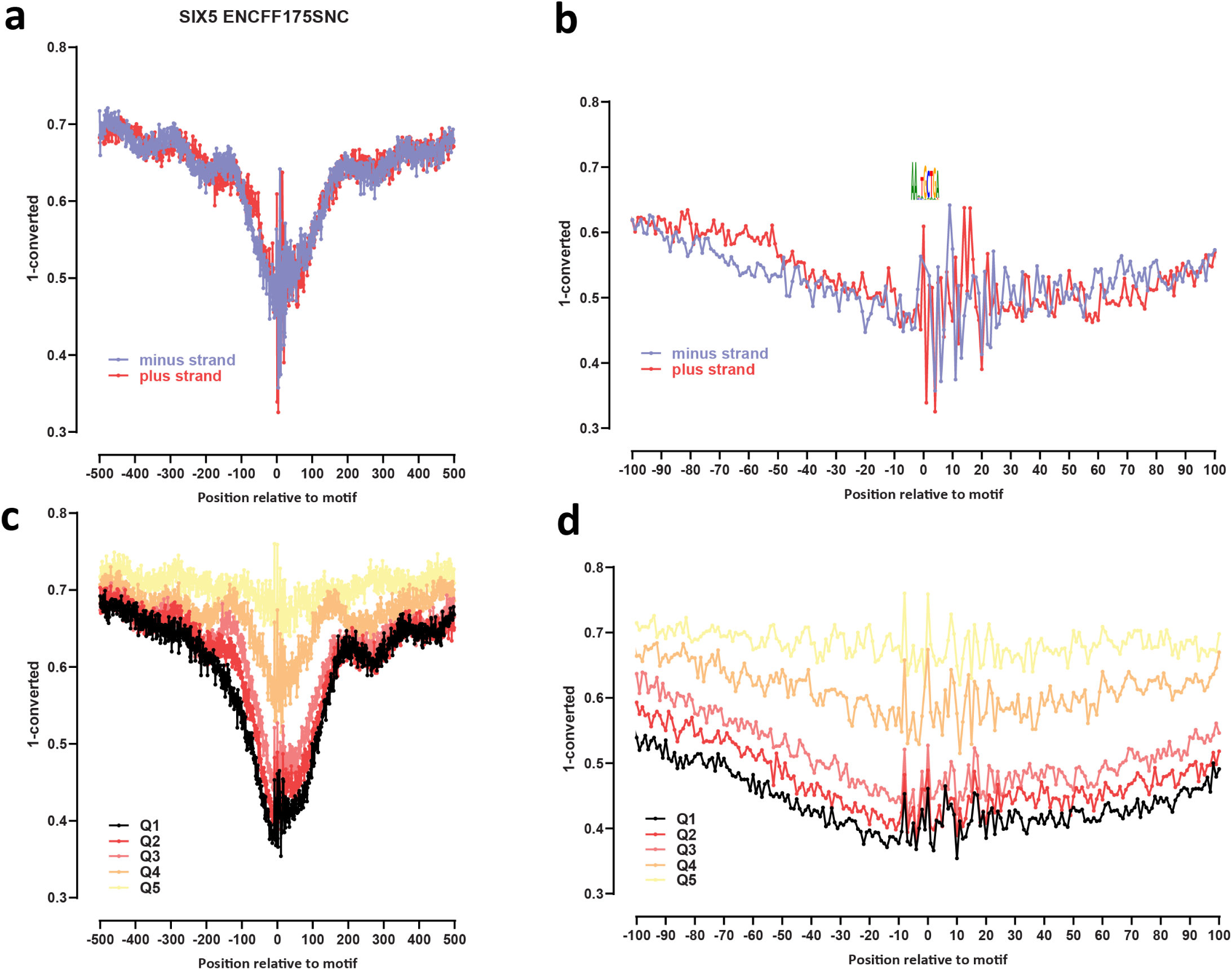
Genome-wide footprint profiles for the SIX5 transcription factor. GM12878 ssDNA-CseDa01-XL-SMF datasets are shown. (a) Genome-wide footprint profiles over the closest motif to ENCODE ChIP-seq peaks (ENCODE dataset ID indicated on top). Profiles are generated in a strand-aware way with respect to the motif orientation in the genome, and are shown separately for forward and reverse-strand reads. (b) Same as in (a) but zoomed in. The transcription factor sequence recognition motif is shown on top. (c) Genome-wide footprint profiles over the closest motif to ENCODE ChIP-seq peaks for the TF divided into quintiles based on ChIP-seq signal strength. (d) Same as in (c) but zoomed in.

**Supplementary Figure 59:**
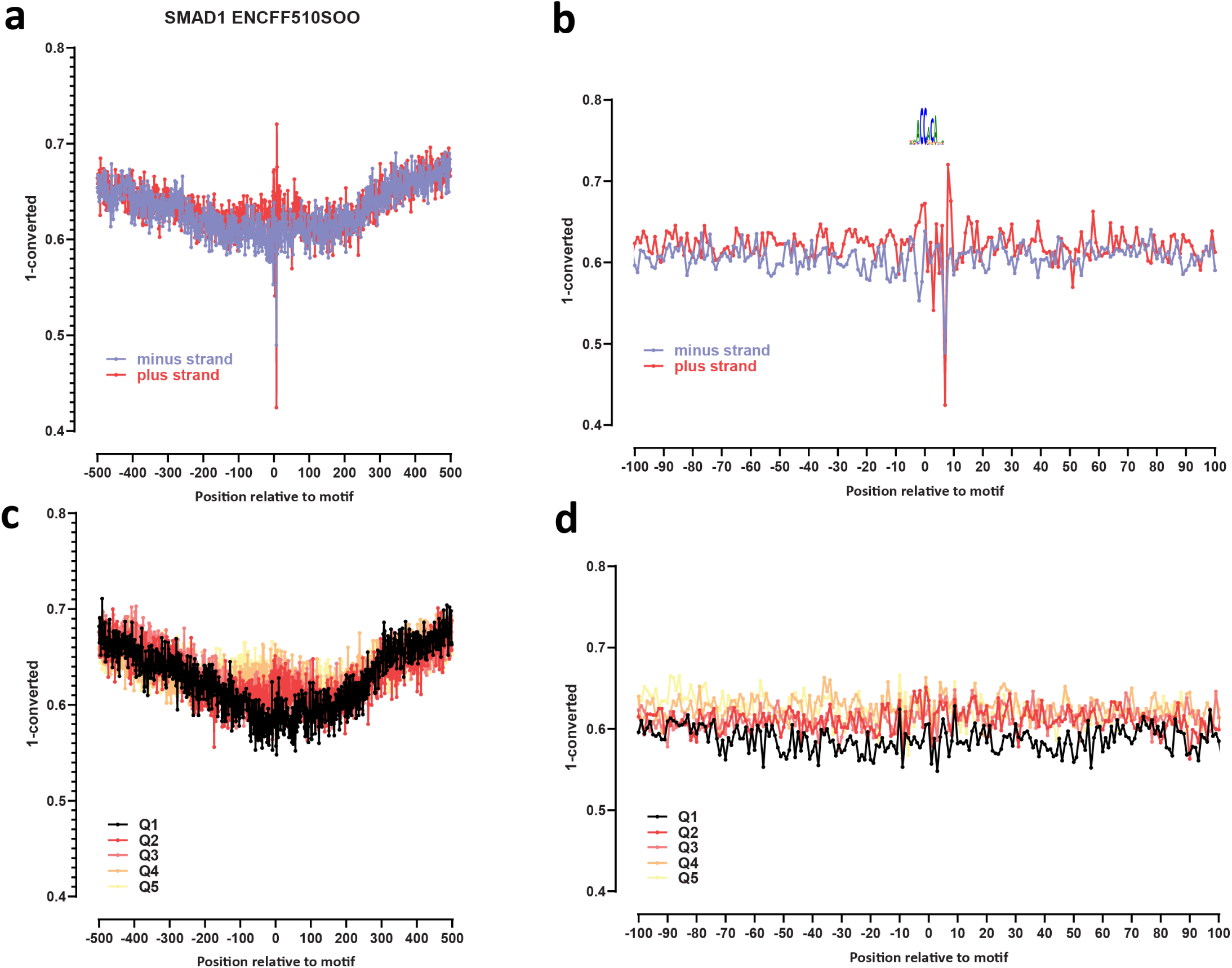
Genome-wide footprint profiles for the SMAD1 transcription factor. GM12878 ssDNA-CseDa01-XL-SMF datasets are shown. (a) Genome-wide footprint profiles over the closest motif to ENCODE ChIP-seq peaks (ENCODE dataset ID indicated on top). Profiles are generated in a strand-aware way with respect to the motif orientation in the genome, and are shown separately for forward and reverse-strand reads. (b) Same as in (a) but zoomed in. The transcription factor sequence recognition motif is shown on top. (c) Genome-wide footprint profiles over the closest motif to ENCODE ChIP-seq peaks for the TF divided into quintiles based on ChIP-seq signal strength. (d) Same as in (c) but zoomed in.

**Supplementary Figure 60:**
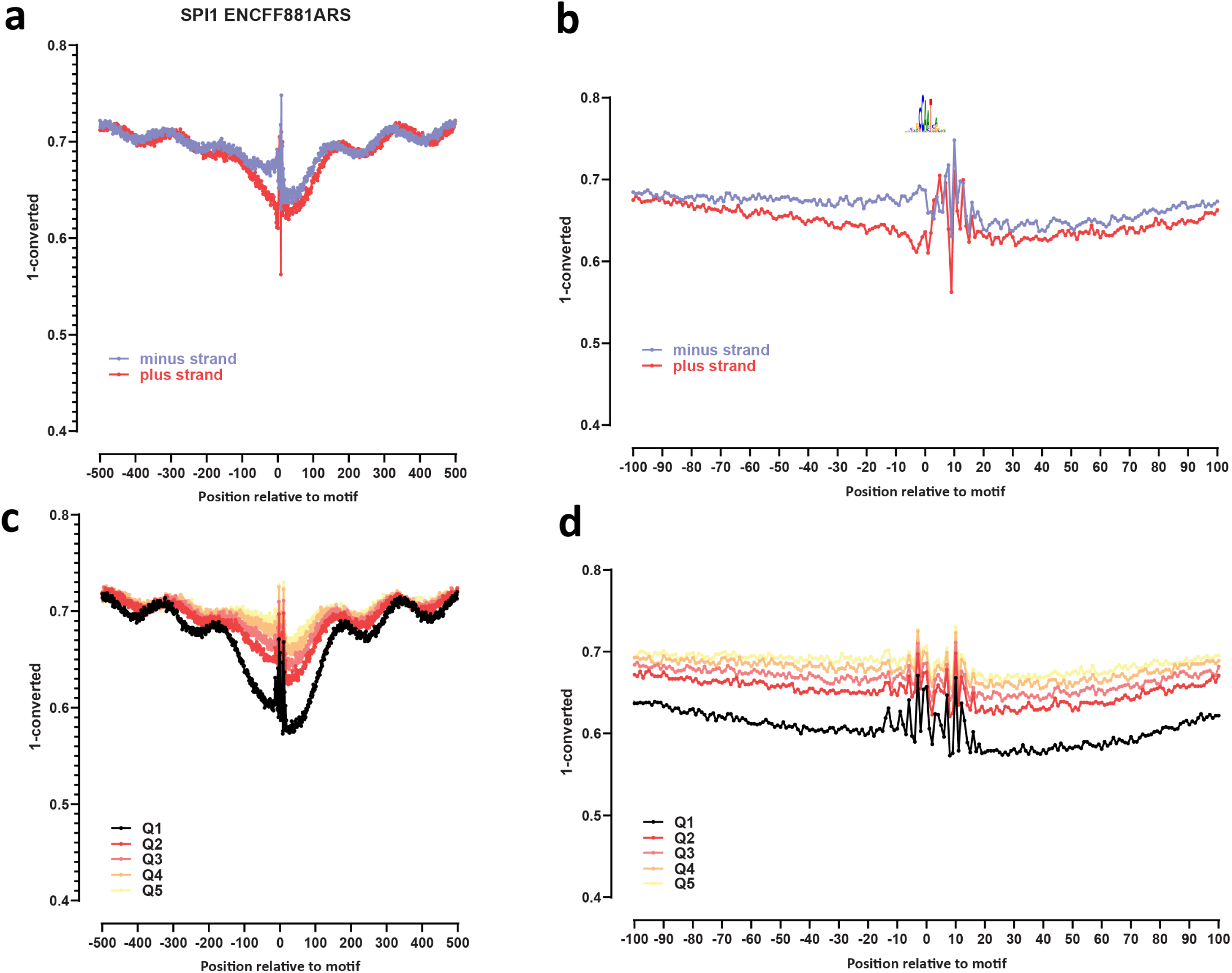
Genome-wide footprint profiles for the SPI1 transcription factor. GM12878 ssDNA-CseDa01-XL-SMF datasets are shown. (a) Genome-wide footprint profiles over the closest motif to ENCODE ChIP-seq peaks (ENCODE dataset ID indicated on top). Profiles are generated in a strand-aware way with respect to the motif orientation in the genome, and are shown separately for forward and reverse-strand reads. (b) Same as in (a) but zoomed in. The transcription factor sequence recognition motif is shown on top. (c) Genome-wide footprint profiles over the closest motif to ENCODE ChIP-seq peaks for the TF divided into quintiles based on ChIP-seq signal strength. (d) Same as in (c) but zoomed in.

**Supplementary Figure 61:**
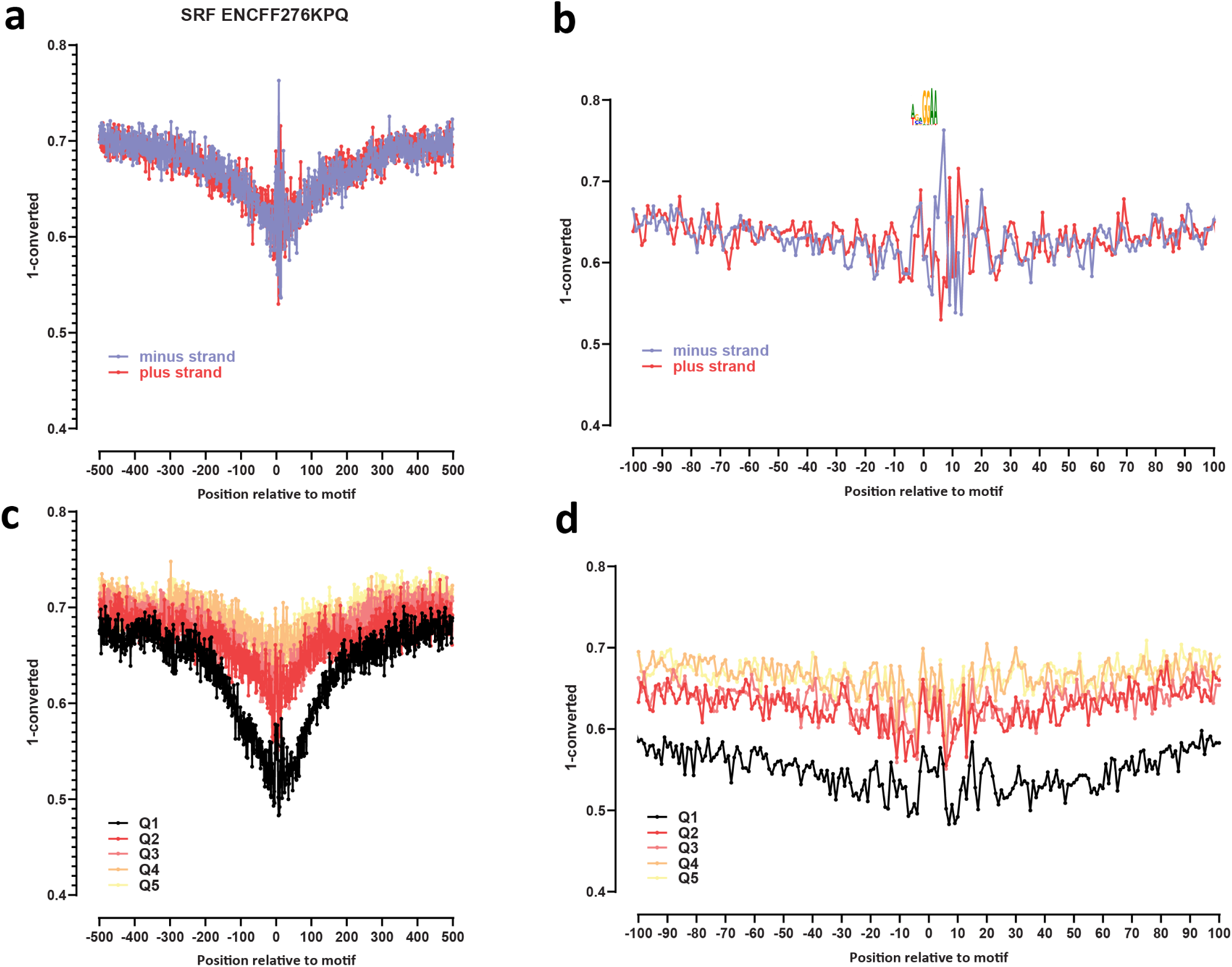
Genome-wide footprint profiles for the SRF transcription factor. GM12878 ssDNA-CseDa01-XL-SMF datasets are shown. (a) Genome-wide footprint profiles over the closest motif to ENCODE ChIP-seq peaks (ENCODE dataset ID indicated on top). Profiles are generated in a strand-aware way with respect to the motif orientation in the genome, and are shown separately for forward and reverse-strand reads. (b) Same as in (a) but zoomed in. The transcription factor sequence recognition motif is shown on top. (c) Genome-wide footprint profiles over the closest motif to ENCODE ChIP-seq peaks for the TF divided into quintiles based on ChIP-seq signal strength. (d) Same as in (c) but zoomed in.

**Supplementary Figure 62:**
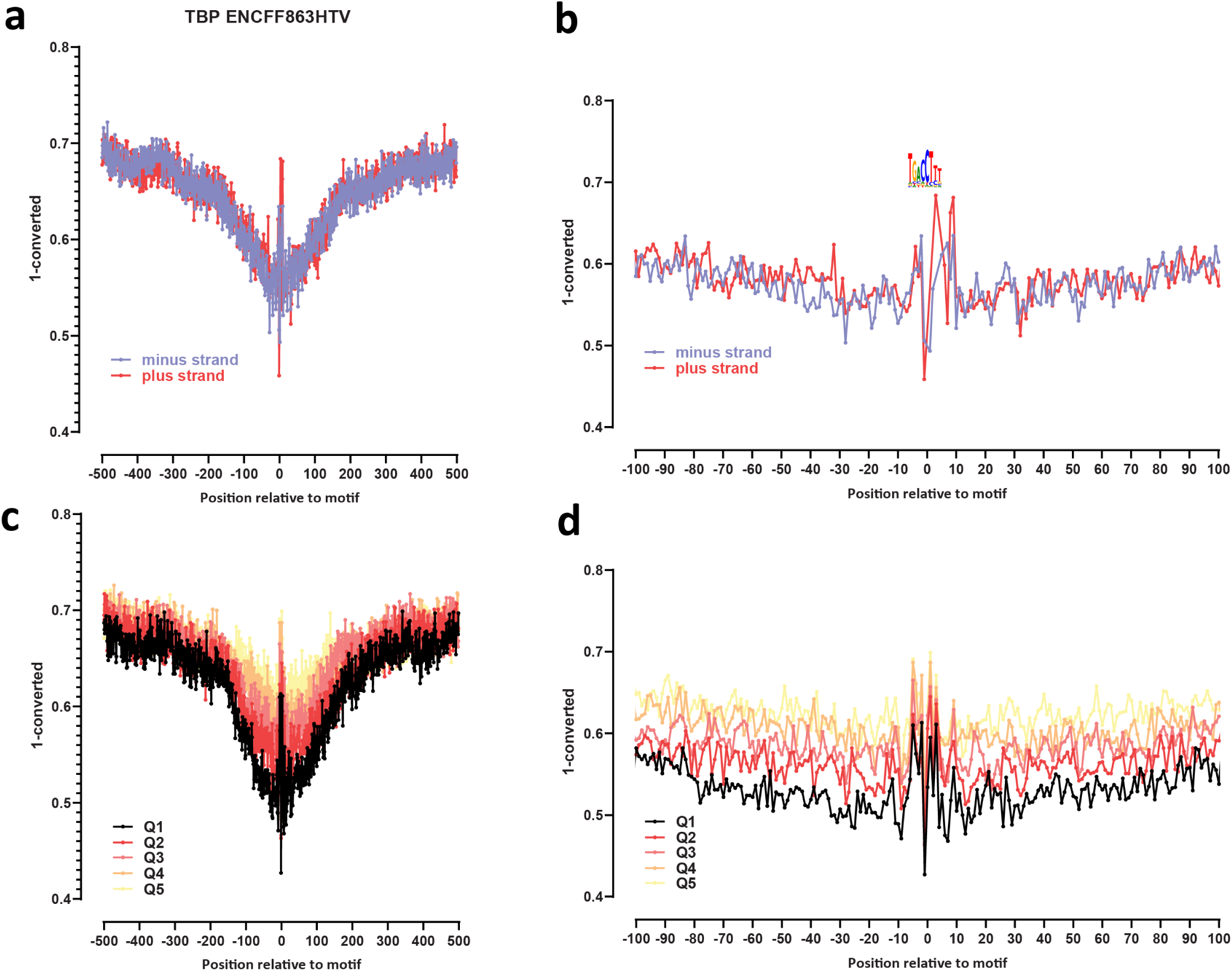
Genome-wide footprint profiles for the TBP transcription factor. GM12878 ssDNA-CseDa01-XL-SMF datasets are shown. (a) Genome-wide footprint profiles over the closest motif to ENCODE ChIP-seq peaks (ENCODE dataset ID indicated on top). Profiles are generated in a strand-aware way with respect to the motif orientation in the genome, and are shown separately for forward and reverse-strand reads. (b) Same as in (a) but zoomed in. The transcription factor sequence recognition motif is shown on top. (c) Genome-wide footprint profiles over the closest motif to ENCODE ChIP-seq peaks for the TF divided into quintiles based on ChIP-seq signal strength. (d) Same as in (c) but zoomed in.

**Supplementary Figure 63:**
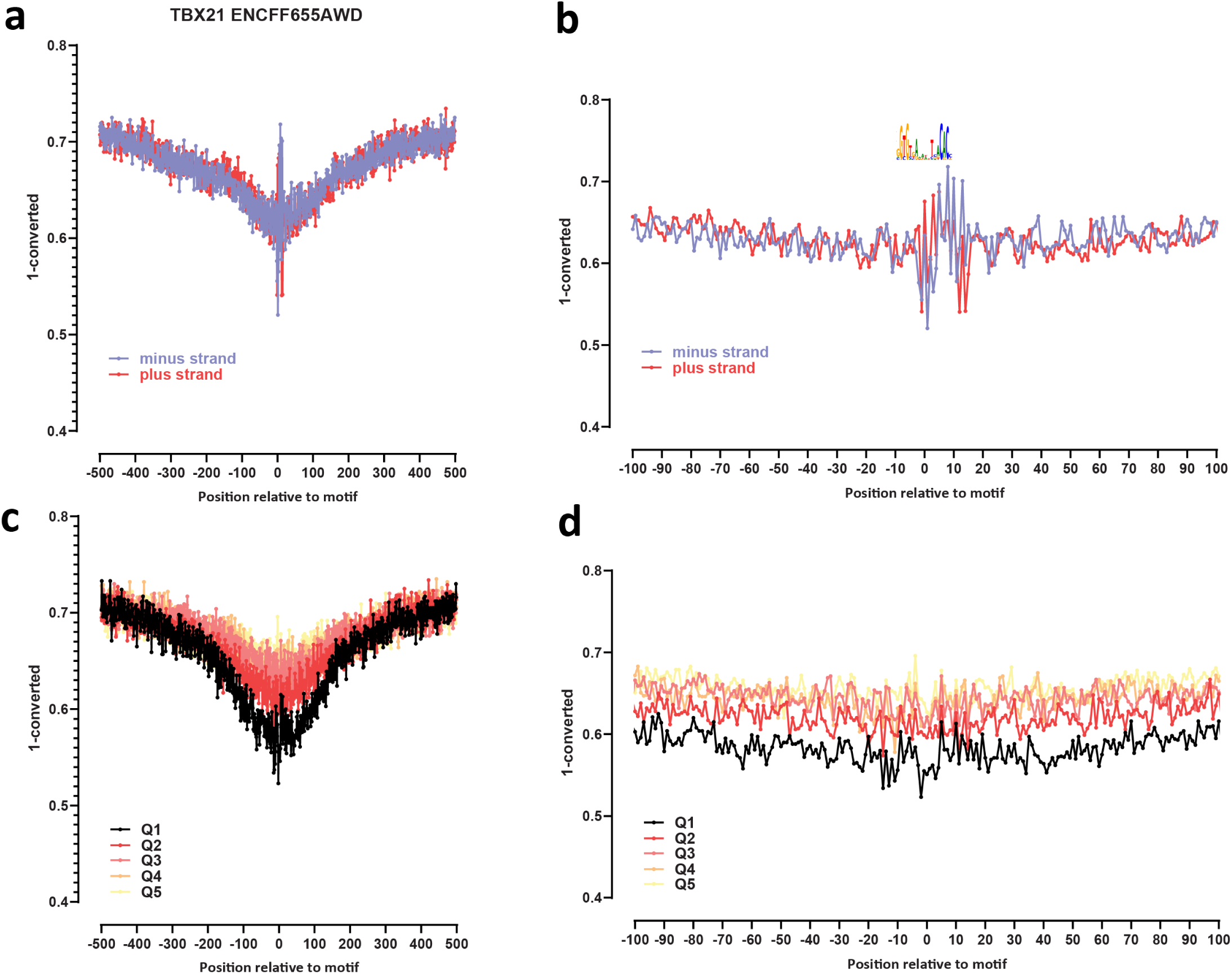
Genome-wide footprint profiles for the TBX21 transcription factor. GM12878 ssDNA-CseDa01-XL-SMF datasets are shown. (a) Genome-wide footprint profiles over the closest motif to ENCODE ChIP-seq peaks (ENCODE dataset ID indicated on top). Profiles are generated in a strand-aware way with respect to the motif orientation in the genome, and are shown separately for forward and reverse-strand reads. (b) Same as in (a) but zoomed in. The transcription factor sequence recognition motif is shown on top. (c) Genome-wide footprint profiles over the closest motif to ENCODE ChIP-seq peaks for the TF divided into quintiles based on ChIP-seq signal strength. (d) Same as in (c) but zoomed in.

**Supplementary Figure 64:**
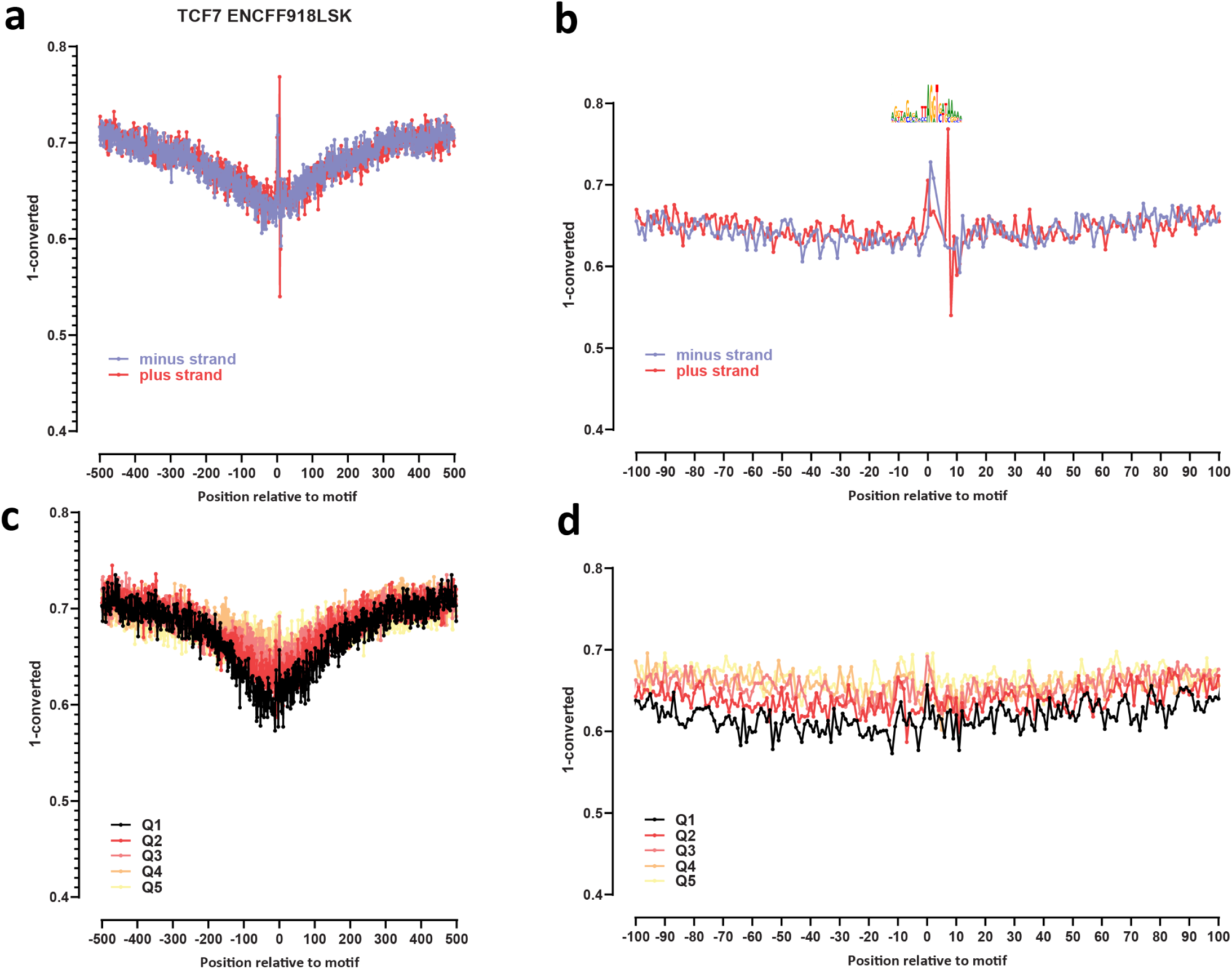
Genome-wide footprint profiles for the TCF7 transcription factor. GM12878 ssDNA-CseDa01-XL-SMF datasets are shown. (a) Genome-wide footprint profiles over the closest motif to ENCODE ChIP-seq peaks (ENCODE dataset ID indicated on top). Profiles are generated in a strand-aware way with respect to the motif orientation in the genome, and are shown separately for forward and reverse-strand reads. (b) Same as in (a) but zoomed in. The transcription factor sequence recognition motif is shown on top. (c) Genome-wide footprint profiles over the closest motif to ENCODE ChIP-seq peaks for the TF divided into quintiles based on ChIP-seq signal strength. (d) Same as in (c) but zoomed in.

**Supplementary Figure 65:**
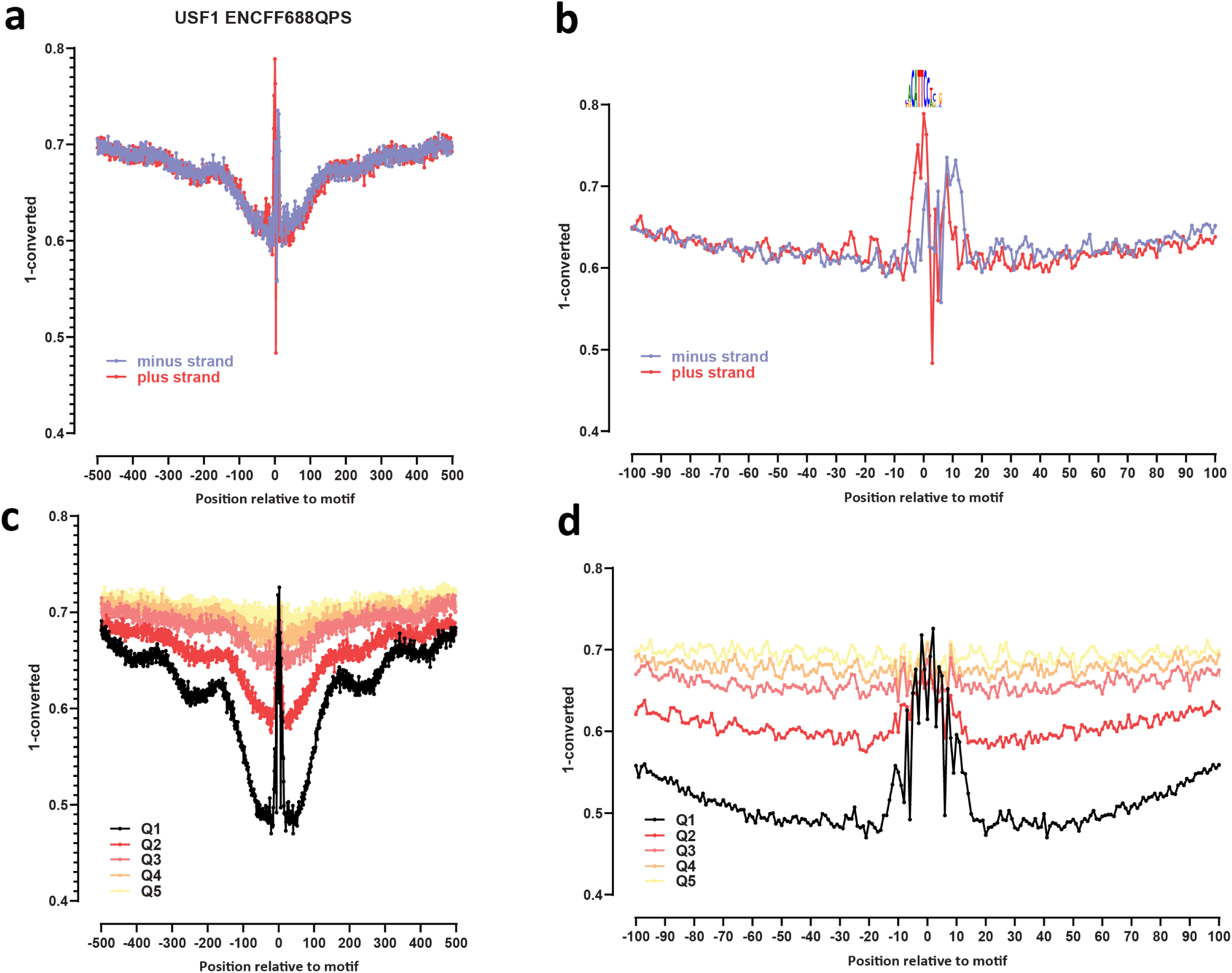
Genome-wide footprint profiles for the USF1 transcription factor. GM12878 ssDNA-CseDa01-XL-SMF datasets are shown. (a) Genome-wide footprint profiles over the closest motif to ENCODE ChIP-seq peaks (ENCODE dataset ID indicated on top). Profiles are generated in a strand-aware way with respect to the motif orientation in the genome, and are shown separately for forward and reverse-strand reads. (b) Same as in (a) but zoomed in. The transcription factor sequence recognition motif is shown on top. (c) Genome-wide footprint profiles over the closest motif to ENCODE ChIP-seq peaks for the TF divided into quintiles based on ChIP-seq signal strength. (d) Same as in (c) but zoomed in.

**Supplementary Figure 66:**
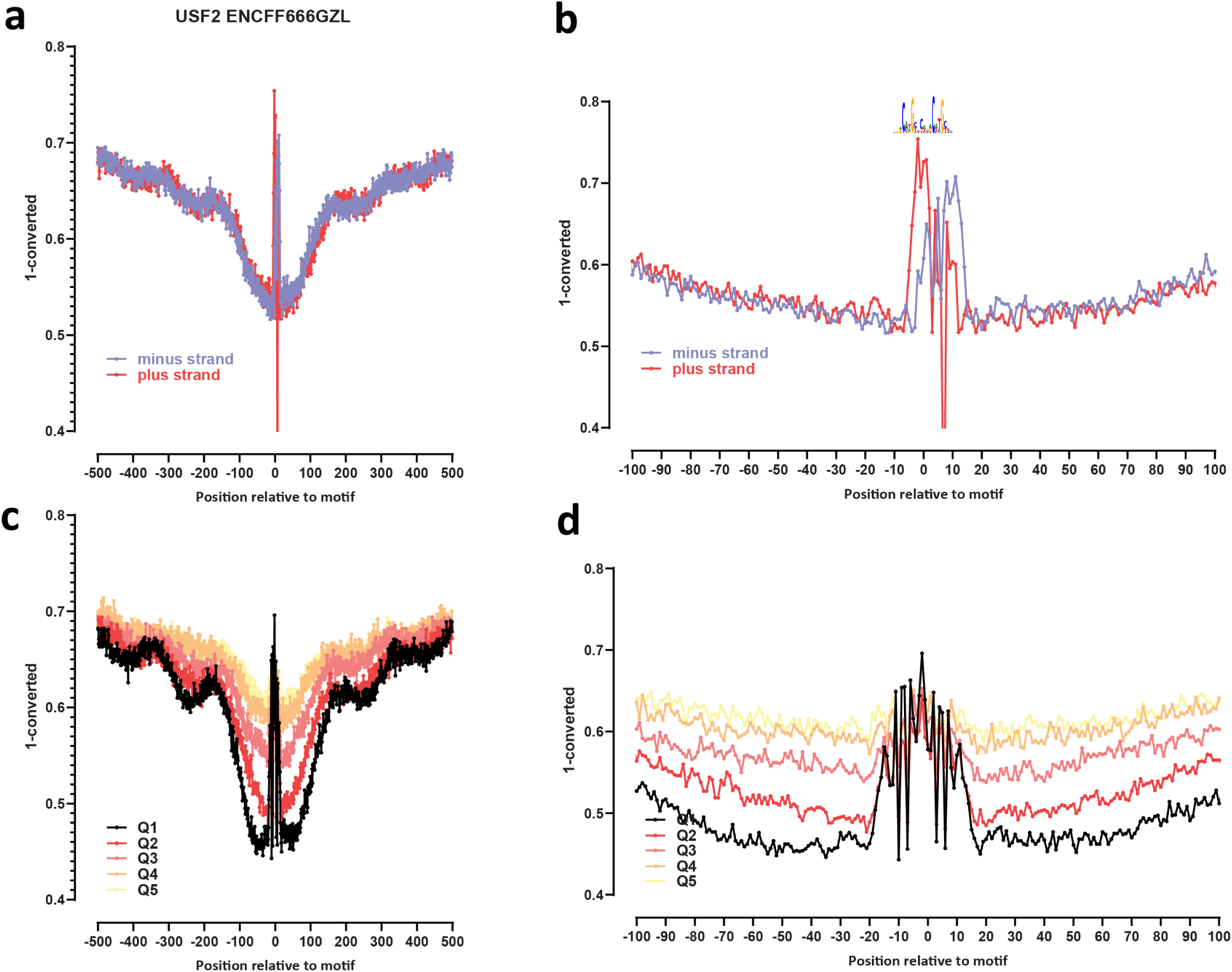
Genome-wide footprint profiles for the USF2 transcription factor. GM12878 ssDNA-CseDa01-XL-SMF datasets are shown. (a) Genome-wide footprint profiles over the closest motif to ENCODE ChIP-seq peaks (ENCODE dataset ID indicated on top). Profiles are generated in a strand-aware way with respect to the motif orientation in the genome, and are shown separately for forward and reverse-strand reads. (b) Same as in (a) but zoomed in. The transcription factor sequence recognition motif is shown on top. (c) Genome-wide footprint profiles over the closest motif to ENCODE ChIP-seq peaks for the TF divided into quintiles based on ChIP-seq signal strength. (d) Same as in (c) but zoomed in.

**Supplementary Figure 67:**
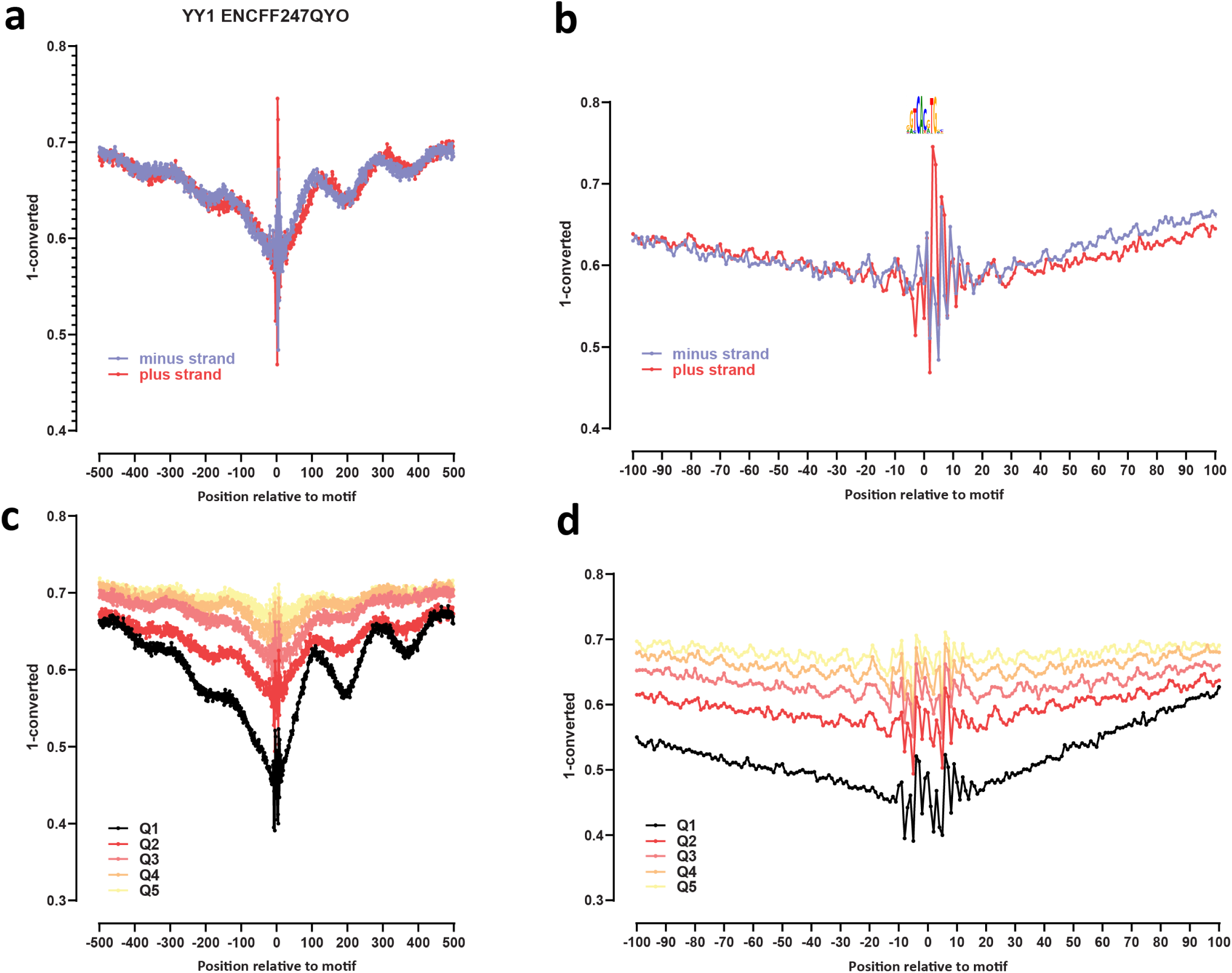
Genome-wide footprint profiles for the YY1 transcription factor. GM12878 ssDNA-CseDa01-XL-SMF datasets are shown. (a) Genome-wide footprint profiles over the closest motif to ENCODE ChIP-seq peaks (ENCODE dataset ID indicated on top). Profiles are generated in a strand-aware way with respect to the motif orientation in the genome, and are shown separately for forward and reverse-strand reads. (b) Same as in (a) but zoomed in. The transcription factor sequence recognition motif is shown on top. (c) Genome-wide footprint profiles over the closest motif to ENCODE ChIP-seq peaks for the TF divided into quintiles based on ChIP-seq signal strength. (d) Same as in (c) but zoomed in.

**Supplementary Figure 68:**
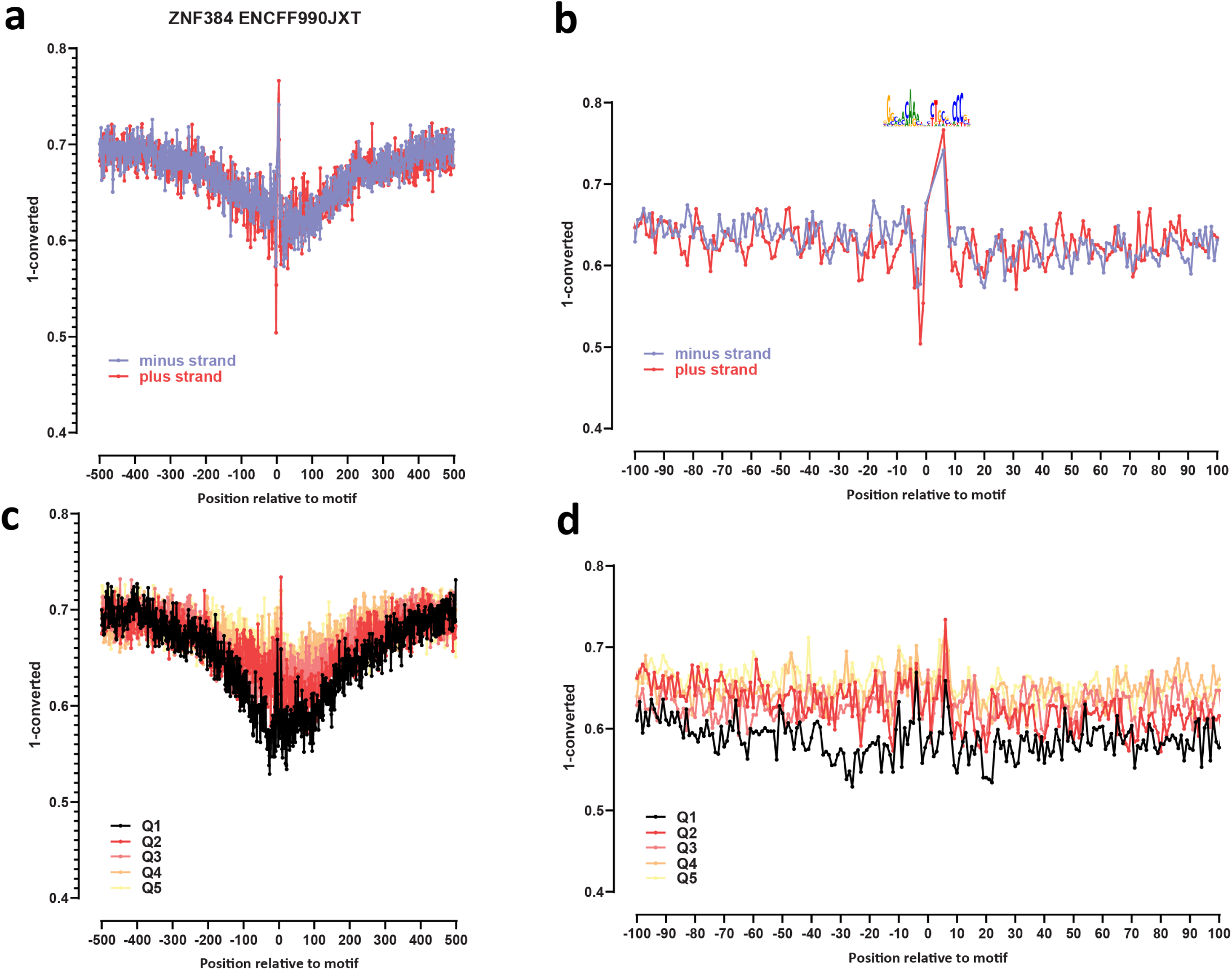
Genome-wide footprint profiles for the ZNF384 transcription factor. GM12878 ssDNA-CseDa01-XL-SMF datasets are shown. (a) Genome-wide footprint profiles over the closest motif to ENCODE ChIP-seq peaks (ENCODE dataset ID indicated on top). Profiles are generated in a strand-aware way with respect to the motif orientation in the genome, and are shown separately for forward and reverse-strand reads. (b) Same as in (a) but zoomed in. The transcription factor sequence recognition motif is shown on top. (c) Genome-wide footprint profiles over the closest motif to ENCODE ChIP-seq peaks for the TF divided into quintiles based on ChIP-seq signal strength. (d) Same as in (c) but zoomed in.

**Supplementary Figure 69:**
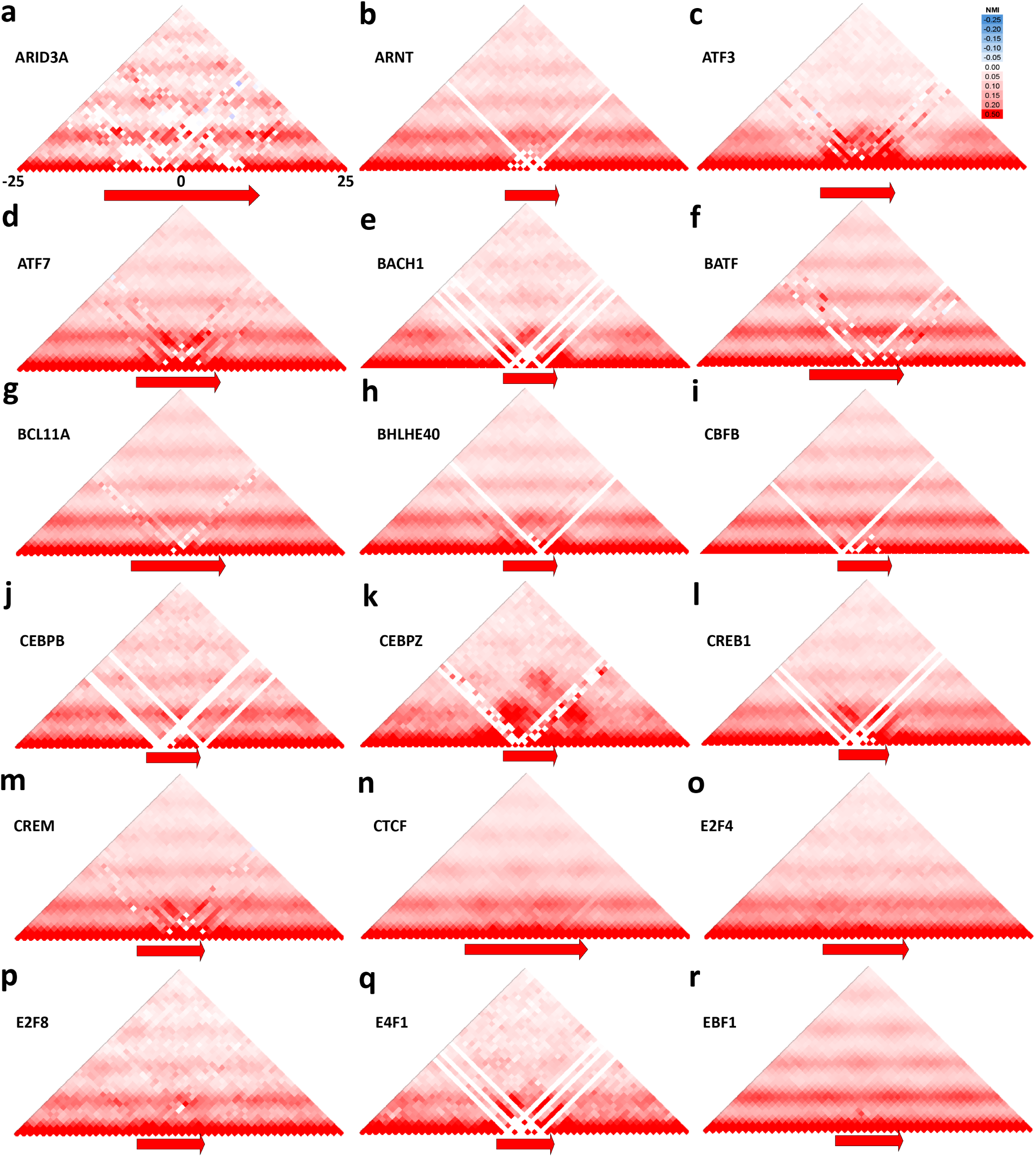
Average co-accessibility profiles around occupied TF recognition motifs. Shown are average NMI values for each position in a 50-bp window around the TF recognition motif for the indicated TFs. Only the nearest motifs within ±100bp of ENCODE ChIP-seq peaks were used as input.

**Supplementary Figure 70:**
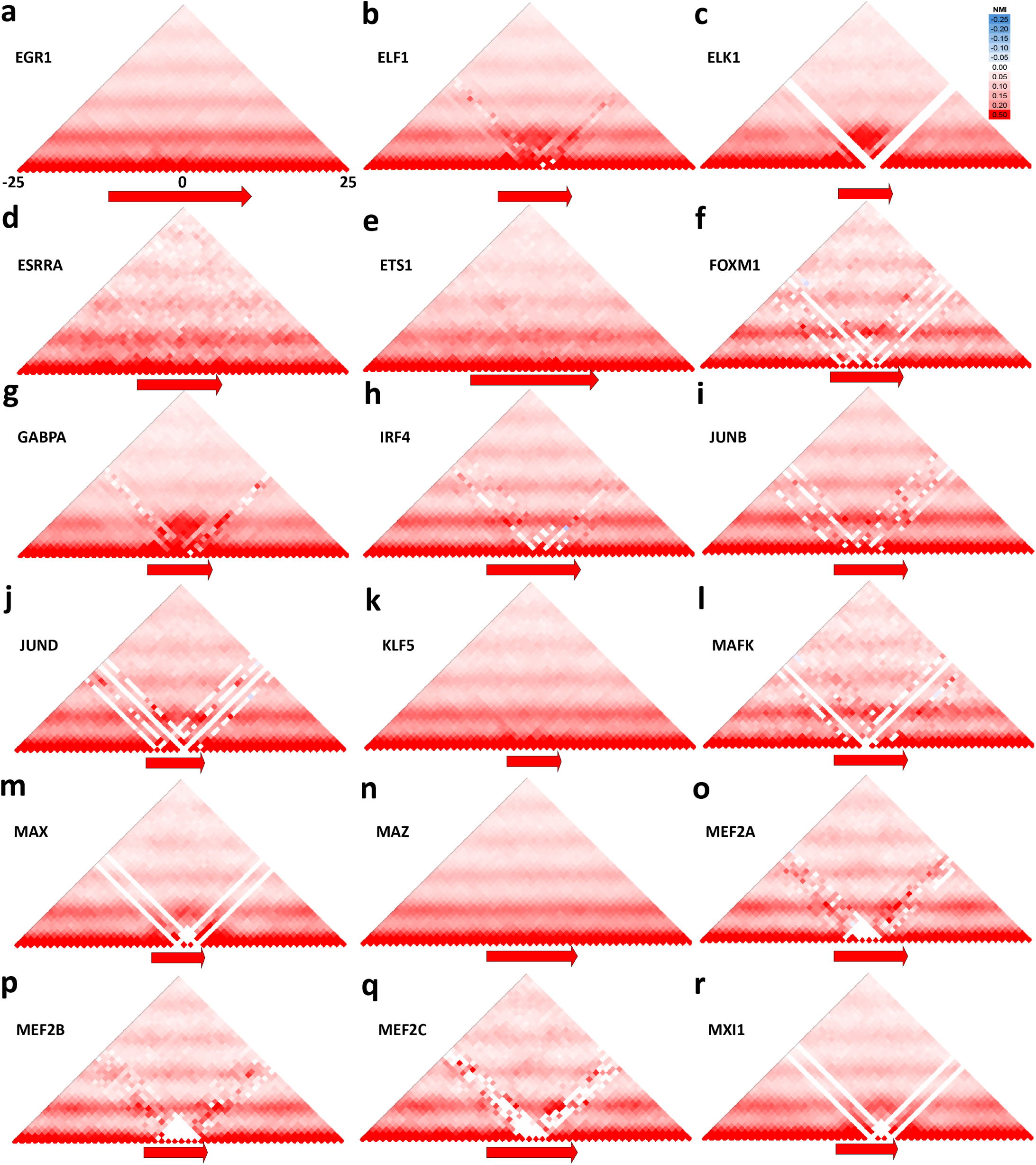
Average co-accessibility profiles around occupied TF recognition motifs. Shown are average NMI values for each position in a 50-bp window around the TF recognition motif for the indicated TFs. Only the nearest motifs within ±100bp of ENCODE ChIP-seq peaks were used as input.

**Supplementary Figure 71:**
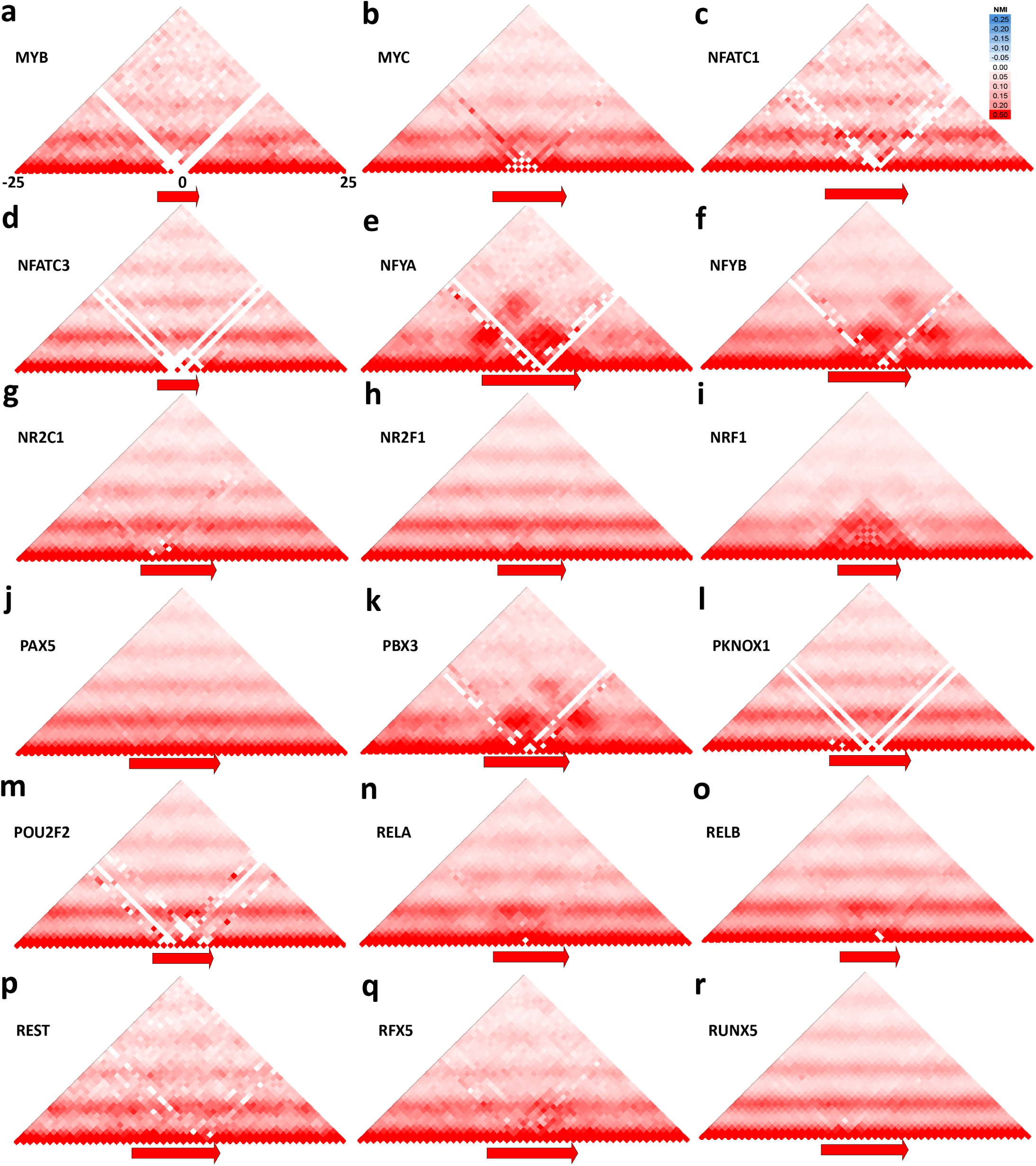
Average co-accessibility profiles around occupied TF recognition motifs. Shown are average NMI values for each position in a 50-bp window around the TF recognition motif for the indicated TFs. Only the nearest motifs within ±100bp of ENCODE ChIP-seq peaks were used as input.

**Supplementary Figure 72:**
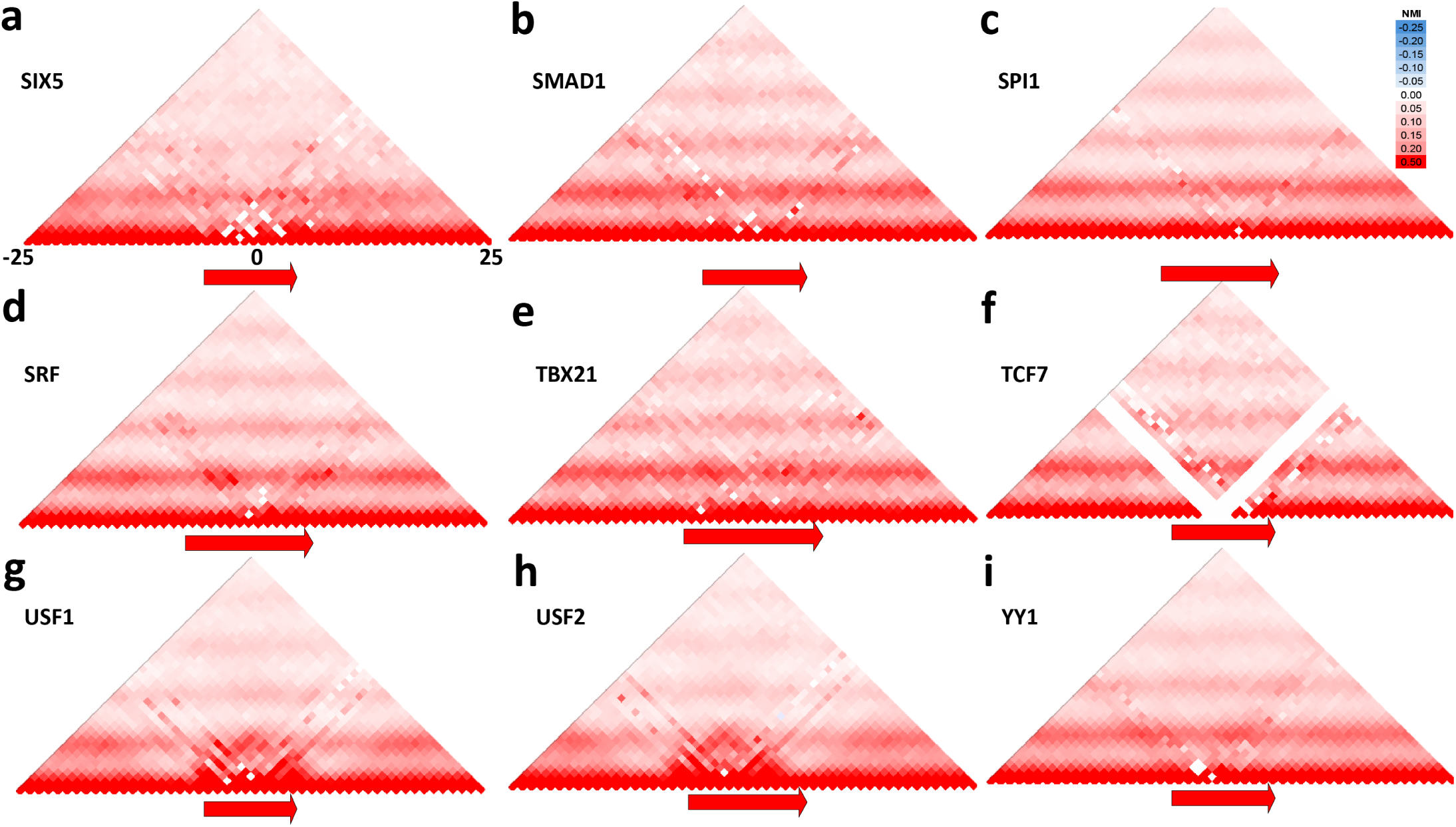
Average co-accessibility profiles around occupied TF recognition motifs. Shown are average NMI values for each position in a 50-bp window around the TF recognition motif for the indicated TFs. Only the nearest motifs within ±100bp of ENCODE ChIP-seq peaks were used as input.

